# Reconstructing EBV reactivation and DNA damage response kinetics in morphologic pseudotime

**DOI:** 10.1101/2025.10.06.680675

**Authors:** Dina G. Tekle, Jonathan Z. Sexton, Elliott D. SoRelle

## Abstract

Epstein-Barr virus (EBV) lytic infection contributes to virus-associated cancers and autoimmunity and depends on viral subversion of host DNA damage responses (DDR). Here, we use high-content screening (HCS) for morphologic and pseudotemporal profiling of single-cell EBV reactivation and DDR dynamics spanning seven lytic induction treatments in nine B cell lymphoma models. We generated an atlas (>750,000 cells) of spatiotemporally distinct phenotypes of immediate-early (IE) and late lytic proteins, viral and cellular DNA replication, and double-stranded break (DSB) DDR factors. Single-cell segmentation, feature extraction, and clustering identify treatment- and model-dependent cell responses and lytic induction. Notably, genotoxin-induced DDR profiles differ in lytic versus latent cells, and lytic protein localization varies across pharmacologic and physiologic stimuli. Pseudotime trajectories of physiologic reactivation reveal viral replication compartment (VRC) nucleation and expansion alongside concomitant host DDR localization. The early DDR marker γH2AX is depleted from VRCs but widespread across host chromatin throughout reactivation. Surprisingly, the lytic-essential late DDR protein 53BP1 is present in lytic cells prior to viral genome replication but subsequently absent from VRCs and host chromatin, indicating spatial and kinetic DDR dysregulation during EBV reactivation. Collectively, these data support a model wherein EBV transiently employs host DSB DDR mediators to initiate genome replication while a host-targeted DDR is initiated but impaired. We further demonstrate biological generalizability and utility of the method across confocal and epifluorescence systems. Thus, this work demonstrates a powerful technique to recover host-virus dynamics from static timepoints and will support future high-throughput single-cell virology applications.

## Introduction

Epstein-Barr virus (EBV) is an important human pathogen due to its ubiquity (globally, over 95% of adults are EBV-seropositive) (1), involvement in numerous cancers (2), and several autoimmune diseases (3). This double-stranded DNA gammaherpesvirus infects human B lymphocytes (1), establishes latency in the memory B cell reservoir for the life of the host (4), and undergoes periodic reactivation to lytic infection (5). Latent and lytic EBV infection each depend on coordinated viral transcription programs that modulate host cell biology, evade innate and adaptive immune control, and contribute to disease through distinct mechanisms (6). Single-cell sequencing studies have demonstrated considerable heterogeneity related to EBV infection phase and strain (7–10), functions of individual viral genes (11), and immune responses to infection in patient samples (12, 13). Such studies provide genomic perspectives of host-virus interactions, however complementary single-cell methods are greatly needed to study other molecular aspects, rare phenotypes, and infected cell dynamics at scale.

The EBV lytic cycle is essential for host-to-host transmission. However, it is well known that lytic infection is challenging to study due to its asynchrony and relative rarity in B cells (14). Thus, we developed a high-content screening (HCS) approach for systematic study of EBV reactivation across B cell lymphoma models induced by common experimental treatments, physiologic stimuli, and clinically relevant drugs. Lytic EBV infection can be activated by diverse stimuli via distinct mechanisms. Differentiation of infected B cells into plasma cells induces reactivation through host transcription factor activation of immediate-early (IE) gene promoters and other mechanisms (15–17). B cell receptor (BCR) signaling activated by receptor crosslinking (18) and SMAD-mediated activation stimulated by TGF-β (19–21) are additional physiologic triggers of lytic infection. Common laboratory methods to induce lytic expression include treatment with phorbol esters (22, 23), butyrate (24), and/or reactive oxygen species (25). Clinically relevant chemotherapeutic agents used to treat B cell lymphomas also induce lytic expression in some B cell models (26, 27), though precise mechanisms are unclear. These stimuli can activate the essential IE proteins, Zta (a.k.a. Z, ZEBRA; encoded by *BZLF1*) and Rta (R; encoded by *BRLF1*), which activate lytic gene expression from intranuclear viral episomes (28). Zta binds Z response elements (ZREs) (29) in epigenetically repressed viral lytic gene promoters including the origin of lytic replication (*oriLyt*), early lytic genes essential for viral DNA replication and gene expression (30), and *BZLF1* itself (31). Intriguingly, Zta is an AP-1 family pioneering transcription factor that mediates epigenetic de-repression of viral and host genes via nucleosome eviction (32–35). Rta indirectly activates *BZLF1* expression, additional early-kinetic lytic genes (36), and itself (37) to enable successful lytic infection. While IE and early genes are expressed prior to viral DNA amplification via rolling circle replication, activation of late-kinetic genes that encode numerous tegument, capsid, and envelope glycoproteins depends on viral genome synthesis (5).

Despite distinct triggers, lytic infection by EBV (and numerous other viruses) requires activation and modulation of host DNA damage response (DDR) pathways (38). Specifically, mediators of the double-stranded break (DSB) repair arm of the DDR are important for EBV lytic infection. These include phosphorylated ataxia telangiectasia mutated (ATM) kinase (39), phospho-Ser139 in histone variant H2A.X (γH2AX) (32), the homologous recombination (HR) repair complex MRE11-RAD51-NBS1 (MRN) (40), and the non-homologous end joining (NHEJ) mediator 53BP1 (41). Extensive prior studies (40, 42, 43) have identified these and other host DNA repair proteins within intranuclear viral replication compartments (VRCs), which exhibit distinct morphologies and effects on nuclear organization before and after viral genome replication (44–47). Prior imaging of DDR proteins during lytic infection have used a variety of model systems, several of which are non-physiologic cell types for EBV infection. In some cases, imaging studies have yielded seemingly inconsistent results for DDR protein localization during reactivation. For example, vector-based expression of Zta (48), BKRF4 tegument protein (49), or the viral DNA polymerase processivity factor EA-D (*BMRF1*) (50) impair formation of 53BP1 in epithelial cell lines via distinct protein-protein interactions, yielding diffuse nuclear patterns. By contrast, 53BP1 focus formation and co-localization with BMRF1 in apparent VRCs are distinct phenotypes observed in Zta-transfected 293 cells harboring the EBV genome as a bacterial artificial chromosome (BAC) (43). These disparate phenotypes may indicate complex roles in viral reactivation. Like 53BP1, the early DDR biomarker γH2AX plays an important role in lytic infection based on biochemical studies (32, 51, 52). However direct visualization of γH2AX in cells expressing EBV lytic genes has been notably limited and restricted to epithelial models transfected with exogenous Zta overexpression vectors (42, 48). Critically, the disparate presence and nuclear localization of DNA damage mediators may be predisposed not only by model differences but also by technical limitations including low sample throughput and a lack of temporal resolution to account for the dynamic nature of lytic infection and concomitant DDR. Paired with the inherent challenge posed by infection heterogeneity, these factors underscore a critical need for systematic quantitative single-cell methods.

To address this need, we developed a high-content screening (HCS) assay with morphologic pseudotime analyses based on >800 automatically quantified cell signal features. This approach leverages the asynchronous nature of infected cell populations to reconstruct high-resolution phenotype transitions from snapshot sampling, thereby providing a powerful technique to dissect host-virus dynamics at single-cell resolution. We show the utility of this method by applying it to resolve previously unrecognized DDR kinetics during EBV lytic infection in B cells. Our data show that lytic gene expression and DNA damage profiles vary across induction methods. Cells that undergo lytic reactivation upon treatment with clinically relevant genotoxins exhibit fundamentally distinct DNA damage patterns relative to latently infected cells in the same treatments. Our data demonstrate that γH2AX is rapidly depleted from VRCs – even prior to viral DNA replication – but persists across cellular chromatin throughout reactivation. The late DDR factor 53BP1 is present prior to EBV DNA replication but is lost subsequently from VRCs. Whereas γH2AX remains detectable in host chromatin, the lack of 53BP1 foci indicates incomplete host-targeted DDR throughout lytic infection. Thus, DNA damage responses are spatiotemporally dysregulated during physiologic EBV lytic reactivation.

We further demonstrate the versatility of HCS and pseudotime analyses to study infected cell biology as well as the method’s technical compatibility across confocal and epifluorescence microscopes. Accordingly, we expect our approach will provide a powerful tool for high-throughput single-cell virology. The method’s general utility should be extensible to studies of other intracellular pathogens and cellular dynamics for which live-cell imaging is challenging or technically intractable.

## Results

### Systematic single-cell analysis of lytic infection across B cell models and induction methods

EBV lytic infection can be triggered by diverse treatments as measured via mRNA and protein expression, yet individual cell responses to these treatments are incompletely understood. Thus, we developed plate-based methods to systematically investigate B cell lymphoma model responses to biologic stimuli and genotoxic drugs at scale with HCS and single-cell analyses (**Figure 1A**). Following treatment in 96-well culture plates, cells were subjected to live-cell labeling (typically, a 1-hour DNA synthesis pulse with 5-ethynyl-2’-deoxyuridine (EdU) to label VRCs or a mitochondria-selective dye). Labeled cells were washed by centrifugation, transferred to imaging 96-well plates coated with poly-D-lysine (PDL), centrifuged to enhance cell deposition, and fixed to promote cell retention through subsequent permeabilization and fluorescence detection steps. Stained cell plates were imaged using automated confocal or epifluorescence systems, and images were processed for high-throughput single-cell analyses as described below and detailed in the Experimental Methods section.

**Figure 1.**
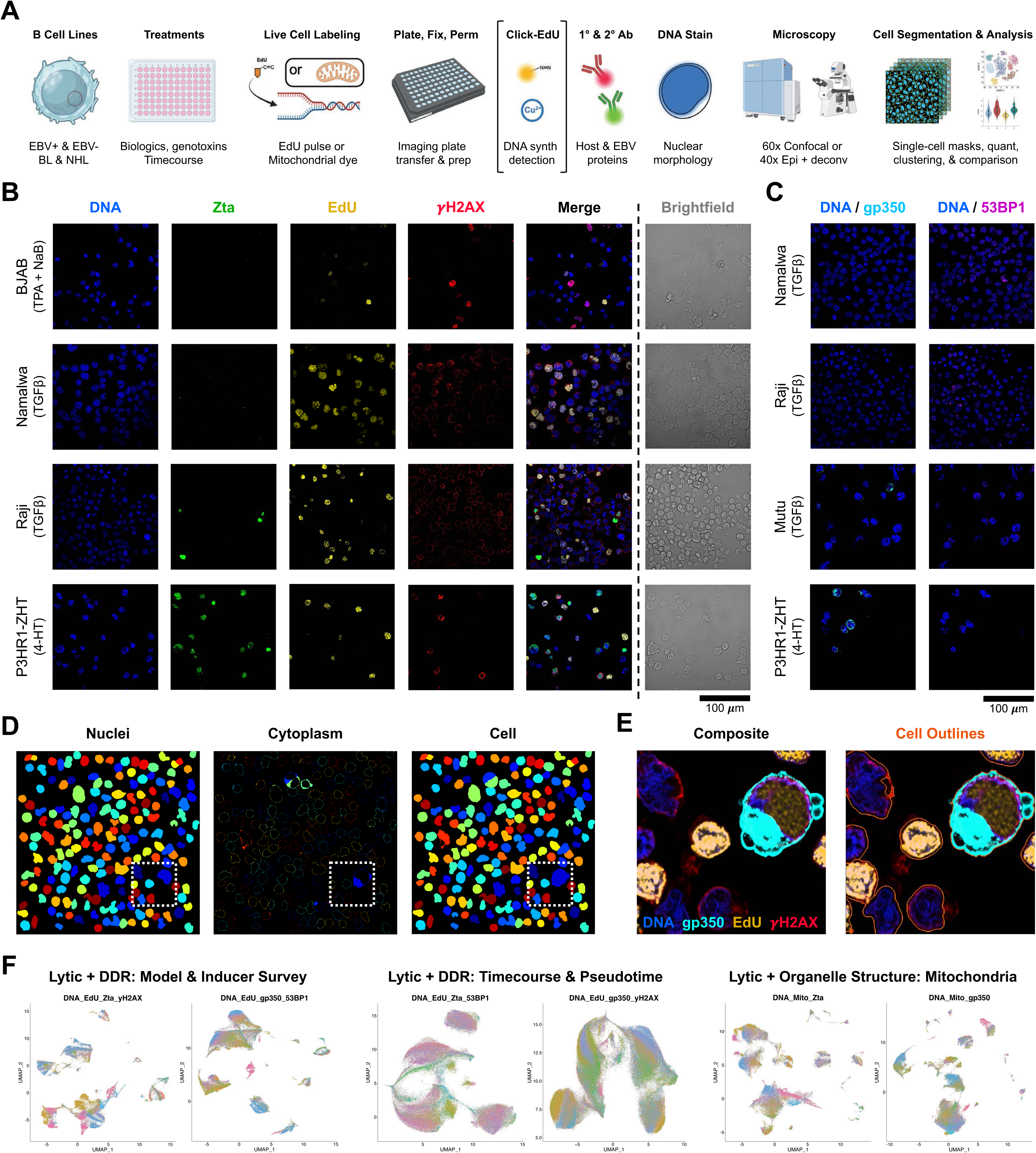
High-content screening assay and study design. **A**) Experimental workflow overview. B cell lymphoma cell lines were treated with biologic and pharmacologic inducers of the EBV lytic cycle then assayed for host and viral processes and proteins. Live cell labeling steps (e.g., pulsed EdU incorporation, mitochondria-selective dyes) were performed prior to harvest, transfer, and fixation on coated 96-well imaging plates. For experiments examining host and viral DNA synthesis, incorporated EdU was detected via copper-catalyzed click chemistry (bracketed step). Host and viral proteins were detected via immunofluorescence using primary and secondary antibodies, after which nuclei were stained, and plates were imaged on automated confocal or epifluorescence microscopes. Individual cells were segmented from acquired images and subsequently analyzed. **B**) Representative images of nuclei (DNA), Zta (BZLF1, immediate early (IE) EBV lytic gene), pulse-labeled DNA synthesis (EdU), phospho-(Ser139) histone H2A.X (γH2AX, early DNA damage repair biomarker), and brightfield for select cell lines and treatments. BJAB (top row) is an EBV-negative Burkitt Lymphoma (BL) cell line; Namalwa is an EBV-positive BL line defective for lytic reactivation due to viral genome integration within host chromosome 1; Raji is an EBV-positive BL line in which IE and early (E) lytic genes are functional but a mutation in the viral gene *BALF2* impairs EBV genome replication and progression to the late lytic phase; P3HR1-ZHT is an EBV-positive BL line engineered for inducible lytic infection upon treatment with 4-hydroxytamoxifen). **C**) Representative images of nuclei, gp530 (BLLF1, late EBV lytic gene), and 53BP1 (late biomarker of DNA damage repair via non-homologous end joining (NHEJ)) in Namalwa, Raji, Mutu (EBV-positive lytic-competent BL cell line), and P3HR1-ZHT cells. **D**) Representative CellProfiler segmentation of nuclei, cytoplasm, and whole cells based on fluorescence signals. **E**) Detail of composite fluorescence signals used for whole-cell segmentation (region denoted by dashed outline in D). Composite signal includes nuclei (blue), gp350 (cyan), DNA synthesis (yellow), and γH2AX (red). **F**) Summary of confocal imaging experiments performed in this study. Each UMAP depicts segmented cells detected from a given stain panel. Two survey experiments (DNA_EdU_Zta_γH2AX and DNA_EdU_gp350_53BP1) were performed to evaluate eight B cell lines across five induction methods as well as biological and technical controls. Two timecourse experiments (DNA_EdU_Zta_53BP1 and DNA_EdU_gp350_γH2AX) were performed in four lines across three induction methods and controls for cells 24-, 48-, and 72-hours post-treatment. Two experiments were performed to investigate mitochondrial localization and morphology during EBV lytic infection (DNA_Mito_Zta and DNA_Mito_gp350). Cells are color-coded by well position on 96-well imaging plates.

Nine B cell lymphoma lines were utilized in the present study (**Table S1**). These included the EBV-negative Burkitt Lymphoma (BL) BJAB cell line; two EBV-positive BL lines with lytic cycle defects (Namalwa and Raji, see Experimental Methods for details); four EBV-positive BL lines competent for complete lytic infection (Daudi, Mutu, Jijoye, and P3HR1-ZHT, which was engineered from a subclone of Jijoye for hydroxytamoxifen-inducible lytic expression); and two EBV-positive non-Hodgkin lymphomas (NHL) with immunoblastic phenotypes (Farage, IBL1). BJAB cells provided a negative control for EBV protein immunofluorescence. Namalwa cells provided an EBV-positive cell line incapable of lytic reactivation as negative control. Raji cells exhibit expression of the EBV immediate early (IE) transactivator Zta but fail to progress to late lytic infection (indicated by gp350 immunofluorescence) due to an inactivating mutation of the early lytic gene *BALF2*, which encodes a lytic-essential single-stranded DNA binding protein (47). P3HR1-ZHT cells treated with 4-hydroxytamoxifen (4HT) served as both a positive control for lytic infection across models and a robust experimental condition yielding relatively high frequencies of lytic cells. (**Figure 1B-C**). Additional technical, biological, and treatment control conditions were prepared and quantified to confirm fluorescence signal specificity and negligible inter-channel cross-talk (**Figure S1**). In total, seven different lytic-inducing treatments encompassing physiologic stimuli (TGF-β, anti-IgG), common experimental conditions (phorbol ester (TPA) + sodium butyrate (NaB), H_2_O_2_), and clinically relevant drugs (doxorubicin, etoposide, panobinostat) were assayed in addition to unstimulated controls and treatment with 4HT (P3HR1-ZHT only).

Multichannel images from all tested cell lines and treatment conditions were analyzed with a CellProfiler pipeline (53) for segmentation and feature extraction for regions of interest (ROIs): nuclei, cytoplasm, and the whole cell (**Figure 1D-E**). Nuclei were segmented from the Hoechst 33342 channel, whole-cell masks were generated from an all-channel composite image, and cytoplasm by subtracting nuclear masks from whole-cell masks. Cells were filtered by size to minimize artifactual detection. Per-cell feature extraction was performed for each ROI to quantify more than 800 measurements including signal intensities, shape, texture, and colocalization between fluorescence channels.

Cell measurements were annotated with experiment metadata and standardized pre-processing was performed with PyCytominer (mad_robustize) (54), UMAP dimensionality reduction, and unsupervised Leiden clustering. The resulting datasets were analyzed by adapting functionality from Seurat v5 (55) to support phenotype identification and quantitative comparisons. In total, over 750,000 cells were segmented and quantified across ~12,000 fields of view from six separate fluorescent staining experiments (**Figures 1F, S2, Table S2**). Two survey experiments were conducted across eight cell lines to investigate lytic infection and corresponding DDR via early and late kinetic biomarkers (DNA_EdU_Zta_γH2AX and DNA_EdU_gp350_53BP1, respectively) at a single timepoint (36 h) for six conditions. Two timecourse experiments examined lytic and DDR responses in four cell lines (Namalwa, Mutu, P3HR1-ZHT, IBL1) in four conditions (unstimulated, TGF-β, TPA+NaB, H_2_O_2_) at three timepoints (24, 48, and 72 h) using swapped immunofluorescence combinations to examine late lytic expression alongside an early DDR indicator (DNA_EdU_gp350_γH2AX) and vice versa (DNA_EdU_Zta_53BP1). Additional experiments were performed to examine mitochondrial structure and localization during EBV lytic infection stages assayed via the IE protein Zta (DNA_Mito_Zta) or the late glycoprotein gp350 (DNA_Mito_gp350). Collectively, this approach enabled us to systematically quantify response state heterogeneity in commonly used EBV lytic infection models and spatiotemporally resolve key host and viral processes in individual reactivating cells.

### Differential lytic response phenotypes in B cell lines resolved by induction treatment

EBV predominantly adopts latent infection in B cells, and many lytic induction treatments are inefficient at latency reversal. HCS addresses this challenge by resolving individual cell response states and enabling identification of lytic phenotypes in a high-throughput manner. As an initial demonstration, we quantified and characterized cells stained for nuclei, pulsed DNA synthesis with EdU, Zta, and γH2AX in a survey of B cell lines and induction treatments (DNA_EdU_Zta_γH2AX). Twenty-one segmented cell phenotypes were identified from this experiment following uniform manifold approximation projection (UMAP) dimensional reduction (56) and Leiden clustering (57) (**Figure 2A**). Morphologic feature enrichment was calculated for each cluster (**Table S3**) and identified cluster 8 exhibiting the highest Zta expression, indicative of cells undergoing lytic infection (**Figure 2B**). Accordingly, this phenotype and several others with Zta expression above background (clusters 7, 18, and 19) were exclusively composed of EBV-positive cells lines competent for lytic cycle entry (i.e., negligible presence of Namalwa) (**Figure 2C**).

**Figure 2.**
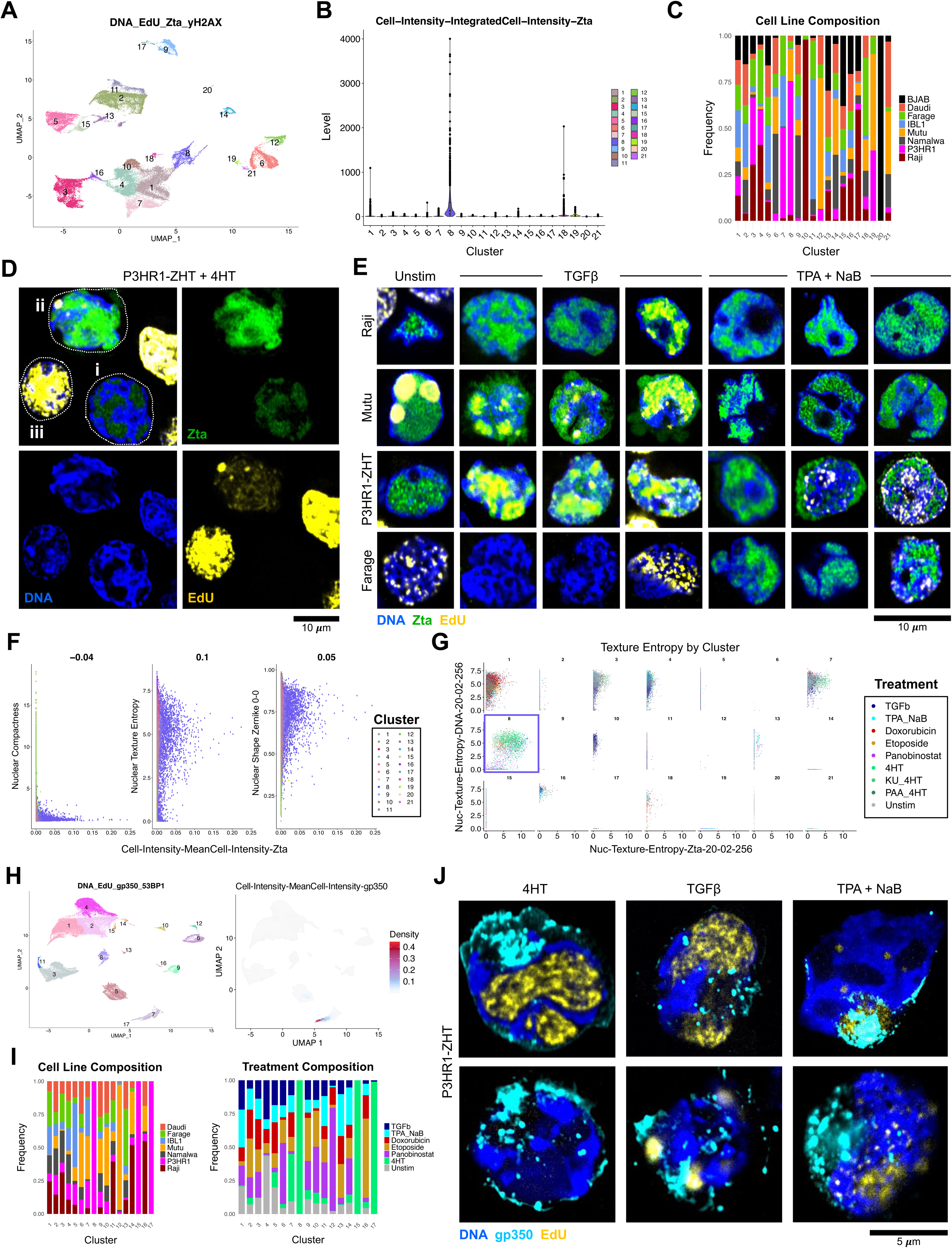
High-throughput quantification of EBV lytic cycle induction and treatment-stratified replication phenotypes in B cell models. **A**) UMAP representation and phenotypic clustering of segmented cells stained for nuclei, DNA synthesis, Zta, and γH2AX. **B**) Cluster-resolved quantification of whole-cell Zta immunofluorescence intensity from the imaging experiment depicted in (A). **C**) Normalized frequency of model cell lines within phenotypic clusters depicted in (A-B). **D**) Detail of cell state heterogeneity within P3HR1-ZHT cells treated with 4HT to induce the lytic infection program. One outlined cell (i) exhibiting a nucleus (blue) with low Zta intensity (green) within intranuclear compartments in the absence of *de novo* DNA synthesis signal (yellow) corresponds to cluster 1. Another cell (ii) exhibiting high Zta intensity overlapping with low DNA synthesis signal within intranuclear compartments corresponds to cluster 8. A third cell (iii) exhibiting high-intensity nucleus-wide DNA synthesis in the absence of VRCs or Zta expression corresponds to cluster 3. **E**) Single-cell images from selected lines and treatments depicting differential Zta expression, replication compartment, and DNA synthesis phenotypes. **F**) Cluster-resolved correlation of Zta intensity with nuclear shape and texture features. Individual points represent segmented cells color coded by cluster as in (A-B). **G**) Cluster-and treatment-resolved correlation between Zta texture entropy and nuclear texture entropy. Segmented cells are color-coded by treatment and split onto separate axes for each cluster. **H**) UMAP representation and phenotypic clustering of segmented cells stained for nuclei, DNA synthesis, gp350, and 53BP1 (left panel). UMAP representation of whole-cell mean gp350 intensity (right panel). **I**) Bar plots of normalized cluster composition by cell line (left panel) and treatment (right panel) for segmented cells presented in (H). **J**) Representative images of gp350+ P3HR1-ZHT cells (corresponding to clusters 7 and 17) from various lytic induction treatments. Nuclei are depicted in blue, gp350 is depicted in cyan, and DNA synthesis is depicted in yellow.

Images of cells from phenotype clusters were readily recovered from >350 GB of raw data using experiment metadata for well and sample IDs, file names, and pixel coordinates for segmented cell bounding boxes (**Figure S3**). This allowed us to easily link quantitative morphologic measurements and clusters with visual phenotypes to explore response heterogeneity across B cell models and treatments. For example, P3HR1-ZHT cells treated with 4HT exhibited Zta-positive cells in distinct stages of the lytic cycle and mostly Zta-negative cells interpreted as refractory to reactivation corresponding to different phenotype clusters (**Figure 2D**). Cells expressing Zta within intranuclear compartments in the absence of viral DNA synthesis (assayed via EdU incorporation) were consistent with a replication-independent reorganization of cellular chromatin (ROCC) phenotype (ROCC I) previously identified in hybrid epithelial/B cell lines (47) (**Figure 2D**, cell i). Cells in which Zta expression spatially overlapped EdU signal within intranuclear compartments corresponded to a kinetically later stage of reactivation characterized by active viral genome replication, replication compartment fusion, and progressive margination of host chromatin to the nuclear lamina (ROCC II) (47) (**Figure 2D**, cell ii). The intensity and spatial distribution of EdU signal in such lytic cells were markedly distinct from those of Zta-negative EdU-positive cells, which corresponded to S-phase based on elevated EdU intensity overlapping cellular dsDNA (**Figure 2D**, cell iii).

After confirming the presence of distinct lytic infection stages within induced P3HR1-ZHT cells, we examined effects of common lytic induction treatments on replication phenotypes of Zta-positive cells across model lines (**Figure 2E**). We first examined Raji cells and found Zta-positive cells were, with rare exceptions, negative for colocalized EdU signal within intranuclear compartments regardless of treatment. This was consistent with genetically defective viral genome replication (**Figure 2E**, first row). Looking at Mutu cells, Zta-positive cells with active EBV replication and ROCC were identified in TGF-β treatment and even rarely in unstimulated conditions; however, Zta-positive cells induced with a combination of the protein kinase C (PKC) activator phorbol 12-myristate 13-acetate (PMA, a.k.a. TPA) and the histone deacetylase inhibitor sodium butyrate (NaB) typically lacked EdU signal used to identify EBV genome replication (**Figure 2E**, second row). This differential response to TGF-β versus TPA + NaB treatment was also observed in P3HR1-ZHT (**Figure 2E**, third row). While some Zta-positive P3HR1-ZHT cells induced by TPA + NaB were EdU-positive, the lack of overlap between these signals indicated cellular rather than viral DNA synthesis. Zta-positive Farage cells exhibited a similar phenotype upon treatment with TPA + NaB, whereas TGF-β was not an effective lytic induction method for these cells (**Figure 2E**, fourth row). Quantification of whole-cell EdU intensity distributions by treatment condition across all cells demonstrated that TPA + NaB significantly depleted DNA synthesis relative to TGF-β and unstimulated conditions independent of its effect of Zta induction (**Figure S4A**). This effect resembled that of genotoxic drugs with current or prospective applications to treat B cell lymphomas (doxorubicin, etoposide, and panobinostat), some of which have been shown to induce expression of certain EBV lytic genes (27, 58). Notably, reduced DNA synthesis in TPA + NaB-treated cells persisted over time even when treatments were washed out within two hours (**Figure S4B**). This treatment effect is consistent with prior studies (59) and highlights an important consideration for the use of TPA + NaB to study the EBV lytic cycle versus lytic gene expression *per se*, as viral DNA replication is essential for progression from early to late kinetic stages and evolution of VRCs (47).

Feature enrichment in Zta-positive cells (cluster 8) identified several proxy morphologic metrics to assess ROCC during lytic infection. Lytic cell nuclei were comparatively large and porous, as measured by low nuclear compactness and high nuclear texture entropy and Zernike features (mathematical descriptors for object shape and pattern composition) (**Figure 2F**). Reasoning that cells with VRCs would exhibit highly entropic nuclear (reorganized chromatin) and entropic Zta (compartmentalized enrichment) signals, we examined these signal texture features stratified by phenotypic cluster and induction treatment. Overall, Zta-positive cells in cluster 8 exhibited higher nuclear and Zta texture entropy (signal non-uniformity or randomness) in response to TGF-β treatment (and 4HT for P3HR1-ZHT) versus TPA + NaB, doxorubicin, etoposide, and panobinostat (**Figure 2G**). Thus, quantitative morphologic measurements can delineate qualitatively different responses of latently infected cells to lytic-inducing stimuli.

A second survey experiment (DNA_EdU_gp350_53BP1) was performed to assay later stages of EBV reactivation across the same B cell models and treatments. Feature selection, dimensional reduction, and clustering were again used to define morphologic phenotypes (**Figure 2H**, left panel) including infrequent late lytic cells in clusters 17 (high gp350) and 7 (low gp350) (**Figure 2H**, right panel). The cell line composition of each cluster indicated that gp350-positive cells were derived from lytic-competent models as expected (**Figure 2I**, left panel). While low-expressing gp350-positive cells were produced by several treatments, high-expressing gp350-positive cells were largely derived from 4HT-stimulated P3HR1-ZHT and a small fraction of TGF-β-treated cells (**Figure 2I**, right panel). Inspection of representative gp350-positive P3HR1-ZHT cells from various treatments confirmed major nuclear chromatin reorganization in late EBV lytic stages (**Figure 2J**). While some gp350-positive cells were identified from TPA + NaB treatment, these cells generally exhibited reduced *de novo* DNA incorporation and more limited ROCC (i.e., compartment formation (ROCC I) without laminar margination of chromatin (ROCC II)) compared to cells stimulated with 4HT or TGF-β. Collectively, single-cell HCS survey experiments revealed line- and treatment-dependent lytic induction and established quantitative fluorescence metrics of reactivated cell morphology and EBV lytic progression.

### Distinct DNA damage responses in EBV-positive B cells resolved by treatment and lytic expression

Given the importance of the double-stranded break (DSB) repair arm of the cellular DDR to dsDNA virus reactivation (38), we aimed to define the kinetics and localization of lytic antigens and key DSB biomarkers. Whole-cell γH2AX and Zta intensity measurements highlighted cell line and treatment variability in the association between lytic induction and early-kinetics DDR (**Figure 3A**). TGF-β and TPA + NaB treatments yielded higher frequencies of Zta-positive cells than doxorubicin, etoposide, or panobinostat, which predominantly induced DNA damage in the absence of lytic protein expression. TPA + NaB induced higher intensity γH2AX than TGF-β in both Zta-positive and Zta-negative cells, indicating treatment-associated DNA damage independent of lytic gene programs. Subsets of P3HR1-ZHT cells treated with etoposide or panobinostat exhibited modest Zta immunofluorescence, however this may reflect stabilized expression of the line’s recombinant fusion of Zta with a murine estrogen receptor hormone-binding domain rather than *bona fide* endogenous Zta (*BZLF1*). P3HR1-ZHT treated with 4HT provided a positive control for cells co-positive for Zta and DDR markers, and reduced γH2AX upon co-treatment with the ataxia-telangiectasia mutated (ATM) serine/threonine kinase inhibitor KU-55933 confirmed the biologic specificity of γH2AX (a histone modification deposited by phosphorylated ATM kinase) (60). The apparent reduction of Zta intensity in P3HR1-ZHT cells co-treated with 4HT and the DNA synthesis inhibitor phosphonoacetic acid (PAA) was unexpected since PAA interferes with early and late (but not IE) lytic stages; however, this may be attributable to low cell recovery from the PAA + 4HT condition (n = 177 cells) in this experiment.

**Figure 3.**
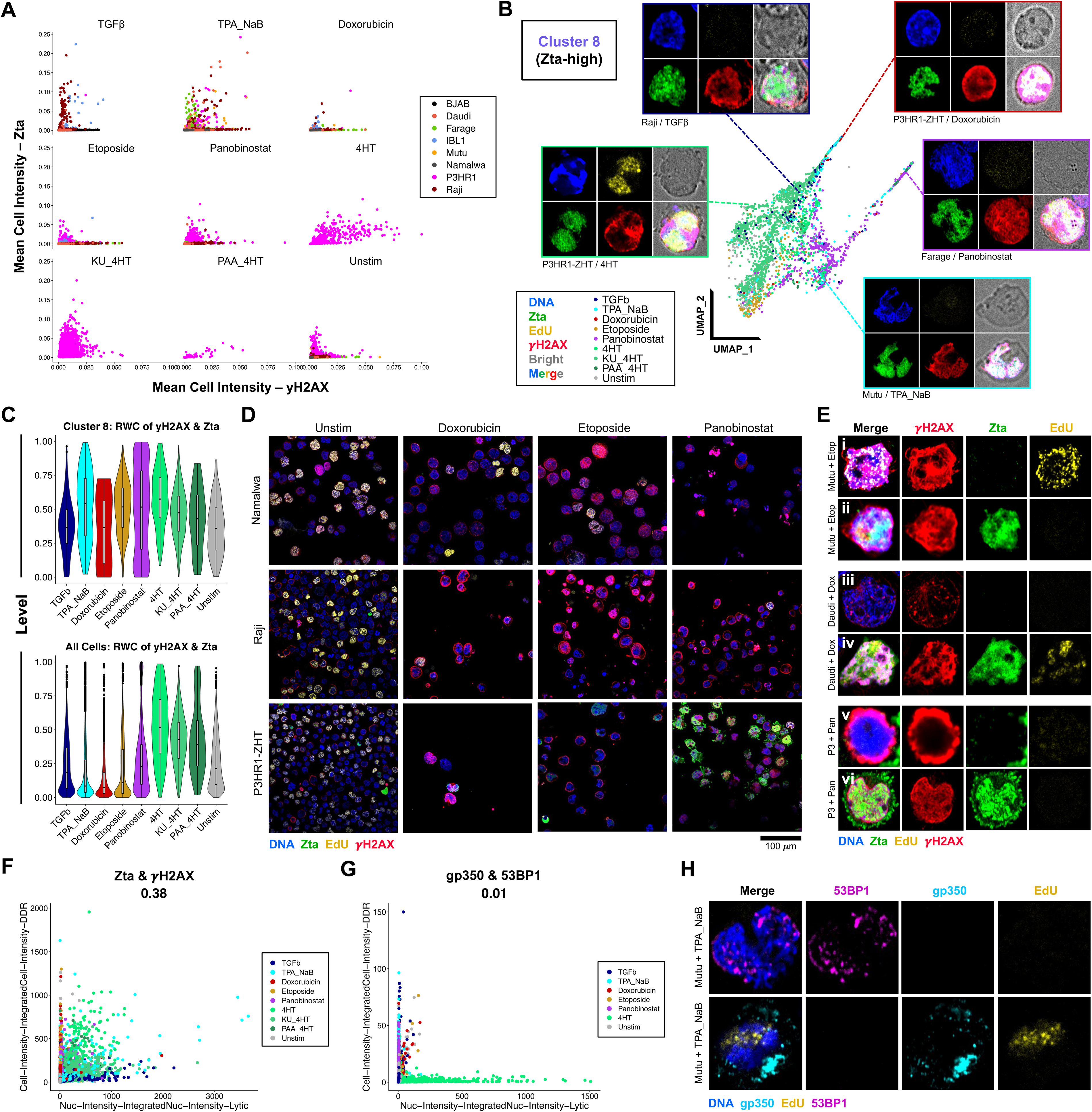
Genotoxin-induced reactivation and DNA damage responses differ quantitatively and qualitatively in B cell lymphoma models. **A**) Scatter plot of cell line- and treatment-stratified mean cell γH2AX intensity versus mean cell Zta intensity. **B**) Detail of Zta-high cluster replication and DNA damage phenotypes by treatment and B cell line. Merged and single channel images (nuclei in blue, Zta in green, EdU in yellow, γH2AX in red, brightfield in gray) are presented for example single-cell phenotypes. **C**) Whole-cell Zta intensity (top panel) and Zta nuclear entropy (bottom panel) distributions in lytic cells (cluster 8 in DNA_EdU_Zta_γH2AX staining experiment) stratified by treatment. **D**) Representative images of cell responses to genotoxins (60x magnification, full field of view). Images are shown for untreated (Unstim), 100 nM doxorubicin, 100 nM etoposide, or 50 nM panobinostat conditions in Namalwa (top row), Raji (middle row), and P3HR1-ZHT (bottom row) cells (nuclei in blue, Zta in green, EdU in yellow, γH2AX in red). **E**) Heterogeneous γH2AX stain patterns induced by genotoxic stress in Zta-negative and Zta-positive cells. Examples are shown for Mutu treated with etoposide (top two rows), Daudi treated with doxorubicin (middle two rows), and P3HR1-ZHT treated with panobinostat (bottom two rows). **F**) Scatter plot of correlation between integrated Zta and γH2AX signal in single cells from DNA_EdU_Zta_γH2AX experiment (color represents treatment). Pearson’s R = 0.38. **G**) Scatter plot of correlation between integrated gp350 and 53BP1 signal in single cells from DNA_EdU_gp350_53BP1 experiment (color represents treatment). Pearson’s R = 0.01. **H**) Example of 53BP1 staining in gp350-negative (top row) and gp350-positive (bottom row) Mutu cells stimulated with TPA + sodium butyrate (NaB).

Among Zta-positive cells, Zta expression and texture entropy were significantly lower for genotoxic treatments (**Figure S5**). This led us to interrogate whether the DDR phenotypes of lytic cells were likewise treatment-dependent (**Figure 3B**). In 4HT-stimulated P3HR1-ZHT cells with clear EBV VRCs, typical γH2AX staining overlapped host chromatin exhibiting ROCC. A similar pattern of spatially inverse Zta and γH2AX intensities was observed to a lesser degree in TGF-β-treated Raji cells. By contrast, rare Zta-positive cells from multiple cell lines treated with genotoxins often exhibited pan-nuclear γH2AX and diffuse Zta staining in the absence of obvious VRCs. Quantitatively, this difference was reflected in modestly higher rank-weighted correlation (RWC), an intensity-dependent measurement of signal overlap, between γH2AX and Zta in cluster 8 (lytic) cells (**Figure 3C**, top panel). This RWC trend was notably reversed across aggregated cell phenotypes for doxorubicin and etoposide but not panobinostat, likely due to the predominance of Zta-negative cells with extensive DNA damage in these conditions (**Figure 3C**, bottom panel and **Figure 3D**). We also investigated whether Zta-positive and Zta-negative cells within given cell lines and treatments exhibited distinct γH2AX profiles (**Figures 3E, S6-7**). Pan-nuclear γH2AX accumulation consistent with widespread DNA damage (and a pre-apoptotic state in some contexts) (61, 62) was the most common early DDR phenotype in Zta-positive cells induced by genotoxins, though other patterns were infrequently observed (**Figure S6A-C**). Zta-negative cells in equivalent treatments exhibited more diverse DDR phenotypes ranging from γH2AX foci (localized DNA repair), the pan-nuclear pattern, and annular staining indicative of early-stage apoptosis (63). The apoptotic ring γH2AX phenotype was nearly exclusive to Zta-negative cells, consistent with apoptosis resistance conferred by multiple EBV lytic genes (64, 65). Panobinostat was the most efficient Zta inducer among the chemotherapeutic agents we tested, particularly in P3HR1-ZHT and Farage cells. Panobinostat-treated Zta-positive cells generally exhibited pan-nuclear γH2AX staining without evidence of EdU incorporation or typical ROCC patterns (**Figure S6B-C)**. We also observed an interesting linear γH2AX stain pattern in several panobinostat-treated cell lines (particularly IBL1 and Farage) that appeared to correspond to long DNA strand domains (**Figure S6C-E**). In some cells, several such linear rays of γH2AX signal appeared to emanate from a shared focus. This may reflect HDACi-mediated nucleosome eviction, possibly in the absence of DSBs (66), and subsequent megabase-scale spreading (67) of the damage mark due to dysregulated DNA organization. We speculate that that this linear ray pattern may reflect the early-stage initiation (or a higher resolution view) of the pan-nuclear γH2AX phenotype. The morphologic correlation between Zta positivity and pan-nuclear γH2AX in response to genotoxins was manually confirmed to be statistically significant for doxorubicin and etoposide as a combined group (Chi-squared p = 1.83e-4, n = 34 cells; **Figure S7A**) and panobinostat (Chi-squared p = 1.25e-9, n = 195 cells; **Figure S7B**).

Given the correlation between IE and early indicators of lytic infection and DSB DDR (**Figure 3F**), we examined whether later kinetic biomarkers were similarly correlated. However, the late lytic glycoprotein gp350 and the late DDR protein 53BP1 exhibited no such correlation (**Figure 3G**). Upon closer examination of cells co-stained for gp350 and 53BP1 (DNA_EdU_gp350_53BP1), we found 53BP1 only in cells lacking gp350 (**Figure 3H**). This finding led us to investigate the coordinated kinetics of lytic infection and cellular DDR in greater detail.

### Depletion of cellular DNA damage biomarkers γH2AX and 53BP1 in EBV VRCs

To better understand EBV reactivation and DDR kinetics, we next investigated the co-expression of IE lytic and late DDR markers (DNA_EdU_Zta_53BP1) as well as late lytic expression alongside the early DDR (DNA_EdU_gp350_γH2AX). For these experiments, the number of cell lines (Namalwa, Mutu, P3HR1-ZHT, and IBL1) and treatments (Unstim, TGF-β, TPA + NaB, and H_2_O_2_) were limited to accommodate three post-treatment timepoints (24, 48, and 72 h) for each condition (n = 2 well replicates) on one 96-well plate per stain panel. This design enabled time-resolved confirmation that, despite being an effective Zta induction method, TPA + NaB treatment results in impaired / delayed late lytic expression associated with DNA synthesis inhibition (**Figures S8-S9**). A similar effect was observed for oxidative stress-induced lytic expression using H_2_O_2_ (**Figures S8-S9**). By contrast, TGF-β treatment yielded appreciable lytic induction with minimal impact on global EdU incorporation, more rapid accumulation of late lytic cells, and comparatively less DNA damage in non-reactivated cells (**Figures S8-S12**). Based on cluster- and treatment-level Zta expression, TPA + NaB appears to induce more widespread Zta expression across phenotypically diverse cells, likely at the expense of more physiological lytic responses elicited by treatments such as TGF-β (**Figure S10A**).

With this approach, we found total 53BP1 intensity was depleted in many Zta-positive cells (**Figures 4A, S10A**; cluster 9 in the DNA_EdU_Zta_53BP1 dataset). Conversely, γH2AX was positively correlated in gp350-positive cells (**Figures 4B, S10B**; cluster 10 in the DNA_EdU_gp350_γH2AX dataset). Collectively, these data indicated that the early DDR is active in early and late lytic cells while at least one critical mediator of the late DSB repair (53BP1) is diminished during EBV lytic infection. While γH2AX associates with the EBV lytic origin of replication (*oriLyt*) upon reactivation (52), we found that γH2AX was rapidly excluded from Zta-positive VRCs but remained intense across cellular chromatin (**Figure 4C**). This distribution is consistent with histone depletion from VRCs and the packaging of histone-free, epigenetically naïve EBV genomes in newly synthesized virions (68, 69). 53BP1 was likewise depleted in Zta-positive compartments with active viral genome replication, whereas Zta-positive cells lacking viral DNA synthesis typically retained 53BP1 signal coinciding with Zta-positive compartments (**Figures 4D-E**). This mutually exclusive 53BP1 and EdU in Zta-positive cells was evident from treatments with differential effects on DNA synthesis (e.g., TPA + NaB versus TGF-β) and from intra-sample phenotypic heterogeneity (**Figures 4D-E, S11A, S12A**). These results support 53BP1 recruitment to early-stage VRCs and depletion as EBV genome replication progresses.

**Figure 4.**
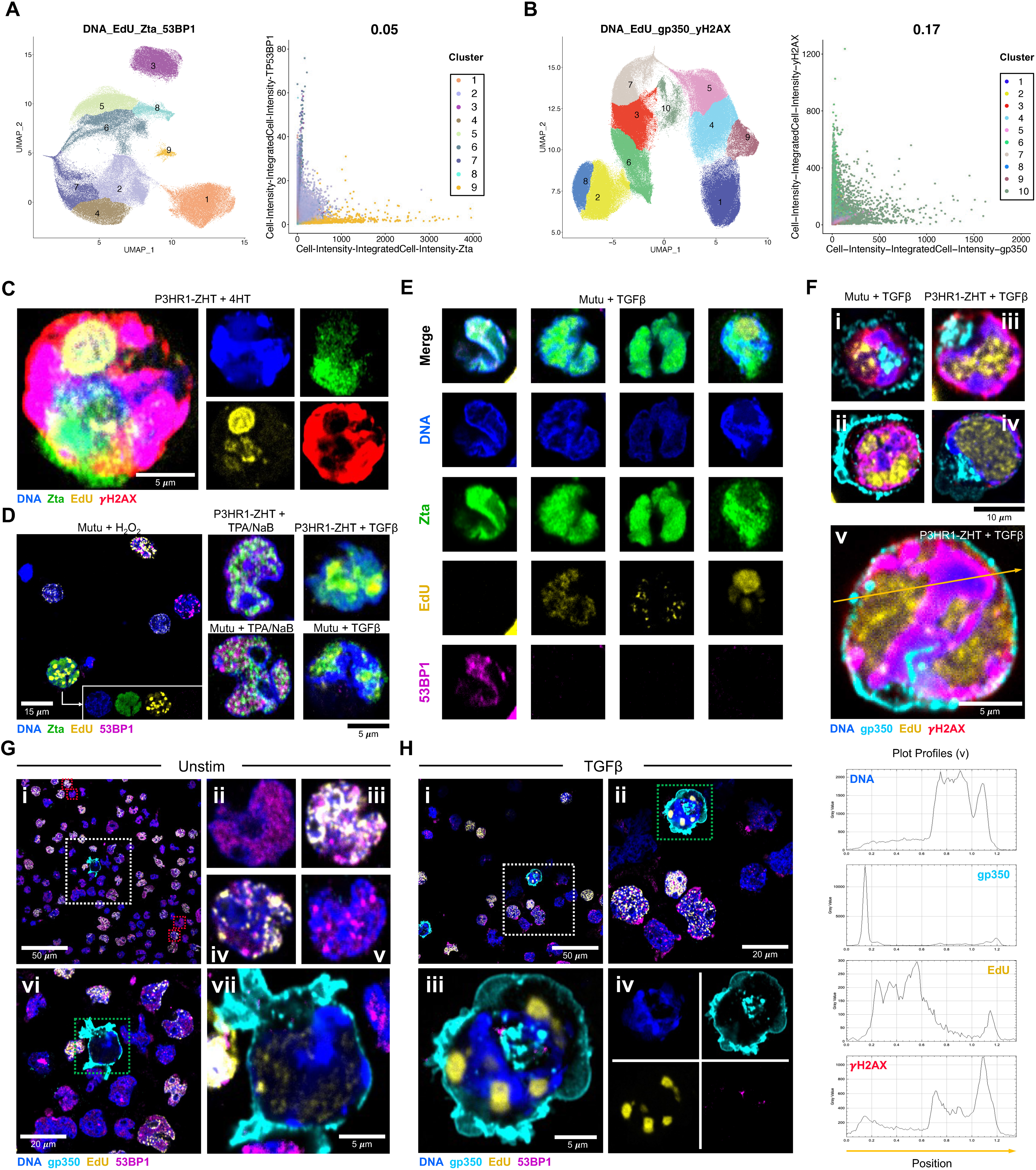
DDR biomarkers are depleted within EBV VRCs. **A**) Co-detection of IE lytic (Zta) and late DDR (53BP1) biomarkers over 72 h timecourse. UMAP representation and phenotypic clustering of segmented cells stained for nuclei, DNA synthesis, Zta, and 53BP1 (left panel). Scatter plot of integrated Zta vs 53BP1 intensity in cells by cluster, Pearson’s R = 0.05 (right panel). **B**) Co-detection of late lytic (gp350) and early DDR (γH2AX) biomarkers over 72 h timecourse. UMAP representation and phenotypic clustering of segmented cells stained for nuclei, DNA synthesis, gp350, and γH2AX (left panel). Scatter plot of integrated gp350 vs γH2AX intensity in cells by cluster, Pearson’s R = 0.17 (right panel). **C**) Detail of γH2AX exclusion from intranuclear VRCs in a P3HR1-ZHT cell stimulated with 4HT (nuclei in blue, Zta in green, EdU in yellow, γH2AX in red). **D**) Treatment-variable 53BP1 staining in Zta-positive Mutu and P3HR1-ZHT cells. Absence of 53BP1 detection in H_2_O_2_-reactivated Mutu cell with pan-nuclear Zta expression (left panel; individual channels in lower right corner). Presence of 53BP1 in Zta-positive EdU-negative cells induced with TPA + NaB (middle panels). Absence of 53BP1 in Zta-positive EdU-positive cells exhibiting VRCs induced by TGF-β (right panels). Nuclei depicted in blue, Zta in green, EdU in yellow, and 53BP1 in magenta for all panels. **E**) 53BP1 colocalization with Zta in VRCs and subsequent depletion associated with the onset of viral genome replication. Nuclei depicted in blue, Zta in green, EdU in yellow, and 53BP1 in magenta. **F**) Representative examples host genome-localized γH2AX in late lytic (gp350-positive) Mutu (panels i and ii) and P3HR1-ZHT (panels iii, iv, and v) cells reactivated via TGF-β treatment. Fluorescence intensity plot profiles for each channel correspond to the line depicted in panel v. Nuclei depicted in blue, gp350 in cyan, EdU in yellow, and γH2AX in red for all panels. **G**) Depletion of 53BP1 signal in late lytic (gp350-positive) P3HR1-ZHT cell without induction treatment (unstimulated). Full-field 60x magnification image (panel i); individual gp350-negative cells with diffuse 53BP1 staining (panel ii), replication-associated 53BP1 foci (panels iii and iv), and damage-associated foci in the absence of DNA replication (panel v) from dashed red boxes in panel i; detail of region including gp350-positive cell (panel vi); depletion of 53BP1 in gp350-positive cells with late VRC(panel vii, detail of dashed green box in panel vi). Nuclei depicted in blue, gp350 in cyan, EdU in yellow, and 53BP1 in magenta for all panels. **H**) Depletion of 53BP1 signal in late lytic (gp350-positive) P3HR1-ZHT cell treated with TGF-β. Full-field 60x magnification image (panel i); detail of region outlined in dashed white box in panel i including late lytic cells and non-lytic cells (panel ii); detail of late lytic (gp-350-positive) cell from dashed green box in panel ii (panel iii); individual fluorescent channels for gp350-positive cell depicted in panel iii (panel iv). Nuclei depicted in blue, gp350 in cyan, EdU in yellow, and 53BP1 in magenta for all panels.

As in Zta-positive cells, accumulation of γH2AX in gp350-positive cells from multiple cell lines and induction treatments was intense within cellular chromatin but depleted in VRCs (**Figures 4F, S12B**). We consistently observed this DDR pattern across cells exhibiting varying replication compartment morphologies and ROCC phenotypes. Interestingly, γH2AX intensity was generally greatest at interfaces between VRCs and adjacent cellular chromatin and attenuated with increasing distance into cellular chromatin regions. Despite intense host-targeted γH2AX, gp350-positive cells did not exhibit appreciable 53BP1 signal in either host or viral genomes for unstimulated and treatment-stimulated conditions (**Figure 4G-H**). Manual quantification of 53BP1 in Zta-positive versus Zta-negative cells in P3HR1-ZHT and Mutu cells stimulated with TGF-β confirmed significant 53BP1 depletion (punctate and diffuse patterns) in P3HR1-ZHT (Chi-squared p = 5.88e-5, n = 107 cells; **Figure S13A**) and Mutu (Chi-squared p = 1.54e-8, n = 125 cells; **Figure S13B**). These data collectively indicate that 53BP1 interactions with EBV lytic proteins, particularly Zta (41), are physically partitioned from virus-mediated initiation of an incomplete host-directed DDR evidenced by widespread γH2AX. Moreover, 53BP1 depletion from compartments in Zta- and EdU-co-positive cells as well as gp350-positive cells implies its importance for IE and early viral processes rather than late reactivation stages.

### Pseudotime trajectory analysis of host-virus morphologic dynamics

We primarily used DNA synthesis pulse labeling to identify EBV VRCs. In conjunction with nuclear staining, EdU labeling of host DNA replication also facilitated cell cycle staging in asynchronous populations with high-resolution S-phase replication timing (**Figure 5A**). Cells in early S-phase displayed comparatively low EdU intensity limited to internal (presumably, euchromatin-rich) nuclear regions. Cells in peak S-phase exhibited intense pan-nuclear EdU signal. Cells in late S-phase contained larger nuclei and had limited EdU incorporation localized to heterochromatin-rich topological boundaries (likely nuclear laminae and nucleoli). DNA and EdU intensities in Zta-positive cells were consistent with several cell cycle stages, though cells with intense Zta staining typically exhibited modest EdU signal (**Figure 5B**). This observation implied that Zta expression is attenuated within peak S-phase cells. The cellular environment during EBV lytic reactivation has been described as pseudo-S-phase in prior studies (39, 70); thus, we aimed to understand the progression of lytic infection relative to cell cycle stage in greater detail.

**Figure 5.**
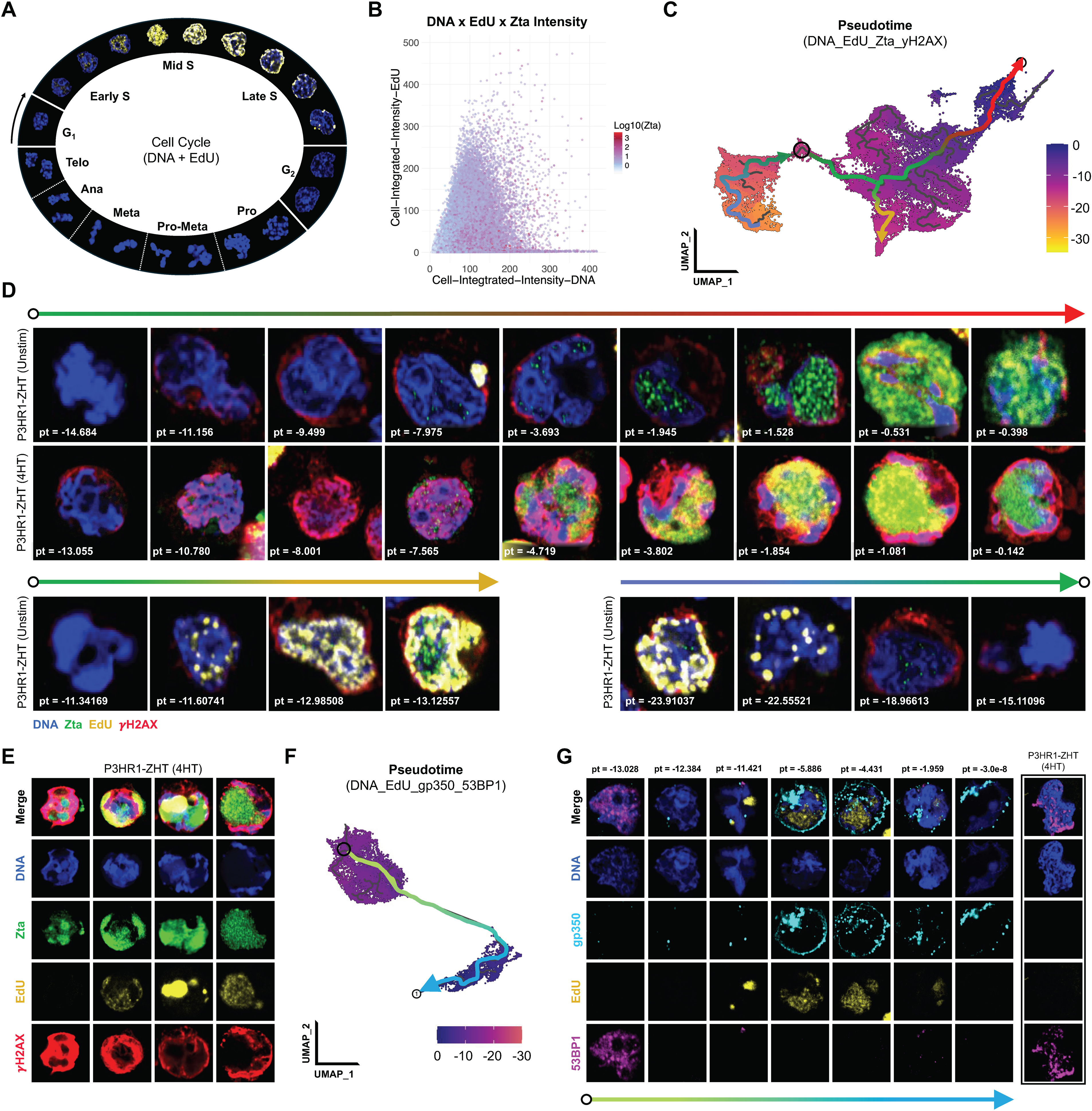
Pseudotemporal inference of EBV lytic infection and DDR phenotypic trajectories. **A**) High-resolution identification of cell cycle stages by DNA and EdU staining in unstimulated Namalwa cells (asynchronous proliferation). **B**) Zta expression resolved by cell cycle staging based on integrated DNA and EdU intensity. Points represent individual cells segmented from DNA_EdU_Zta_γH2AX staining and are color-coded by Zta intensity. Cells with low DNA intensity and low EdU intensity correspond to G1 and sub-G1 stages; cells with medium DNA intensity and elevated EdU intensity correspond to S-phase (and pseudo-S phase for reactivating EBV+ cells); cells with low EdU intensity and high DNA intensity correspond to G_2_/M phase. **C**) Morphologic pseudotime trajectories for DNA_EdU_Zta_γH2AX staining experiment. Pseudotime values were calculated relative to late lytic reactivation (pt = 0) determined by Zta expression and replication compartment morphology. Three morphologic state trajectories are denoted by color gradient lines with arrows indicating directionality. **D**) Phenotypic progression inferred along each of the three pseudotime trajectories annotated in (C). The green-to-red trajectory (top row) captures cells in the transition from latency to lytic reactivation. The green-to-gold trajectory (bottom row, left) captures cells entering the cell cycle and progressing through S-phase. The blue-to-green trajectory captures cells in the late stages of S-phase through G_2_/M and subsequent return to G_1_. Pseudotime values are provide for each depicted cell (pt value). Nuclei are presented in blue, Zta in green, EdU in yellow, and γH2AX in red for all images. **E**) Inferred progression (left to right) of nuclear morphology, Zta expression, VRCs, and host genome damage (γH2AX) through immediate early, early, and late lytic EBV infection for P3HR1-ZHT cells treated with 4HT. **F**) Morphologic pseudotime trajectory for DNA_EdU_gp350_53BP1 staining experiment. Pseudotime values were calculated relative to late lytic reactivation (pt = 0) determined by maximal gp350 intensity and replication compartment morphology. The morphologic state trajectory from latent to late lytic infection is denoted by color gradient line with arrow indicating directionality. **G**) Phenotypic progression inferred along pseudotime trajectory depicted in (F). The trajectory is initiated from a non-lytic (gp350-negative) cell in G_1_ (EdU-negative) exhibiting diffuse 53BP1 staining. Pseudotime values are provided for each cell as in (D). A non-lytic cell with 53BP1 foci identified from the same 60x field of view as several depicted late lytic cells is included for reference (outlined in black, not part of presented trajectory). Nuclei are presented in blue, gp350 in cyan, EdU in yellow, and 53BP1 in magenta for all images.

While lytic EBV cells are infrequent in B cell models, the scale and robustness of the HCS assay enabled the capture of thousands of lytic cells in asynchronous response phases. In lieu of live-cell tracking data, which is incompatible with EdU labeling, these thousands of cell snapshots effectively constitute out-of-order movie frames depicting dynamic lytic reactivation. Originally built to identify developmental differentiation trajectories from single-cell transcriptomes, pseudotime analysis (71) is a powerful graph-based approach for organizing heterogeneous yet continuous cell states from snapshot timepoints into ordered phenotypic progressions. Pseudotime analyses have been used previously to analyze EBV scRNA-seq data (8, 9). Because scRNA-seq and HCS datasets are similarly cell-by-feature quantitative matrices, we tested whether pseudotime trajectory analysis could resolve dynamic host and viral processes from morphologic profiles. Moncole3 (72) was used to calculate a contiguous pseudotime graph anchored from cells with the highest Zta intensity (pseudotime=0) in the DNA_EdU_Zta_γH2AX induction survey experiment (**Figure 5C**). Pseudotime values were calculated for cells based on graph distance from this anchor point (**Figure 5C**, dark purple cells at graph origin), where higher magnitude negative values indicated cells phenotypically distant from lytic cells. We also defined a common reference point (cells negative for EdU, Zta, and γH2AX) shared across several distinct pseudotime sub-trajectories (**Figure 5C**, black ring and gradient arrows). Segmented cell metadata was used to extract representative cells along each trajectory; for consistency, we highlighted P3HR1-ZHT cells for spontaneous and HT-induced reactivation (**Figure 5D**). One trajectory (green-red gradient arrow) captured the development of Zta-high cells, along which gradual emergence of punctate Zta signal concomitant with colocalized intranuclear compartment formation progressively increased toward more uniform high Zta intensity and compartment expansion (**Figure 5D**, top two rows). This phenotypic transition was consistent with Zta function as a pioneer transcription factor (73). Two additional trajectories corresponded to G_1_-to-S entry (green-orange gradient arrow) and S-phase exit followed by subsequent cell cycle progression (blue-green gradient arrow), consistent with manually curated cell cycle reference states. Cells at several stages (mostly in late S-phase) along each of these two trajectories exhibited sparse punctate Zta signal (**Figure 5D**, bottom row). We also identified progressive changes in nuclear morphology, Zta compartmentalization and intensity, viral DNA replication, and γH2AX profiles in 4HT-treated P3HR1-ZHT cells (**Figure 5E**). While untreated cells exhibited moderate perinuclear γH2AX staining, higher intensity host genome-wide γH2AX staining was present in all 4HT-treated Zta-positive cells and directly correlated with DNA intensity throughout the transition from ROCC type I to ROCC type II. Persistent EBV-mediated DNA damage has previously been reported in multiple cell lines without external stimulation (74). Additionally, the difference in γH2AX profiles between unstimulated and 4HT-treated cells may reflect a consequence of spontaneous versus overexpressed Zta. Similar analysis confirmed depletion of 53BP1 as viral replication commences (**Figure S14**).

Pseudotime analysis also recovered a lytic reactivation trajectory from cells stained for DNA, EdU, gp350, and 53BP1 (**Figure 5F-G**). The calculated graph was again anchored to cells with maximal lytic signal (pseudotime = 0), and reactivation progression was inferred beginning from a non-lytic reference cell exhibiting diffuse 53BP1 and the absence of gp350 and EdU. 53BP1 signal was virtually absent from all phenotypes with discernible VRCs (EdU-positive) or gp350. Early replication compartment nucleation in pseudotime preceded expression of gp350, which became intense across cells with expanded and fused VRCs as well as apparent lysed cells with fully marginated host chromatin and no EdU signal. This phenotype sequence matched known viral lytic kinetics (75) and the dependence of late-stage ROCC (type II) on viral genome replication, thus demonstrating the accuracy and utility of morphologic pseudotime reconstruction to study viral infection dynamics using high-throughput snapshot imaging data.

### HCS assay customization to study lytic cell biology

Plate-based HCS methods and downstream analyses can be adapted readily to study other biological aspects and dynamics of infected cells. To demonstrate the biological versatility of this method, the same cell line and treatment conditions previously surveyed for lytic and DDR responses were stained for nuclei, mitochondria, and Zta and processed into dimensionally reduced maps of single-cell phenotypes (**Figure 6A**). Cells with high Zta expression in this experiment were identified (clusters 4 and 17) and examined for associated mitochondrial signal features (**Figures 6B-C**). This led us to identify differential mitochondrial morphology in Zta-negative cells, Zta-positive cells with the ROCC I phenotype, and Zta-positive cells with the ROCC II phenotype (**Figure 6D**). These states were exemplified in P3HR1-ZHT cells lacking stimulation (Zta-negative; **Figure 6D**, top row), cells pre-treated with PAA prior to 4HT to impair replication-dependent lytic progression (ROCC I; **Figure 6D**, middle row), and 4HT-induced cells (ROCC II; **Figure 6D**, bottom row), respectively. Mitochondrial localization in most cells was anisotropic and compact. We noted a progressive transition from fused filamentous mitochondrial networks in Zta-negative cells (**Figure 6D**, top row) toward mitochondrial fission (globular fragments) in Zta-positive cells exhibiting late VRCs (**Figure 6D**, bottom row) suggestive of mitophagy (76) and consistent with morphologic changes seen for Zta overexpression (77). In Zta-positive cells displaying marginated host DNA, fragmented mitochondrial networks were present and characteristically juxtaposed concave nuclear regions (**Figure 6D**, bottom row). Co-staining for mitochondria and gp350 in a separate experiment further supported accumulation of fragmented mitochondria proximal to sites of EBV virion assembly and release in P3HR1-ZHT cells (**Figure 6E**). This finding was confirmed in Mutu cells treated with TGF-β via immunostaining for gp350 and TOM20 in addition to nuclei and pulsed EdU incorporation (**Figure 6F**). These experiments highlight the modularity of single-cell HCS to support biologically tailored studies of concomitant viral and cellular features or processes.

**Figure 6.**
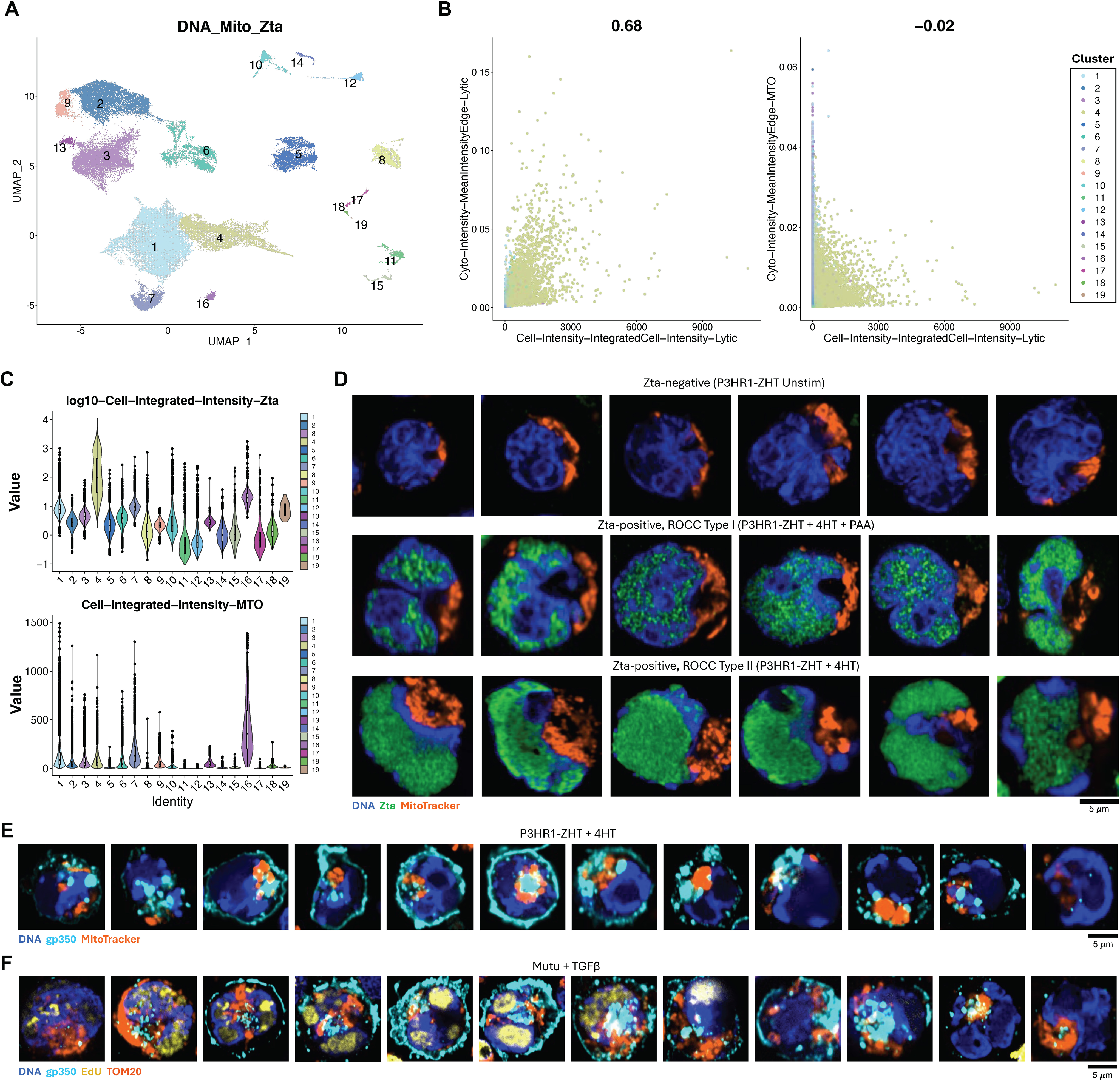
Mitochondrial network reorganization and colocalization during late EBV lytic infection. **A**) UMAP representation and phenotypic clustering of segmented cells stained for nuclei, mitochondria (MitoTracker), and Zta. **B**) Scatter plots of cytoplasmic region edge intensities for Zta (left panel, y-axis) and MitoTracker (right panel, y-axis) versus whole-cell integrated Zta intensity (both panels, x-axis). Points are color-coded by phenotypic cluster as shown in (A). Pearson’s R for pairwise feature correlation across all clusters are denoted above each scatter plot. **C**) Cluster-resolved quantification of Zta intensity (log10-transformed) for DNA_Mito_Zta staining experiment (top panel). Cluster-resolved quantification of integrated mitochondrial stain intensity for the DNA_Mito_Zta experiment (bottom panel). Clusters are color-coded as in (A-B). **D**) Representative images of nuclear and mitochondrial morphology in Zta-negative and Zta-positive cells by reorganization of cellular chromatin (ROCC) phenotype. Zta-negative cells (top row) are presented from unstimulated P3HR1-ZHT images. Zta-positive cells exhibiting ROCC type I (middle row; replication compartment formation without laminar margination of chromatin) are presented from P3HR1-ZHT cells co-treated with 4HT (lytic induction) and Phosphonoacetic acid (PAA; viral DNA synthesis inhibitor). Zta-positive cells exhibiting ROCC type II (bottom row; cellular chromatin margination to nuclear lamina) are presented from P3HR1-ZHT cells treated with 4HT alone. Nuclei are depicted in blue, Zta in green, and mitochondria in orange for all images. **E**) Representative images of nuclear and mitochondrial morphology in late lytic P3HR1-ZHT cells (gp350-positive, ROCC). Cells are approximately ordered by lytic progression (left-to-right; transiently increasing gp350 and transition from ROCC type I to ROCC type II). Nuclei are depicted in blue, gp350 in cyan, and mitochondria in orange for all images. **F**) Representative images of nuclear and mitochondrial morphology with viral DNA replication in late lytic Mutu cells following TGF-β treatment (gp350-positive, VRCs, ROCC). Cells are approximately ordered by lytic progression (left-to-right; transiently increasing gp350 and EdU intensity along with transition from ROCC type I to ROCC type II). Nuclei are depicted in blue, gp350 in cyan, and mitochondria in orange for all images. Mitochondria were detected by immunofluorescence (TOM20) instead of MitoTracker for this experiment.

### Method generalizability to widefield imaging systems

While previous experiments employed a laser-based, confocal high-content imaging system with high numerical aperture water immersion objectives, widefield epifluorescence imaging may provide comparative advantages (e.g., fluorophore compatibility, live-cell studies, and cost) for some applications at the expense of spatial resolution. Thus, we investigated whether HCS and morphologic pseudotime analyses were extensible to widefield imaging data. EBV-positive BL lines (Mutu, Daudi, P3HR1-ZHT, and Jijoye) were prepared as in prior experiments, stained with a five-color fluorescence panel (nuclei, Zta, mitochondria, the proliferation marker Ki-67, and near-infrared viability dye), and imaged at 40x magnification with real-time computational deconvolution (**Figures 7A-B, S15A-B**). Cell segmentation and measurement were performed with a modified pipeline (**Figure S15C**, **File S3**), and measurement matrices were analyzed to produce dimensionally reduced data representations by line, treatment, and morphologic phenotype (**Figure 7C-E**). Unsupervised clustering yielded a clear population of lytic cells exhibiting high nuclear texture entropy (as in confocal experiments) and model-dependent reactivation efficiency (**Figure 7F-G**). Pseudotemporal ordering likewise delineated latent cell states from progressive reactivation evidenced by Zta intensity, host chromatin, and mitochondrial morphology (**Figure 7H**). As for confocal experiments, we observed the gradual accumulation of Zta intensity and nuclear chromatin deformation along the pseudotime reactivation trajectory from widefield data (**Figure 7H**). These data show proof-of-principle that HCS and pseudotime methods can be successfully adapted for imaging platforms over a range of performance and technical specifications.

**Figure 7.**
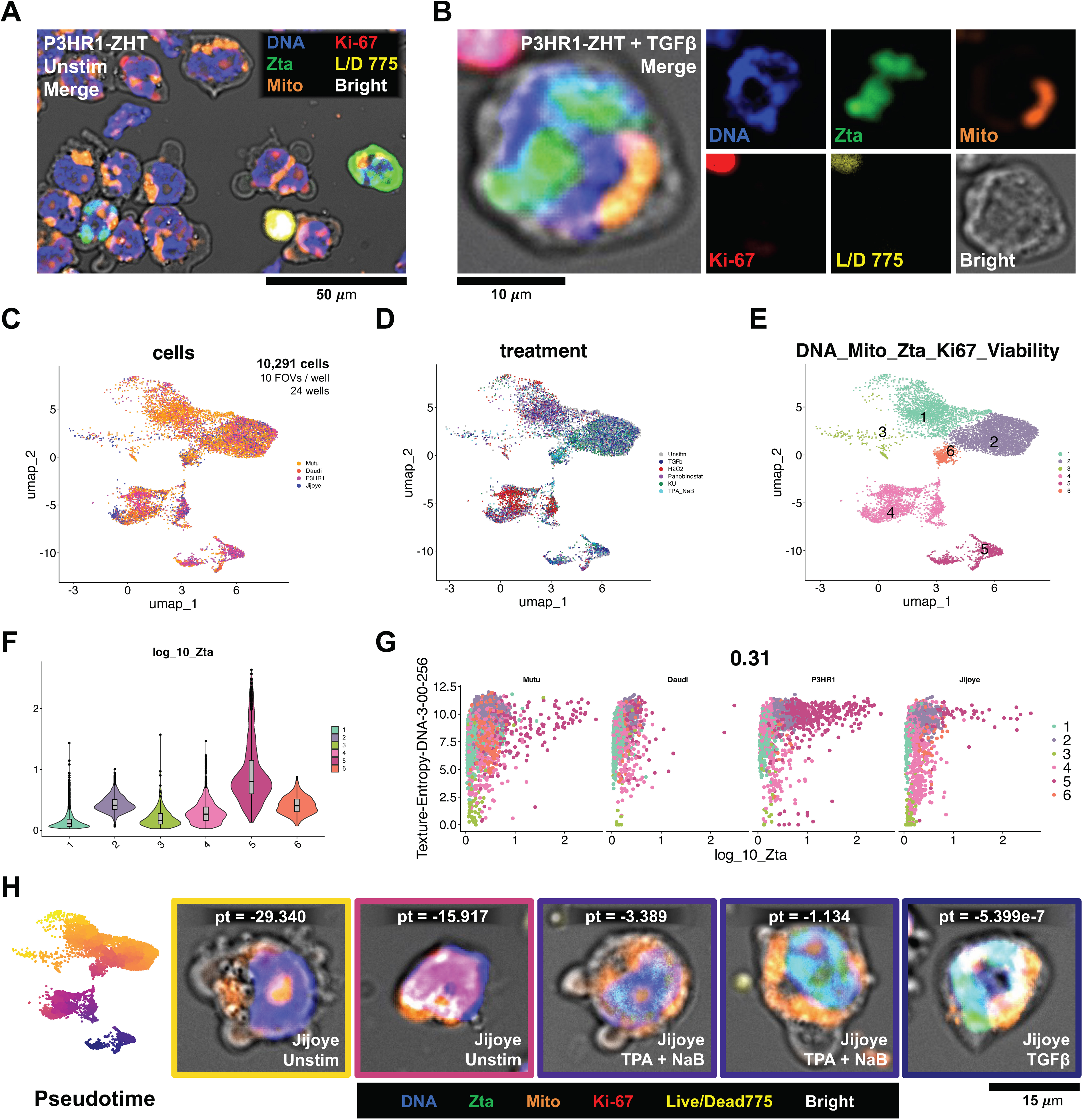
Generalizability of HCS and morphologic pseudotime analysis across imaging systems. **A**) Image detail for six-channel (five epifluorescence + one transmitted light) widefield microscopy with computational deconvolution using Leica THUNDER instant computational clearing (ICC) method (40x magnification, NA = 0.95). DNA in blue, Zta in green, mitochondrial dye in orange, Ki-67 in red, near infrared viability dye (live/dead NIR 775) in yellow, and brightfield in gray. **B**) Detail of lytic cell with VRCs from six-channel ICC imaging of P3HR1-ZHT cells treated with TGF-β. **C**) UMAP dimensional reduction of widefield ICC image data across four cell lines and six treatment conditions. Ten FOVs were analyzed for each of 24 wells (n = 10,291 cells post-QC). Cells are color-coded by original cell line (Mutu, Daudi, P3HR1-ZHT, and Jijoye). **D**) As in C, with cells color-coded by treatment condition (Unstimulated, TGF-β, H_2_O_2_, Panobinostat, KU-55933, and TPA + NaB). **E**) As in C-D, with cells colored by phenotype based on Leiden clustering. **F**) Log-normalized Zta intensity distribution by morphologic phenotype. Clusters are color-coded as in E. **G**) Correlation between Zta immunofluorescence intensity and DNA signal texture entropy (identified as a quantitative metric for cellular chromatin reorganization in confocal imaging experiments). Data are stratified by cell line and color-coded by morphologic cluster. Pearson’s R = 0.31 for dataset. **H**) Pseudotime trajectory analysis of widefield ICC morphologic phenotypes. Lytic cells in cluster 5 are designated as pseudotime = 0. Image channels are color-coded as in A-B.

## Discussion

EBV and many other DNA viruses form distinctive genome VRCs in host nuclei (78), wherein inter-species genome conflicts are manifest (79). Prior studies have investigated EBV replication compartment formation in B95-8 marmoset lymphoblast (40, 44), adherent epithelial (42, 45, 80), and hybrid epithelial / B cell (45, 47) lines with chemically-inducible Zta expression vectors. By developing a HCS-based assay to examine EBV reactivation and host responses in B cells, we have substantially extended insights beyond these prior investigations. Specifically, our approach quantitatively dissects viral reactivation dynamics in B cells treated with physiologically and pharmacologically relevant stimuli at high throughput with minimal manual curation bias. Our findings confirm the formation of distinct VRC architectures in multiple B cell lymphoma models using vector-based induction and physiologically relevant stimuli. Consistent with findings from a recent library screen of viral mutants (47), early replication compartment formation in EBV-infected B cells induces partial reorganization of cellular chromatin (ROCC type I) independent of viral DNA synthesis, which is necessary for subsequent compartment fusion and margination of host DNA to the nuclear lamina (ROCC type II). ROCC quantification via nuclear signal texture and other features supported pseudotemporal reconstruction of replication compartment evolution that revealed nascent compartment nucleation coinciding with the AP-1 homolog Zta. While modest Zta expression was occasionally detected in late S-phase and G_2_ cells identified by nuclear staining and EdU replication timing, full lytic reactivation pseudotemporally originated from Zta-positive G_1_ cells. This aligns with Zta-mediated cyclin-dependent kinase inhibitor (CDKi) activation that restrains entry into a normal cellular S-phase (81) and favors viral induction of a pseudo-S-phase cellular environment that supports compartmentalized EBV replication while blunting checkpoint-mediated arrest (39, 70). Unlike in reactivation-competent P3HR1-ZHT and Mutu lines, the defective transition from ROCC I to ROCC II apparent in Raji cells is consistent with the line’s inactivating mutation in *BALF2*, which encodes an early lytic ssDNA binding protein essential for viral genome replication (44, 47).

HCS also revealed substantial qualitative and quantitative heterogeneity in VRC formation following exposure to common physiologic and pharmacologic lytic stimuli (82). TGF-β treatment induced ROCC type I with Zta-positive compartments in multiple BL lines (Raji, Mutu, P3HR1-ZHT), however this response was not observed in either DLBCL lines we examined (Farage, IBL1). Prior studies have demonstrated downregulation of type II TGF-β receptor mediated by EBV (83) and functional interference with downstream SMAD signaling by herpesvirus (and virus-induced cellular) microRNAs (84–88), which may explain cell line-dependent Zta induction. Since it did not compromise DNA synthesis, TGF-β treatment was well-suited to study full reactivation trajectories in responsive lytic-competent cells (Mutu, P3HR1-ZHT). Though TPA + NaB effectively induces Zta (89) and, indirectly, its transcriptional target early lytic genes (73, 90), our data demonstrate that even transient TPA + NaB treatment impairs DNA replication and thus attenuates or delays late lytic infection by interfering with viral genome replication, similar to the effect of phosphonoacetic acid (91). This effect is independent of lytic gene expression and consistent with prior studies demonstrating reduced DNA synthesis in TPA-treated Namalwa cells (59). Notably, reduced colocalization between Zta and EdU in co-positive cells suggests that TPA + NaB treatment may modestly affect replication compartment formation. H_2_O_2_ treatment, which induces Zta expression and activity (92, 93), yielded a similar but transient effect on DNA synthesis. Thus, treatments that induce the complete lytic cycle via different mechanisms likely impact the timing and efficiency of viral gene expression and virion production due to indirect effects.

Clinically relevant genotoxins also induce EBV IE and early lytic gene expression (27). Doxorubicin-induced lytic reactivation has been demonstrated previously (26, 27) and was indeed observed in the present study in small Zta-positive subpopulations (typical < 2% of cells) in multiple tested cell lines. Rare doxorubicin-treated cells co-positive for gp350 and EdU indicate that this treatment permits the complete lytic cycle unless perturbed by viral DNA synthesis inhibitors (i.e., “kick-and-kill” regimens) (26). Etoposide and panobinostat also induced Zta-positive cells across lines, however the absence of late lytic phenotypes observed in these treatments suggest predominantly abortive reactivation. Increased efficiency of panobinostat-induced Zta in P3HR1-ZHT may derive from enhanced expression of the engineered fusion construct, however successful Zta induction in Raji and Farage cells indicates a genuine response of endogenous EBV genetic elements to panobinostat. Analogous to chemotherapy-dependent EBV lytic initiation (27), the apparent differences in genotoxin-dependent lytic progression are noteworthy and warrant consideration of the type(s) of DNA damage versus chromatin damage (94) induced by each drug. Doxorubicin and etoposide are both anthracycline topoisomerase II poisons (95, 96) that promote chromatin damage via nucleosome eviction (97, 98), however doxorubicin also directly generates mechanistically independent DNA DSBs (97, 99). HDAC inhibitors (e.g., panobinostat, sodium butyrate) also remodel chromatin (100) via a distinct mechanism without direct DNA strand damage. Interestingly, HDACi-mediated nucleosome eviction induces strand break-independent pATM activation (66), which would facilitate EBV reactivation (39, 82, 101). This effect on higher-order chromosome structure appears consistent with the latent-to-lytic switch observed upon disruption of chromatin insulators that suppress lytic gene promoters (102–106). However, HDAC inhibitor-driven nucleosome eviction may be kinetically premature for lytic infection, since reactivation-induced γH2AX association with the EBV *oriLyt* promoter (52) would be lost upon histone H2A.X removal and early lytic gene transcription occurs from chromatinized viral genomes (69). While extensive studies are clearly needed in the future, it is conceivable that direct DNA damage by genotoxins may initiate specific damage responses conducive to EBV lytic cycle initiation and completion (DSB DDR), whereas chromatin damage-inducing agents may only support partial reactivation.

Independent of whether and how genotoxins induce EBV reactivation, our data reveal distinct damage responses in chemotherapy-induced lytic cells versus non-lytic cells in the same treatment. The pan-nuclear γH2AX profile in Zta-positive cells commonly observed across physiologic and pharmacologic treatments contrasts with genotoxin-induced classic damage foci and annular patterns (the “apoptotic ring”) (63) in Zta-negative cells. In the context of radiation-induced DNA lesions, pan-nuclear γH2AX does not strictly correlate with widespread damage or apoptosis and can be reversible (62). Mutual exclusivity between Zta-positive cells and cells with annular γH2AX comport with prior demonstration of lytic cell apoptotic resistance (65). Likewise, the absence of focal 53BP1 accumulation in Zta-positive cells treated with doxorubicin, etoposide, or panobinostat (despite being enriched in Zta-negative cells under the same treatments) implies defects in p53-mediated arrest (107) mechanistically distinct from Zta interference with p53 (108, 109). Impaired 53BP1 focus formation is likely explained by functions of several viral proteins. Interaction of the tegument protein encoded by BKRF4 with histone H2A-H2B dimers impairs formation of 53BP1 foci by preventing DNA assembly onto nucleosomes (49, 110, 111), and the viral DNA polymerase processivity factor BMRF1 (EA-D) interacts with the NuRD complex in a manner that disrupts RNF168-mediated 53BP1 ubiquitination (50), which is critical for DSB repair (112). It is noteworthy that BKRF4 and BMRF1 are both transcriptionally early lytic genes expressed prior to viral DNA replication (75), which temporally aligns with our finding that 53BP1 depletion in lytic cells occurs concomitant with or preceding detectable genome amplification. Understanding such subversion of cellular DDR machinery in response to genotoxic stress bears particular clinical relevance and warrants further study since DDR abrogation can promote genomic instability. While lytic induction may sensitize some infected cells to antiviral drugs and expand the targetable antigen repertoires of EBV-positive tumors, such an approach may be double-edged given the impact of lytic infection, including abortive reactivation, on DNA damage responses and genomic instability (6, 113, 114).

Our data provide a unique view of DDR spatiotemporal dynamics during EBV lytic infection, including depletion of γH2AX and 53BP1 from compartments with detectable viral genome replication. The early importance of γH2AX (51, 52) and 53BP1 (41, 43) and their later exclusion from VRCs underscores the changing landscape of required versus antagonistic DDR components during reactivation stages. This in turn likely reflects the dynamic nature of viral DNA lesions and epigenetic topology from initially chromatinized viral genomes; early nucleosome eviction; strand nicking, ssDNA displacement, and stabilization during rolling circle amplification; and DSB generation as concatemers are processed into individual genomes for packaging (69). The spatiotemporal dynamics of 53BP1 are particularly intriguing in the context of viral genome topology – 53BP1 is present in early compartments when viral genome DSBs are presumably infrequent but is depleted from late compartments in which DSBs are generated by concatemer processing. This timing would seem to preclude potential γH2AX-independent recruitment of 53BP1 to DSBs (115). By contrast, components of the MRE11-RAD50-NBS1 (MRN) complex (which can also be recruited to DSBs independently of γH2AX (116)) appear to associate with newly synthesized viral DNA (40), though the precise timing of intra-compartment MRN in lytic cells merits further study. While host factor repair of viral genome DSBs generated by concatemer processing has been speculated (40), such repair would seemingly interfere with linear viral genome packaging in new virions. Alternatively, it is conceivable that DSB-binding host proteins might protect viral genome ends while downstream repair factors and damage signaling are subverted.

The loss of γH2AX in VRCs starkly contrasts with the modification’s ubiquity across host chromatin through early and late lytic stages. Host-directed γH2AX accumulation has been observed during KSHV reactivation (117, 118), potentially suggesting a conserved mechanisms among gammaherpesviruses. Interestingly, similar pan-nuclear γH2AX induction is also observed during primary adenovirus infection (119). Considering the absence of 53BP1 focus formation observed here and by others (42, 48, 117), persistent pan-nuclear γH2AX may perturb genome stability and host transcription through global chromatin remodeling in lytic cells while checkpoint-mediated arrest is abrogated. Whereas viruses gain advantage from orchestrating dynamic damage responses conducive to replicating their own genomes, so too may they benefit from a host genome stalled in a comparatively static and incomplete phase of the damage response.

Might such widespread epigenetic alteration of host genome organization due to impaired DDR mechanistically repress cellular genes? Reduction of host gene expression (“host shutoff”) is a hallmark of reactivation for many viruses and is classically attributed to viral factors that directly or indirectly promote host mRNA decay (120). In the human gammaherpesviruses, KSHV SOX (121) and EBV BGLF5 (122) exhibit RNA endonuclease activities that degrade cellular (and viral) transcripts. However, RNA-sequencing of lytic infection with recombinant Δ*BGLF5* EBV revealed that host transcript reduction is primarily BGLF5-independent, and many host transcript aberrantly retain introns or evidence of gene readthrough (14). As noted by Casco *et al* (14), cellular chromatin compaction driven by the expansion of VRCs is likely another source of diminished transcriptional machinery access to chromatin. DNA lesions including DSBs can block or stall RNA Pol II processivity (123). Notably, γH2AX is an important component of ATM-dependent transcriptional repression of genes proximal to DNA DSBs (124–126), albeit in contexts of 53BP1 focal accumulation. Conversely, slow kinetic ATM-mediated DSB repair can occur in heterochromatin (127), and such repair notably involves local chromatin decondensation and transition to nearby euchromatin (128–130). Indeed, transiently increased chromatin accessibility at DNA damage sites is an essential step for proper DDR in the “Access-Repair-Restore” model (131). Thus, it is also worth considering whether EBV-mediated interference with the restoration of repressive epigenetic status in γH2AX-positive heterochromatin regions lacking DSBs, possibly in conjunction with Zta-mediated chromatin remodeling (34, 35, 73), may contribute to aberrant cellular transcription. Many cellular transcripts during lytic infection exhibit intron retention and gene readthrough and may thus be non-functional (14); notwithstanding, multiple herpesviruses activate transcription of genes typically silenced after early development and, for at least some genes, their translation (9, 132, 133). One such gene, the transcription factor DUX4, is critical for reactivation of herpesviruses and other DNA viruses (133). For oncogenic viral infections, this degree of host reprogramming and plasticity is suggestive of a cancer hallmark (134). Given the importance of lytic gene expression for oncogenesis (6, 113, 135) and evidence that cells can remain viable after lytic reactivation (6, 114, 136), future studies should investigate the impact of chromatin remodeling stemming from viral subversion of host DNA damage responses on potentially pathogenic gene dysregulation. From a technical standpoint, HCS is ideally suited for such future work.

Early implementations of HCS as a standalone method included basic studies of cells infected with filoviruses (137) and influenza A (138), and was more recently applied to study heterogeneity in BK polyomavirus infection (139). However, HCS has not yet been adopted widely as a technique for virology. We propose that the HCS approach demonstrated herein will be broadly advantageous for studies of other viruses and their host cells since infection outcomes vary widely between individual cells. Such outcomes of primary infection include rapid sensing and neutralization by host innate immune systems (140, 141), distinct forms of cell death (142–144), and long-term persistence (145, 146). Cells harboring latent viruses exhibit additional biological heterogeneity depending on dynamic host-pathogen interactions (106, 147). Lytic viral replication likewise constitutes a major source of heterogeneity (148) including incomplete (abortive) versus successful outcomes and, in certain cases, variation in virion burst sizes among cells (149, 150). Notably, infrequent or rare infection phenotypes may be key determinants of viral pathogenesis (151). This phenotypic breadth highlights an imperative for using high-throughput single-cell approaches to better understand fundamental aspects of infections and virus-associated diseases. Our demonstrated HCS workflow offers major practical and biological advantages over conventional techniques to address these challenges. The developed plate-based assay supports high-throughput screening of EBV in its native host cell type, and robust retention of thousands of cells per condition greatly enhances the study of rare but critical infection phenotypes. The approach is readily customizable to investigate host-virus interactions of interest, including biological processes that are challenging to study in live cells. This resulting automated quantification reduces bias in image data interpretation and mitigates potentially unrepresentative low-throughput sampling; clustering and dimensional reduction support systematic phenotype discovery and streamlined exploration of raw image datasets that easily exceed hundreds of gigabytes; and pseudotime application to large asynchronous infected cell populations enables reliable inference of dynamic host-virus interactions from confocal or epifluorescence data without live-cell tracking. We have also provided a template R script with instructions for loading, processing, and analyzing CellProfiler (53) morphologic measurement files with Seurat v5 (55) and monocle3 (71, 72) to help users with less image processing and programming experience (**File S1**). Especially useful future applications of this technique include morphology-based screening (152–154) to identify new antiviral compounds and mechanisms of action as well as investigations of host and viral gene functions in different phases of infection. By coherently reconstituting the many dynamic “attitudes” of cells in motion (155), we hope this methodology will catalyze research in the accelerating discipline of single-cell virology and be broadly applied to study biological phenomena for which live single-cell imaging is intractable.

## Limitations of the Study

The assay demonstrated herein is not currently compatible with live-cell imaging of cells grown in suspension culture. The number, sequence, and complexity of image processing steps required for accurate cell segmentation will vary across model systems and cell types. We speculate that successful pseudotime implementations will require detection of at least one phenotypically continuous cellular structure or process. Accurate inference of rapid or sharp state transitions may be challenging due to reduced probability of detecting sufficient intermediate phenotypes. Accurate interpretation of morphologic states and dynamics require manual validation and should be informed by prior knowledge of relevant host-virus interactions if available. Supervised machine learning and other methods for trained phenotypic classification may be preferable to purely unsupervised methods for some applications. In the absence of effective isolation strategies (e.g., FACS, magnetic separation), validation and downstream analyses of rare phenotypes will be challenging. The depth of phenotypic profiling is limited by the number of acquired non-redundant image channels. Thus, future efforts are needed to achieve higher dimensional target detection, possibly through iterative cycles of detection, imaging, and washing.

## Materials and Methods

### Cell lines

All cell lines (BJAB, Namalwa, Raji, Mutu I, Daudi, P3HR1-ZHT, IBL1, Farage) were a generous gift from the Luftig Lab (Duke University School of Medicine, Durham, USA). BL-derived cell lines include BJAB, Namalwa, Raji, Mutu I, Daudi, P3HR1-ZHT. BJAB is an EBV-negative line derived from a tumor biopsy of a 5-year-old Kenyan female with BL (156). Namalwa is an EBV-positive line derived a young Kenyan female with BL (157) and is defective for lytic reactivation due to viral genome integration at two loci on chromosome 1. Raji is an EBV-positive line derived from an 11-year-old Nigerian male Nigerian with BL (158) that exhibits abortive lytic infection due to a mutation in *BALF2* that impairs viral genome replication. Mutu I is a type I latency EBV-positive BL line derived from a Kenyan patient (159). Daudi is a type I latency EBV-positive line derived from a 16-year-old Kenyan male with BL (160). P3HR1-ZHT is derived from a subclone of the Jijoye cell line engineered for hydroxytamoxifen-inducible lytic infection via transfection with a vector encoding *BZLF1* fused to the hormone binding domain of the murine estrogen receptor (161–163). IBL1 and Farage are EBV-positive non-Hodgkin derived cell lines. IBL1 is a type II/III latency line derived from an AIDS immunoblastic lymphoma of a 40-year-old white male patient diagnosed with AIDS and cutaneous Kaposi’s sarcoma (164). Farage is a type III latency line derived from a lymph node biopsy of a 70-year-old female patient diagnosed with diffuse large B cell non-Hodgkin lymphoma (165). All cell lines were cultured at 37°C with 5% CO_2_ on 48- or 96-well culture plates (Corning, 07-200-86) in RPMI 1640 medium (Gibco 11-875-119) supplemented with 10% Fetal bovine serum (Corning, MT35010CV).

### Lytic induction treatments

Cells in culture plates were treated with lytic inducing agents at the following concentrations: 5 ng/mL TGF-beta (R&D Systems, 7754-BH-005), 10 µg/mL goat anti-human IgG (Jackson ImmunoResearch, 109-005-170), 25 nM TPA (Phorbol-12-myristate-13-acetate) (Millipore Sigma, 52-440-05MG) + 3 mM Sodium Butyrate (Sigma-Aldrich, 567430), 300 mM Hydrogen Peroxide (Thermo-Scientific, H325-500), 100 nM Doxorubicin Hydrochloride (Biotechne, 2252/10), 100 nM Etoposide (Sigma-Aldrich, 341205-25MG), 50 nM Panobinostat (MedChemExpress, HY-10224R), 100 nM 4-Hydroxytamoxifen (4-HT) (MedChemExpress, 50-187-3523), 10 µM KU-55933 (Selleck Chem, S1092), 1 µM Phosphonoacetic acid (PAA) (Thermo Scientific Chemicals, AAA1211709). Cells were incubated in treatments for 36 hours in single-timepoint survey experiments. Cells in timecourse experiments were incubated in treatments for 1 hour, centrifuged at 350 x g for 5 mins to pellet and remove treatments, then resuspended in fresh R10 and incubated for 24, 48, or 72 hours prior to harvest.

### Live cell labeling

Treated cells were pulse-labeled with 10 μM of 5-ethynyl-2’-deoxyuridine (EdU) (Invitrogen, C10639) and incubated for 1 hour to assay DNA synthesis immediately prior to harvest. In separate experiments, cells were stained for mitochondria detection instead of EdU at this stage with MitoTracker Orange CMTMRos (Invitrogen, M7510). Following incubation with either EdU or MitoTracker, cells were centrifuged at 350 x g for 5 mins and resuspended in 1x Phosphate Buffered Saline (PBS) (Gibco, 10010049). In epifluorescence imaging experiments, cells were stained with viability dye (Thermo Fisher Fixable Live/Dead NIR Kit, L34975).

### Cell transfer to optical plates, fixation, and permeabilization

Treated cells labeled with EdU, MitoTracker, and/or viability reagents were transferred to 96-well optical plates (Revvity PhenoPlate, 6055302) coated with 10 ug/mL poly-D-lysine (Gibco, A3890401) and centrifuged at 350 x g for 5 minutes. Alternatively, glass-bottom plates (Greiner Bio-One CellView 96-well, 655891) were used for epifluorescence imaging experiments. Supernatants were removed and cells were fixed with 100 μL of 4% Paraformaldehyde (Electron Microscopy Sciences, 15710) in 1x PBS per well for 15 minutes. Fixative was removed and cells were washed twice with 3% BSA (Thermo Scientific, 37525) in 1x PBS. Cells were permeabilized with 100 μL of 0.5% Triton X-100 (Fisher, BP151-100) in 1x PBS for 20 minutes then washed twice with 3% BSA in 1x PBS. For EdU pulse-labeling experiments, permeabilized cells were stained for EdU detection with A594-azide according to the Invitrogen Click-It EdU kit manufacturer protocol (Invitrogen, C10639). Following EdU detection, cells were washed twice with 3% BSA in 1x PBS before proceeding with immunofluorescence steps.

### Immunofluorescence and DNA staining

Cells were either incubated specified pairs (one mouse, one rabbit) of the following primary antibodies: anti-BZLF1 mouse monoclonal IgG1κ (1:250 dilution, Santa Cruz Biosciences, sc-53904); anti-gp250/350 mouse monoclonal IgG1 (1:250 dilution, Santa Cruz Biosciences, sc-57724); anti-γH2AX (phospho-S139) rabbit monoclonal IgG (1:250 dilution, Abcam, ab81299); anti-53BP1 rabbit monoclonal IgG (1:250 dilution, Abcam, ab175933); anti-TOM20 rabbit polyclonal IgG (1:200 dilution, ProteinTech, 11802-1-AP); and anti-Ki67 rabbit polyclonal IgG (1:200 dilution, Abcam, ab15580). All primary antibody cocktails were prepared in 0.1% BSA in 1x PBS. Primary antibodies were incubated on samples overnight at 4°C and washed twice with 3% BSA in 1x PBS. Secondary antibody cocktails were prepared in 0.1% BSA in 1x PBS with 1:500 goat anti-mouse IgG H&L AlexaFluor 488 (Abcam, ab150113) and 1:250 goat anti-rabbit IgG H&L AlexaFluor 647 (Abcam, ab150079) and incubated in wells for one hour at room temperature. Cells were washed twice with 3% BSA in 1X PBS before nuclear counterstaining with Hoechst 33342 in 1x PBS for 10 minutes. Excess Hoechst was removed by one wash with 1x PBS, and wells were topped to 200 μL of 1x PBS for imaging.

### Microscopy and image processing

Confocal images were acquired on a Yokogawa CellVoyager CV8000 High-Content Screening system (Yokogawa Electric Corporation, Tokyo, Japan) with four excitation lasers (405 nm, 488 nm, 561 nm, and 640 nm), a 60x water immersion objective (NA = 1.2; UPLSAPO60XW), and a 25 μm pinhole dual microlens spinning disk confocal unit. The pixel size in acquired images was 0.1075 μm/pixel. Four fluorescence and one brightfield channels were acquired in all experiments with exposure times ranging from 100-500 ms per channel. Five to six z-planes with 2 μm spacing were captured for each of thirty-two fields of view (FOV) per well in all experiments. Nuclei, cell, and cytoplasm masks in all FOVs were segmented, and feature extraction was performed using CellProfiler 4.2.6, where nuclear segmentation was performed using the “IdentifyPrimaryObjects” module with Otsu two-class thresholding applied to log-transformed Hoechst 33342 images. Cell boundaries were identified using the “IdentifySecondaryObjects” module applied to a sum of all images with nuclei serving as seed objects, and cytoplasmic regions were defined by subtracting nuclear areas from whole-cell areas using the “IdentifyTertiaryObjects” module. A feature set of ~800 features was extracted for each cell, consisting of intensity measurements, intensity co-localization features, texture features, and morphometric features for nuclear, cytoplasmic, and cellular compartments.

Widefield epifluorescence images were acquired on a Leica DMi8 THUNDER Cell imaging system (Leica Microsystems, Wetzlar, Germany) equipped with an eight-line LED light source (PE 800), two quad bandpass cubes (DAPI/FITC/TRITC/CY5 and CFP/YFP/RFP/NIR configurations), an external emission bandpass wheel, and 4.2 megapixel (2048 x 2048) scientific CMOS sensor (Leica K8 camera). Six-channel images (five fluorescence, one brightfield) were acquired using a 40x air objective (Leica #11506414, HC PL APO 40x/0.95 CORR). Widefield images were processed in real-time with THUNDER instant computational clearing (ICC; Leica) to reduce out-of-plane light for image enhancement. Single cells from confocal and ICC epifluorescence images were segmented and measured using CellProfiler (53). Example CellProfiler pipelines used for this study are included as supplementary files (**Files S2-S3**).

### Data visualization and analysis

Single-cell feature matrices containing approximately 300,000 cell observations per 96-well plate were processed using PyCytominer (version 0.2.0) with data normalization performed using the median absolute deviation (MAD) robustize method (mad_robustize), which normalizes features by subtracting the median and dividing by the median absolute deviation to provide robust standardization resistant to outliers. Quality control procedures included removal of wells with insufficient cell counts (< 100 cells per well), identification and removal of outlier wells based on feature distributions, and assessment of plate effects with batch corrections where necessary. Low-variance features were identified and removed using a variance threshold of 0.01 to eliminate uninformative measurements, and the resulting filtered feature matrix was subjected to dimensionality reduction using Uniform Manifold Approximation and Projection (UMAP) as implemented in umap-learn (version 0.5.3) with default parameters (n_neighbors=15, min_dist=0.1, n_components=2) to generate a two-dimensional representation of the high-dimensional feature space. Unsupervised clustering was performed on the UMAP-embedded data using the Leiden algorithm as implemented in the leidenalg package, with clustering resolution optimized by testing multiple values and selecting the resolution that maximized modularity while maintaining biologically interpretable cluster sizes. All analyses were performed in Python 3.8+ using standard scientific computing libraries including NumPy, Pandas, SciPy, and scikit-learn, with statistical significance assessed using appropriate non-parametric tests (Mann-Whitney U test for pairwise comparisons, Kruskal-Wallis test for multiple group comparisons) given the typically non-normal distribution of morphologic features, and multiple testing correction applied using the Benjamini-Hochberg false discovery rate (FDR) method where appropriate.

Measurements were loaded into R Studio and adapted for morphologic analysis and visualization with functions from Seurat v5 (55). Measurement .csv files including sample metadata were prepared as Seurat objects. Measurements were processed using modified normalization, scaling, and dimensional reduction steps for epifluorescence datasets or as described above for confocal imaging experiments (see **File S1**). Morphologic feature visualization and differential feature analysis were achieved with standard Seurat functions (e.g., *FindMarkers()*, *DimPlot()*, *FeaturePlot()*, *VlnPlot()*, *FeatureScatter()*). Pseudotime trajectories were calculated using a Seurat wrapper for monocle3 (72). Data for individual cells in UMAP plots were interactively accessed using the *HoverLocator()* function enabled by the plotly R package (166), which enabled recovery of cell images through file metadata and xy bounding box coordinates. Single-channel and merged images were processed with Fiji (167) for visualization.

### Statistical analyses

Morphologic features with statistically significant differences among cell clusters were identified by Wilcoxon rank-sum testing with Bonferroni correction for multiple hypothesis testing. Significant differences in quantitative feature distributions stratified by treatment condition were determined by Kolmogorov-Smirnov (KS) tests. Significant associations between DNA damage patterns (e.g., pan-nuclear γH2AX, 53BP1 foci) and lytic status (Zta expression) were evaluated by chi-squared tests.

## Data and Code Availability

Images and single-cell quantification data will be made available upon reasonable request. Code for loading, processing, and analyzing CellProfiler output measurement files (.csv) with Seurat and Monocle3 in R is included as a supplementary file. Please direct relevant requests to the corresponding author.

## Acknowledgments

We wish to thank Tracey Schultz and Craig Dobry for lab management activities that supported the work herein. We also wish to thank Jesse Wotring and Matthew McConnachie for instrument support and Benjamin Halligan for help with alternative data analysis pipelines not presented in the study. Special thanks are in order for University of Michigan Department of Microbiology and Immunology staff members, particularly Brenda Franklin and Stephanie Himpsl. Finally, we wish to thank colleagues in the Department, especially Drs. Katherine C. Barnett, Christiane E. Wobus, and Bethany B. Moore for thoughtful feedback on the manuscript.

## Author Contributions

D.T. contributed to the study design, performed experiments, co-developed cell segmentation pipelines, contributed to data interpretation and analysis, co-wrote the original manuscript draft, and provided edits for the final version. J.Z.S. provided technical support for confocal imaging and data processing, co-developed cell segmentation pipelines, wrote methods for the original manuscript draft, and provided edits for the final version. E.D.S. designed the study, performed experiments, co-developed cell segmentation pipelines, developed cluster analysis and pseudotime software implementations, analyzed and interpreted the data, co-wrote the original manuscript draft, and prepared the final version.

## Ethics Statement

All experiments in this study were conducted with Institutional Biosafety Committee (IBC) approval from the University of Michigan (#IBCA00002858).

## Supporting Figure Legends

**Figure S1**. **Imaging and treatment controls for HCS of EBV lytic reactivation.**

**A)** Fluorescence detection controls. 1^st^ row: cells stained with Hoechst only. 2^nd^ row: cells stained with Hoechst and EdU-A594. 3^rd^ row: cells stained with Hoechst and incubated with fluorophore-conjugated secondary antibodies in the absence of primary antibodies. 4^th^ row: cells with Hoechst staining and anti-gp350 detection via A488-conjugated secondary antibody. 5^th^ row: as in row 4, plus anti-53BP1 detection via A647-conjugated secondary antibody. 6^th^ row: full 4-color staining panel to detect nuclei, gp350, EdU, and 53BP1. 7^th^ row: as in row 6, with nuclei, Zta, EdU, and γH2AX detection. Intensity scaling within each channel (column) is uniform with the exception of higher max intensity for γH2AX detection (647 nm laser). The same A488-conjugated secondary was used to detect viral proteins, and the same A647-conjugated antibody was used to detect host targets.

**B)** Lytic reactivation and DNA damage control conditions. Integrated Zta intensity distributions for segmented cells from 4HT-induced versus unstimulated P3HR1-ZHT (top panel). Integrated γH2AX intensity distributions for segmented cells induced with 4HT, pre-treated with KU-55933 to impair pATM activity prior to 4HT induction, and unstimulated conditions (bottom panel).

**C)** Quantification of channel crosstalk controls. EdU-A594 intensity in segmented cells from specified conditions (top panel). Anti-gp350 intensity in segmented cells from specified conditions (middle panel). Anti-53BP1 intensity in segmented cells from specified conditions (bottom panel).

**Figure S2. Confocal HCS experimental data overview.**

**A)** UMAP of segmented cells coded by phenotypic cluster, treatment condition, cell line, and calculated pseudotime for DNA_EdU_Zta_γH2AX survey experiment.

**B)** As in A, for DNA_EdU_gp350_53BP1 survey experiment

**C)** As in A, with cell coding by time post-treatment instead of pseudotime for DNA_EdU_Zta_53BP1 experiment.

**D)** As in C, for DNA_EdU_gp350_γH2AX experiment.

**E)** As in A (no time / pseudotime calculation), for DNA_Mito_Zta experiment.

**F)** As in E, for DNA_Mito_gp350 experiment.

**Figure S3. Interactive exploration of quantitative morphology and recovery of corresponding cell images.**

**Figure S4. Treatment effects on DNA synthesis (1-hour EdU pulse immediately pre-harvest).**

**A)** Treatment-stratified per-cell EdU intensity distributions from treatment survey experiments. Measurements of integrated intensity are log-normalized. Asterisks denote statistically significant differences in intensity distributions (Kolmogorov-Smirnov test; ***p<0.001).

**B)** Treatment- and time-stratified per-cell EdU intensity distributions from temporal response experiments. Measurements of integrated intensity are log-normalized.

**Figure S5. Immunofluorescence signal quantification in Zta-positive cluster by treatment.**

**A)** Log-normalized integrated Zta intensity distributions for cluster 8 cells by treatment. Data are from DNA_EdU_Zta_γH2AX survey experiment. Asterisks denote statistically significant differences in Zta intensity distributions (Kolmogorov-Smirnov test; **p<0.01; ***p<0.001).

**B)** Zta signal entropy distributions in segmented nuclei for cluster 8 cells by treatment. Data are from DNA_EdU_Zta_γH2AX survey experiment. Asterisks denote statistically significant differences in Zta intensity distributions (Kolmogorov-Smirnov test; *p<0.05; ***p<0.001).

**Figure S6. Additional examples of genotoxin-induced γH2AX patterns in EBV-positive B cell lymphoma models.**

**A)** Merge and individual channel images for Zta-positive cells from doxorubicin treatment. Nuclei depicted in blue, Zta in green, EdU in yellow, and γH2AX in red for all panels.

**B)** Merge and individual channel images for Zta-positive P3HR1-ZHT cells across treatments. Channels are colored as in A.

**C)** Merge and individual channel images for Zta-positive Farage cells from panobinostat treatment. Channels are colored as in A and B.

**D)** Merge and individual channels of IBL cells from panobinostat treatment. Zta-positive cells not detected.

**E)** Detail of linear DNA damage pattern in panobinostat-treated IBL1 cells (from D).

**Figure S7. Association between Zta-positive cells and pan-nuclear γH2AX staining in response to genotoxins.**

**A)** Observed (left) and expected (right) numbers of cells by Zta and γH2AX status for doxorubicin and etoposide treatments (* n = 34 cells; cell derived from multiple cell lines). Result of chi-squared test is denoted.

**B)** As depicted in A, for panobinostat-treated cells (**n = 195 across P3HR1-ZHT cells). Result of chi-squared test is denoted.

**Figure S8. Cell treatment response over timecourse experiment: DNA, EdU, Zta, and 53BP1.**

**A)** Representative full-field 60x images from unstimulated cells (Namalwa, P3HR1-ZHT, and Mutu).

**B)** As in A, for cells treated with TGF-β.

**C)** As in A-B, for cells treated with TPA + NaB.

**D)** As in A, for cells treated with H_2_O_2_.

**Figure S9. Representative images of cell treatment response over timecourse experiment: DNA, EdU, gp350, and γH2AX.**

**A)** Representative full-field 60x images from unstimulated cells (Namalwa, P3HR1-ZHT, and Mutu).

**B)** As in A, for cells treated with TGF-β.

**C)** As in A-B, for cells treated with TPA + NaB.

**D)** As in A, for cells treated with H_2_O_2_.

**Figure S10. Log-transformed integrated signal intensity for fluorescence markers by treatment and phenotypic cluster.**

**A)** Violin and feature scatter plots of feature levels and correlation color-coded by cluster: DNA_Zta_EdU_53BP1 staining.

**B)** As in B, for DNA_EdU_gp350_γH2AX staining.

**Figure S11. Additional examples of individual Mutu cells treated with TGF-β.**

**A)** Example cells from DNA_EdU_Zta_53BP1 staining experiment (note: distinct EdU staining patterns and overlap with Zta delineate likely cellular versus viral DNA replication.

**B)** Single and multichannel images of DNA_EdU_gp350_γH2AX cells from sequential z planes.

**Figure S12. Additional signal line profiles in Mutu cells demonstrating inverse spatial patterns of viral DNA replication and host DNA damage markers.**

**A)** Mutu cell treated with TGF-β and stained for DNA, Zta, Edu, and 53BP1.

**B)** As in A, with cell staining for DNA, EdU, gp350, and γH2AX.

**Figure S13. Association between Zta-positive cells and 53BP1 foci in cells treated with TGF-β.**

**A)** Observed (left) and expected (right) numbers of cells by Zta and presence of 53BP1 puncta for TGF-β treated P3HR1 cells (* n = 107 manually evaluated cells). Result of chi-squared test is denoted.

**B)** As depicted in A, for Mutu cells (**n = 125 manually evaluated cells). Result of chi-squared test is denoted.

**Figure S14. Pseudotime analysis of Mutu cells treated with TGF-β and stained for DNA, EdU, Zta, and 53BP1.** Callout lines denote the approximate UMAP location of each depicted cell.

**Figure S15. Epifluorescence image deconvolution and single-cell segmentation.**

**A)** Detail of cell from original six-channel epifluorescence image. DNA in blue, Zta in green, mitochondrial dye in orange, Ki-67 in red, near infrared viability dye in yellow, and brightfield in gray. Image was acquired using a 40x air objective with 0.95 NA.

**B)** Detail of images in A after computational deconvolution using Leica THUNDER instant computational clearing (ICC) method. Colors are denoted as in A.

**C)** Representative cell segmentation for ICC epifluorescence images. From left to right: composite channel signal (excluding bright field), denoised image, size-filtered single-cell masks from watershed segmentation, and overlay of masked signal into brightfield image.

