## Supplementary figures and images for "Reconstructing EBV reactivation and DNA damage response kinetics in morphologic pseudotime"

### Figure S1

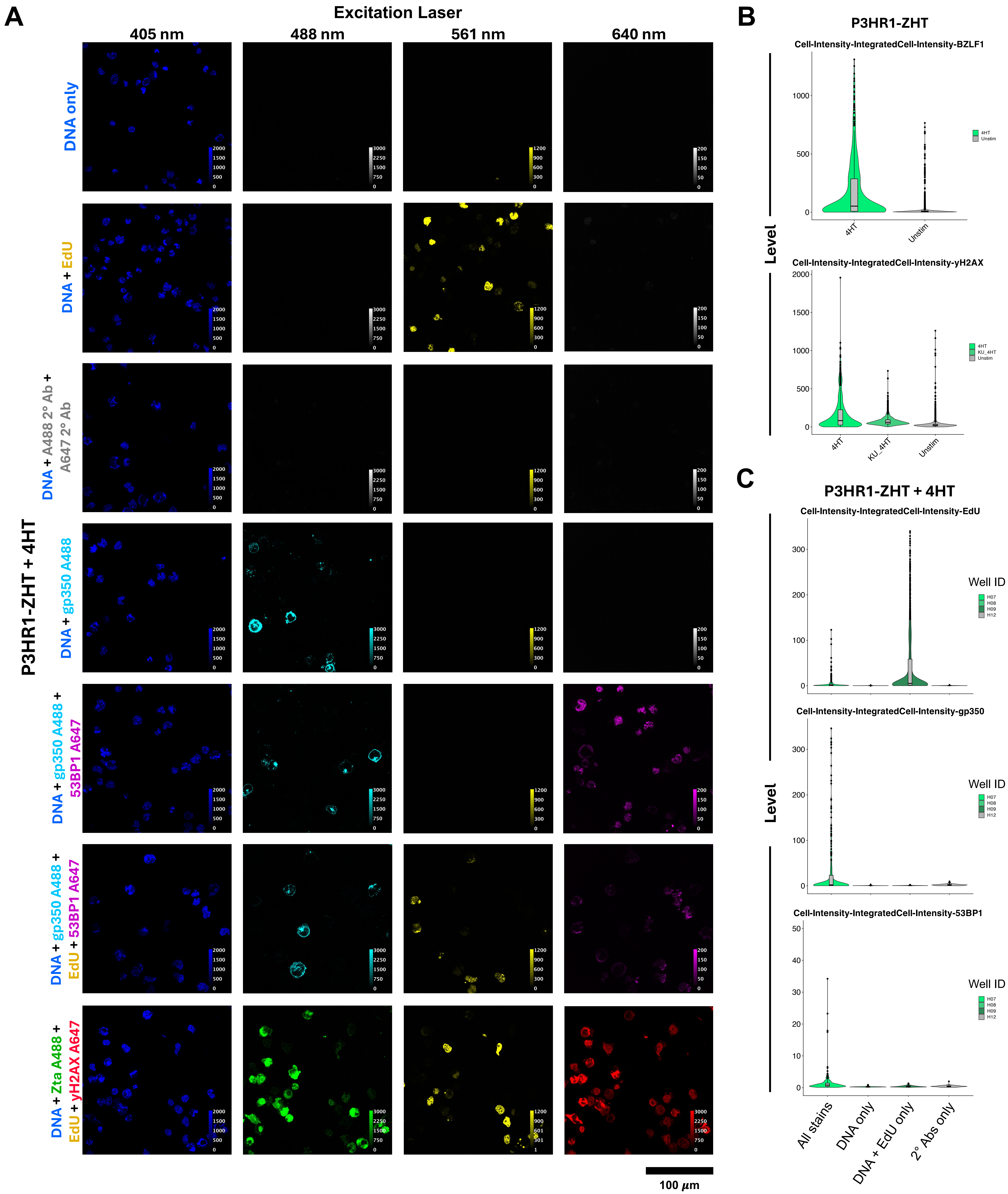

### Figure S2

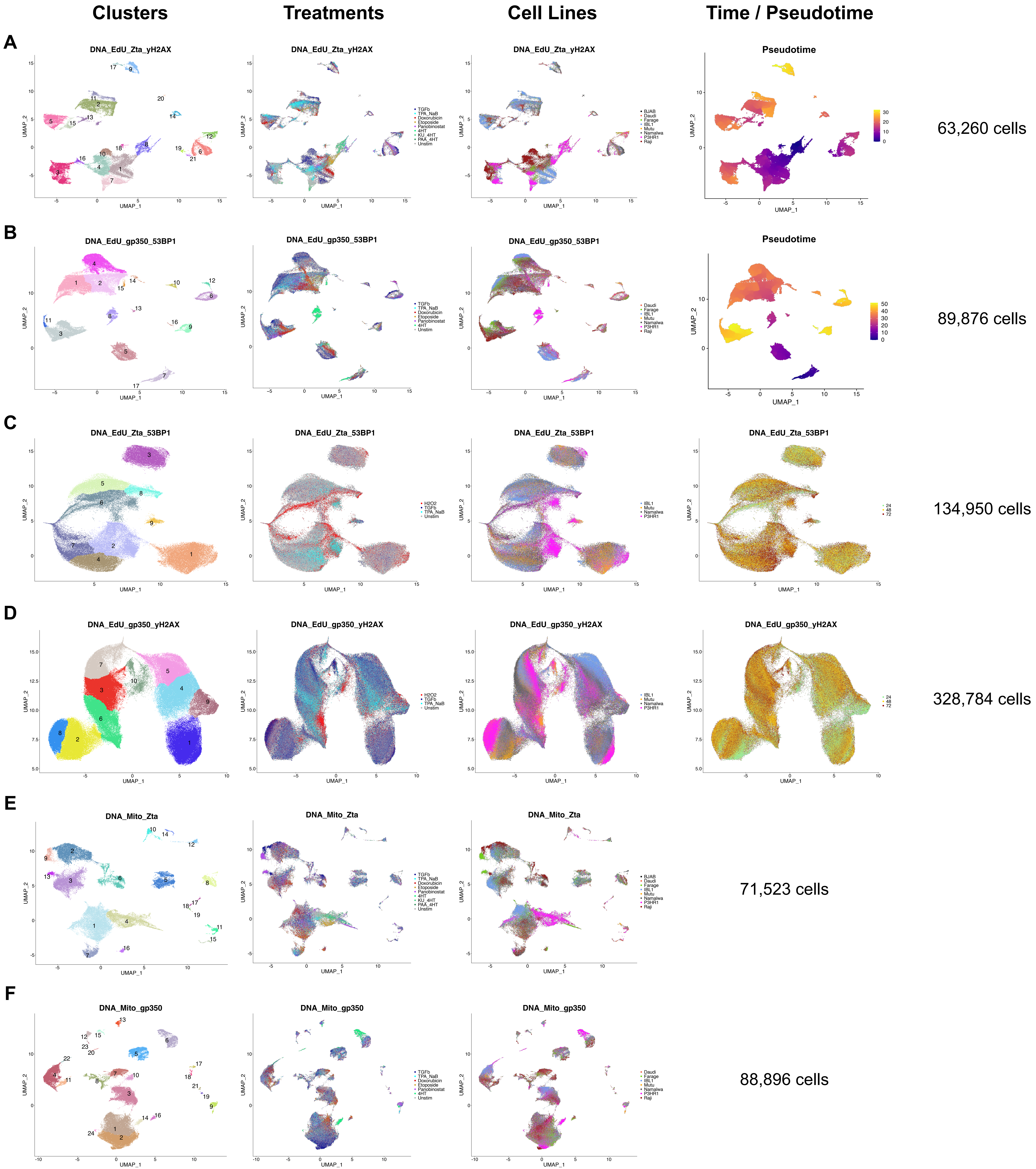

### Figure S3

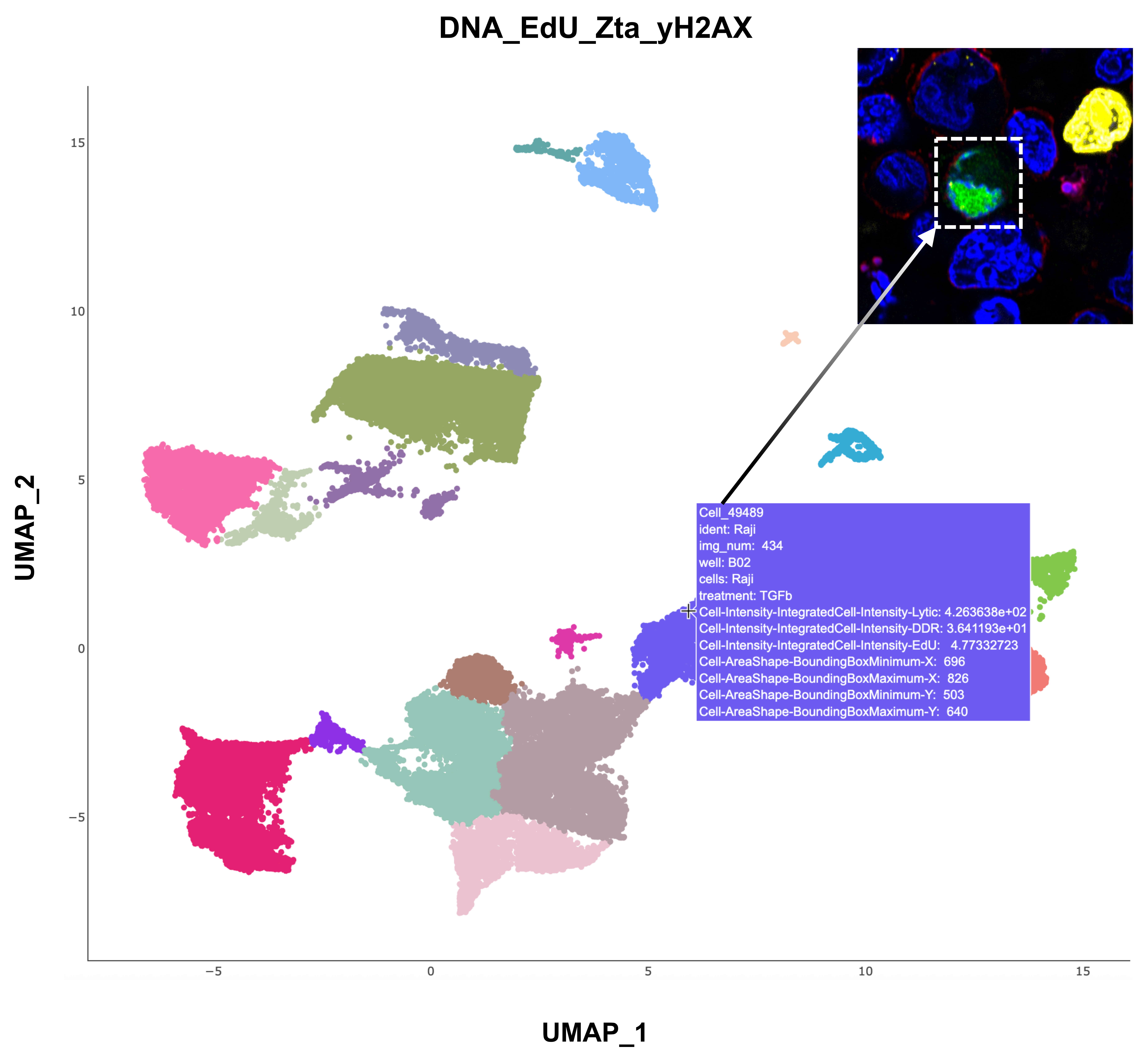

### Figure S4

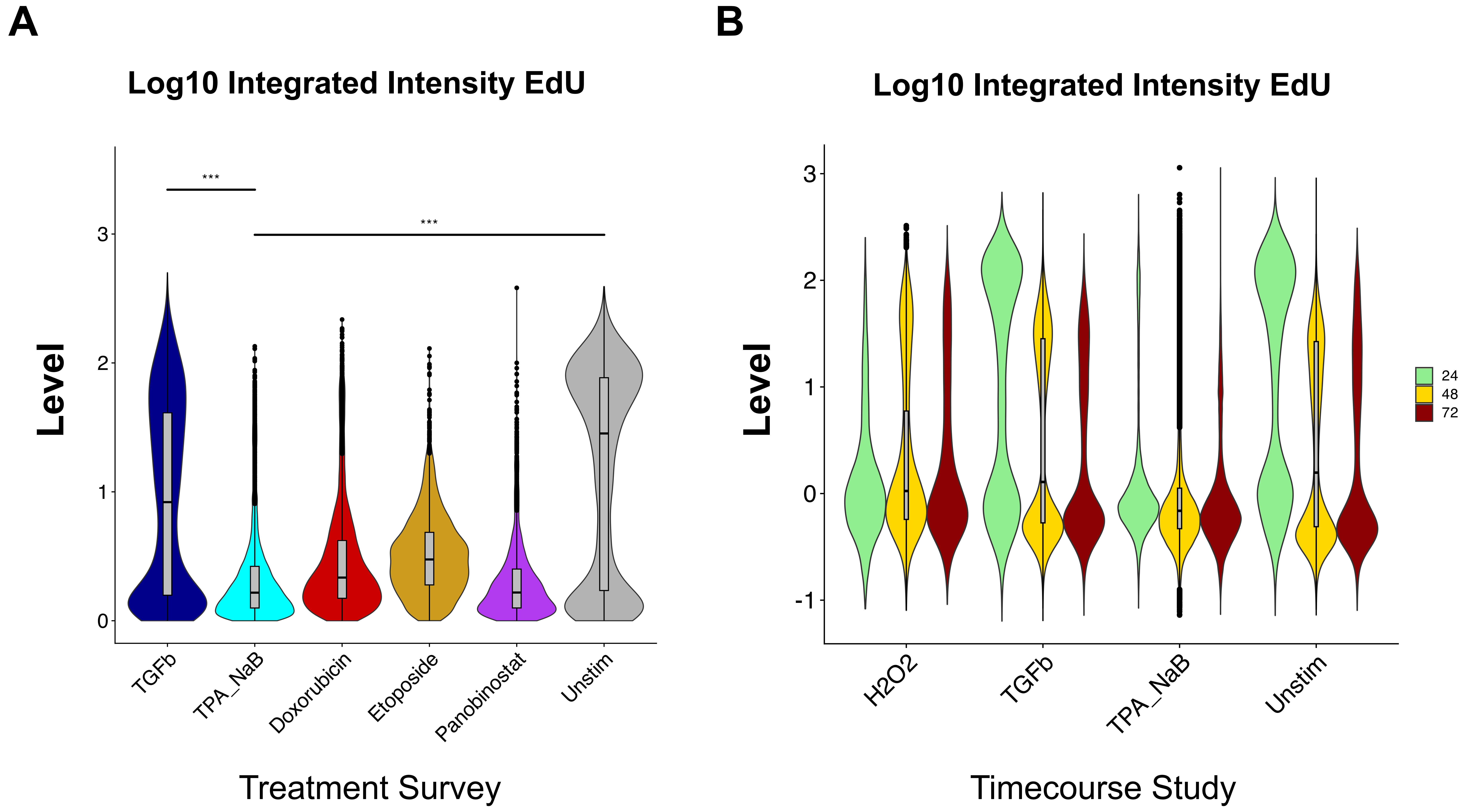

### Figure S5

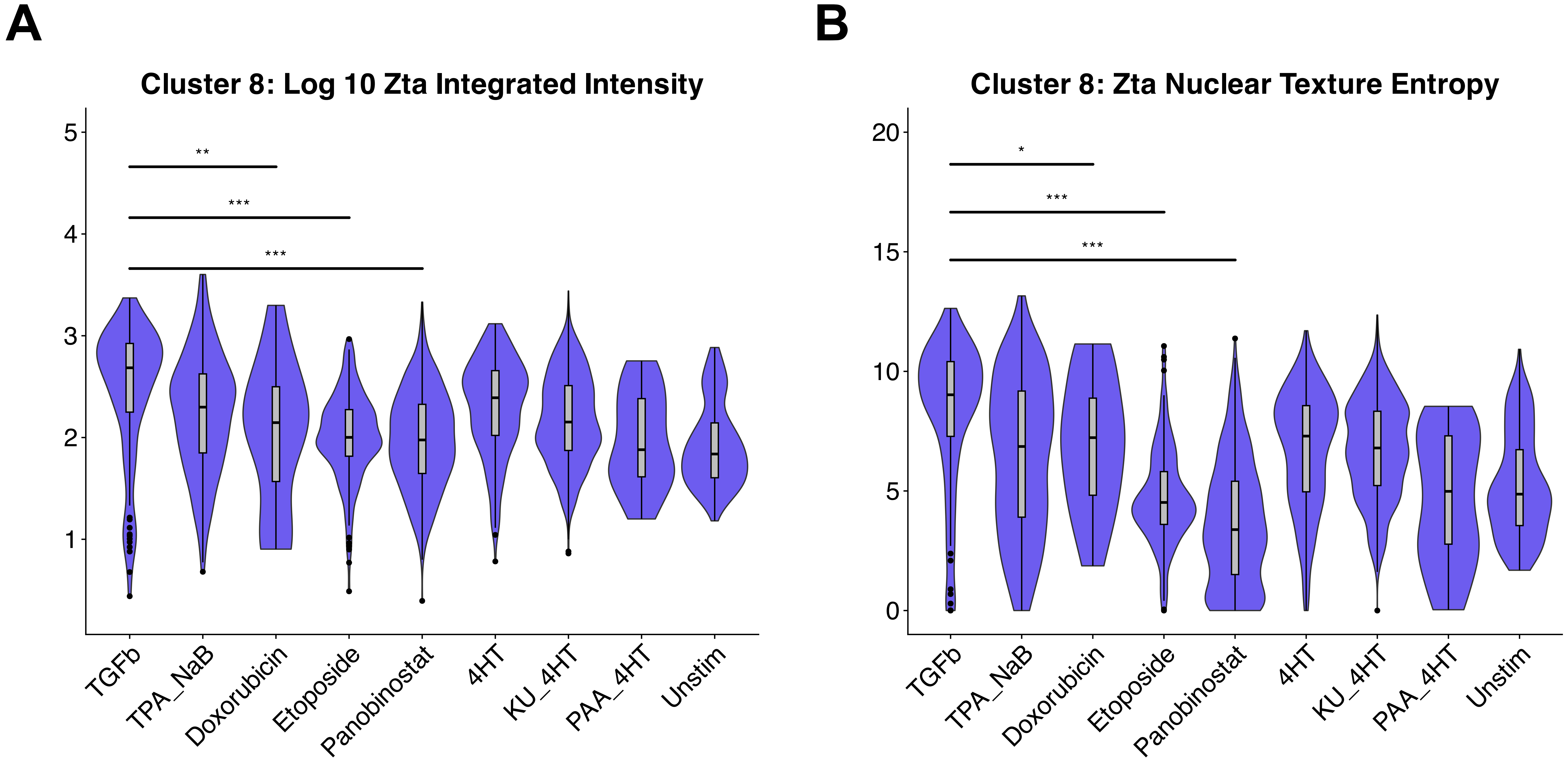

### Figure S6

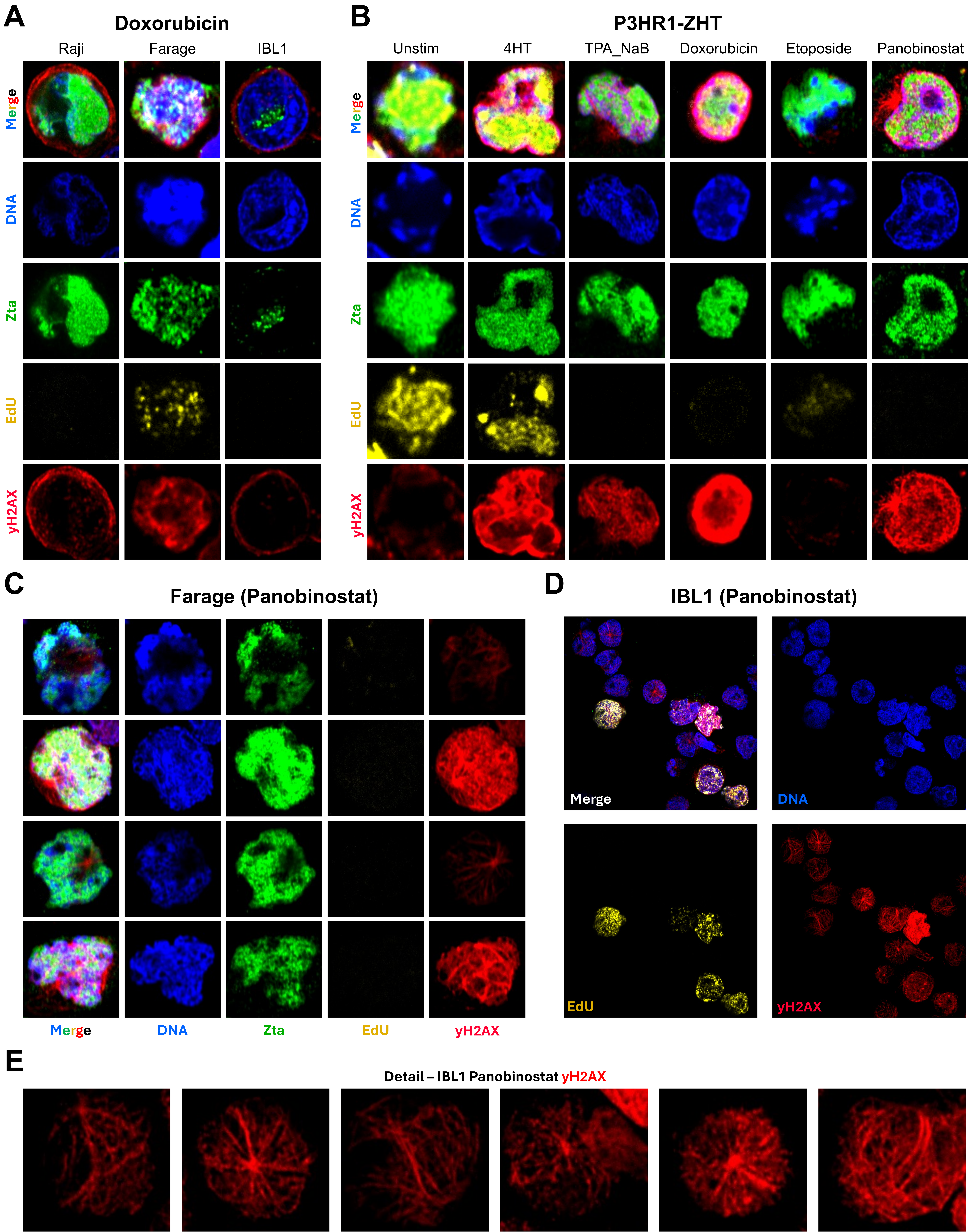

### Figure S8

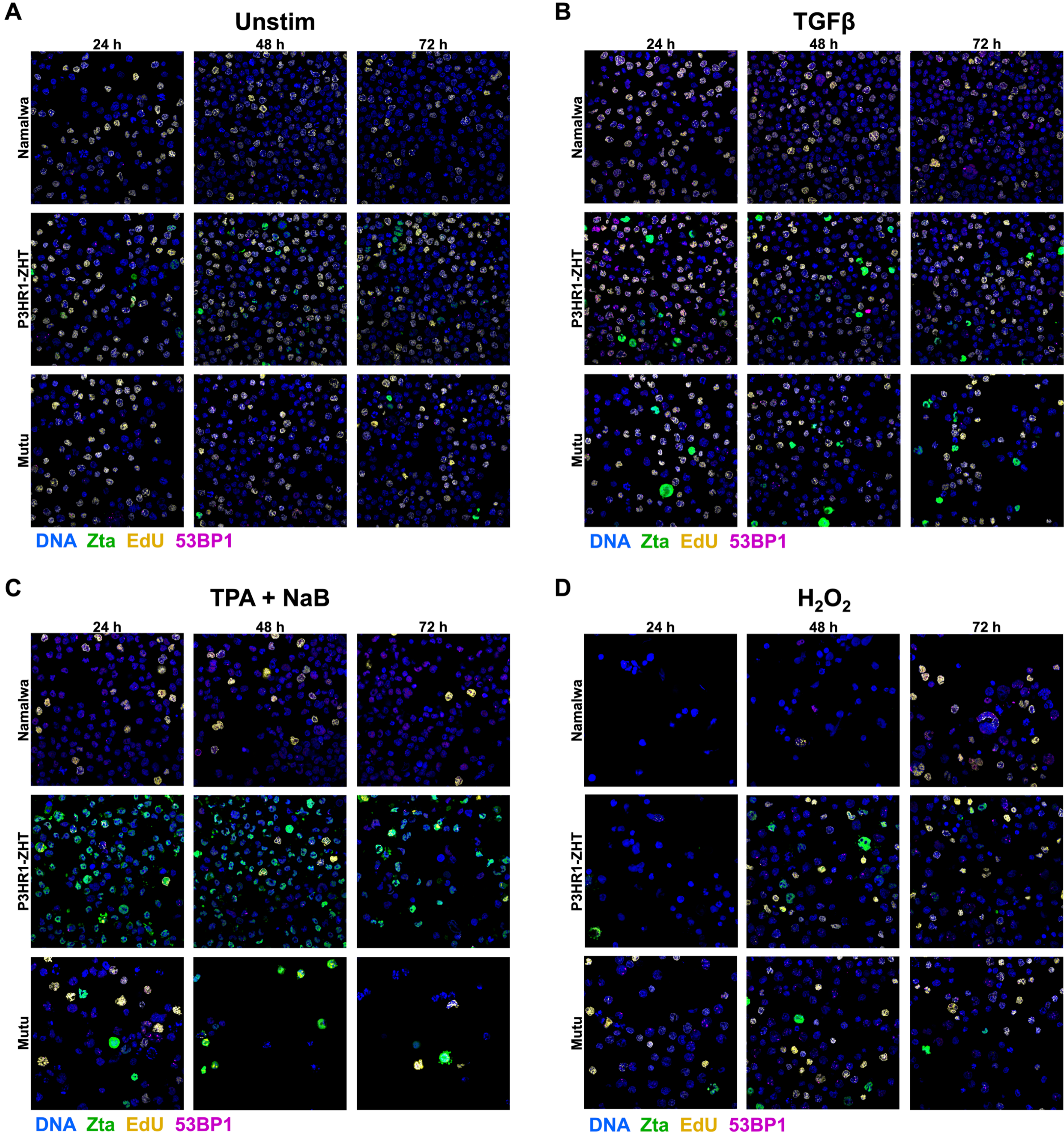

### Figure S9

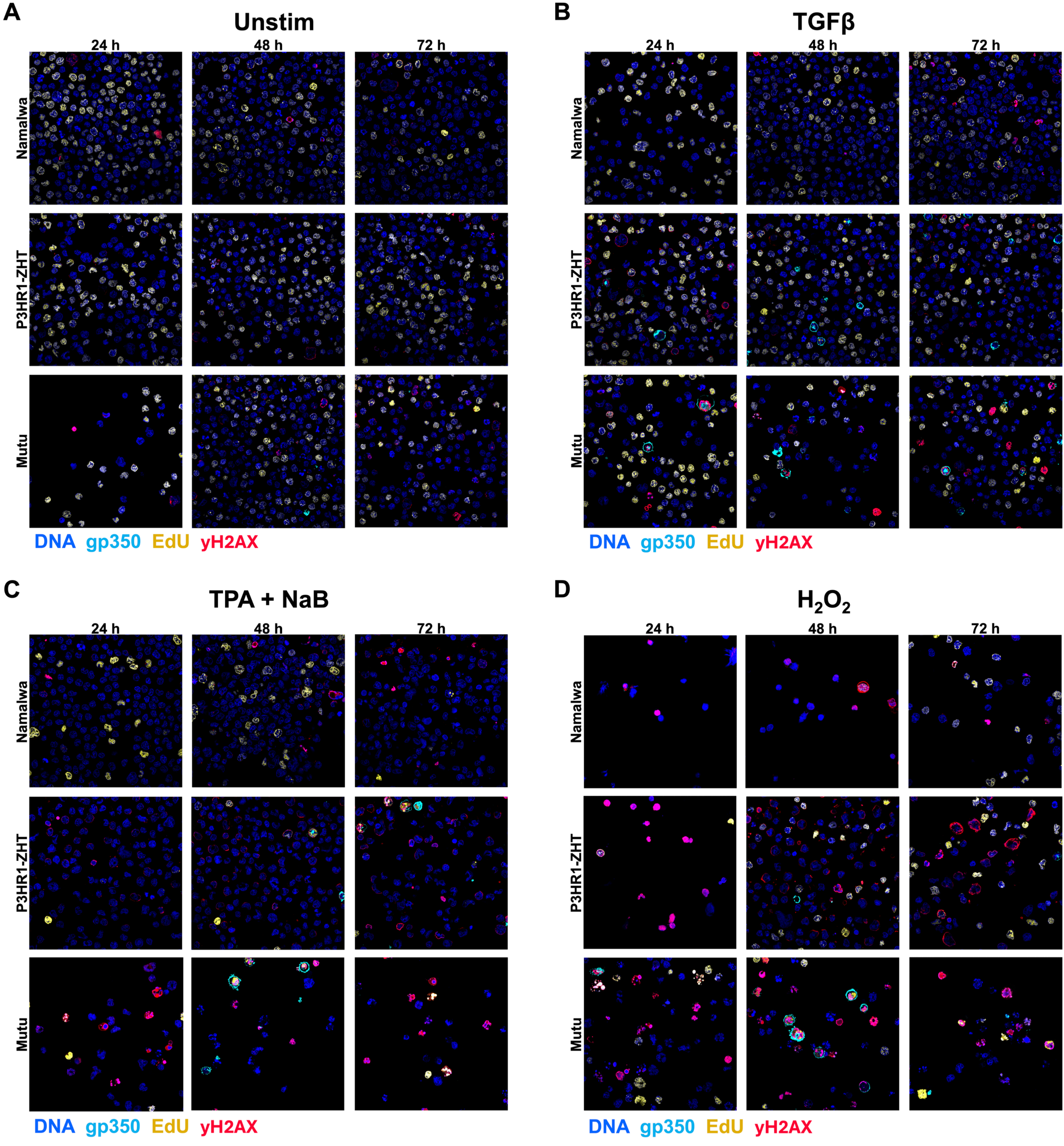

### Figure S10

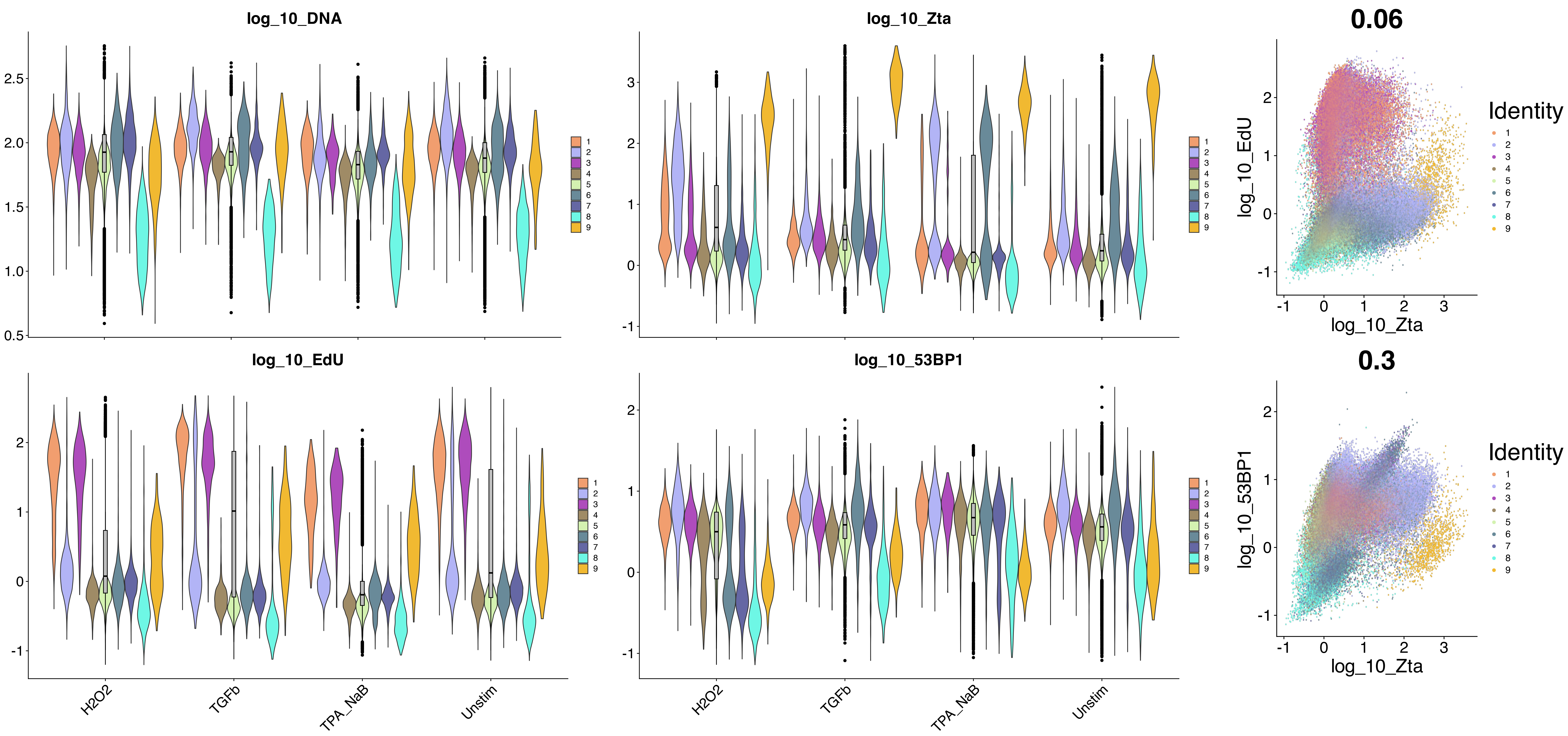

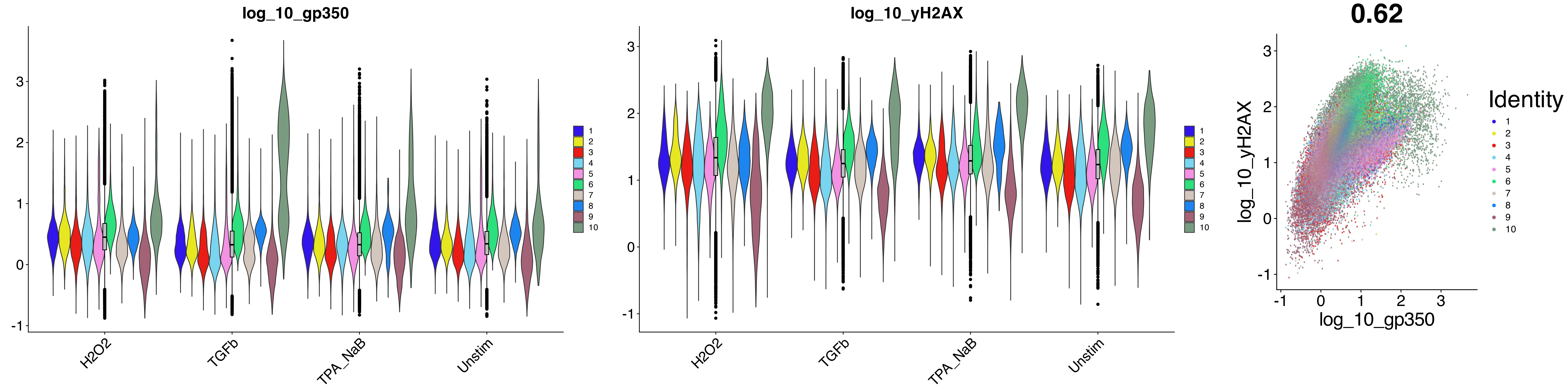

### Figure S11

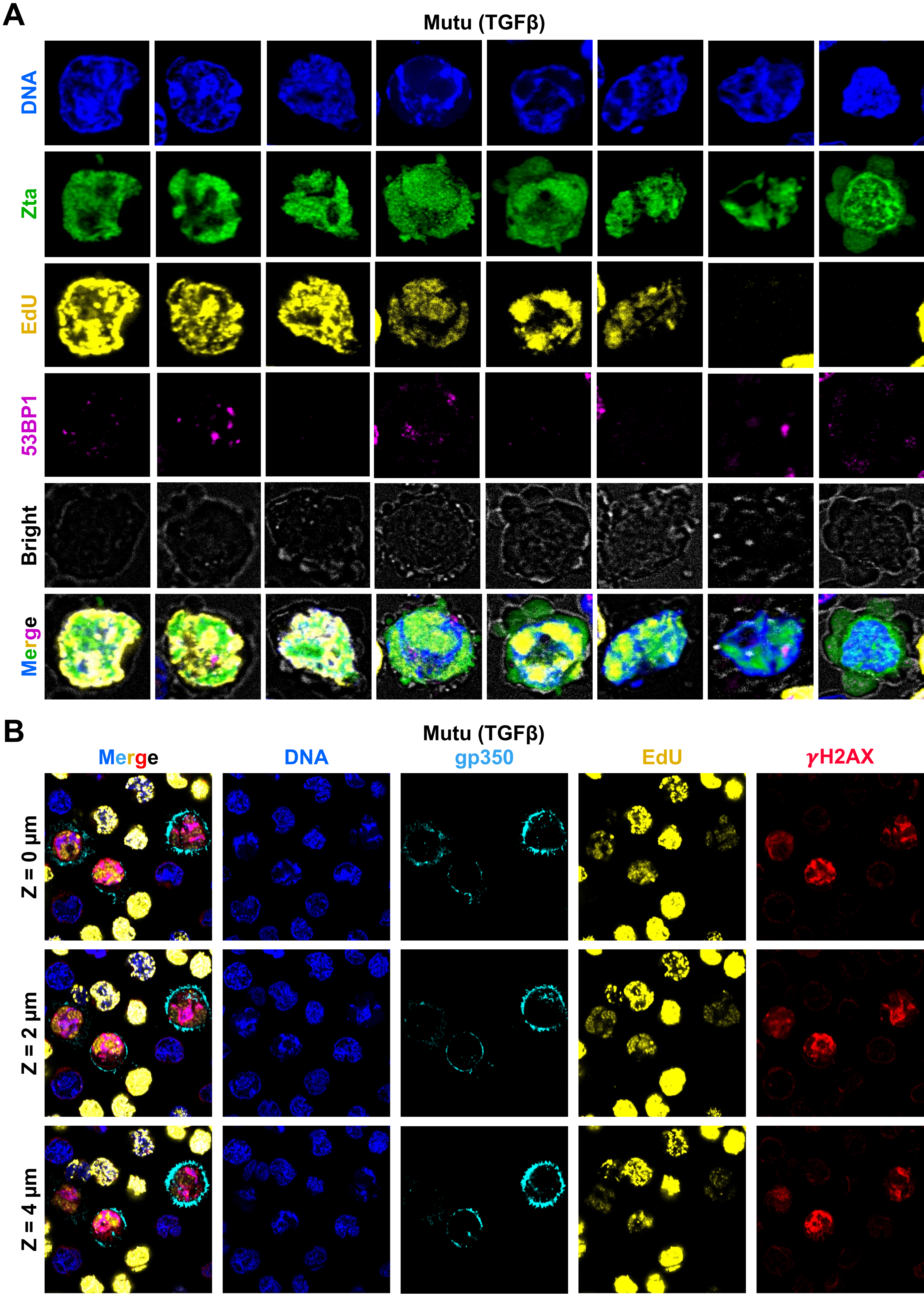

### Figure S12

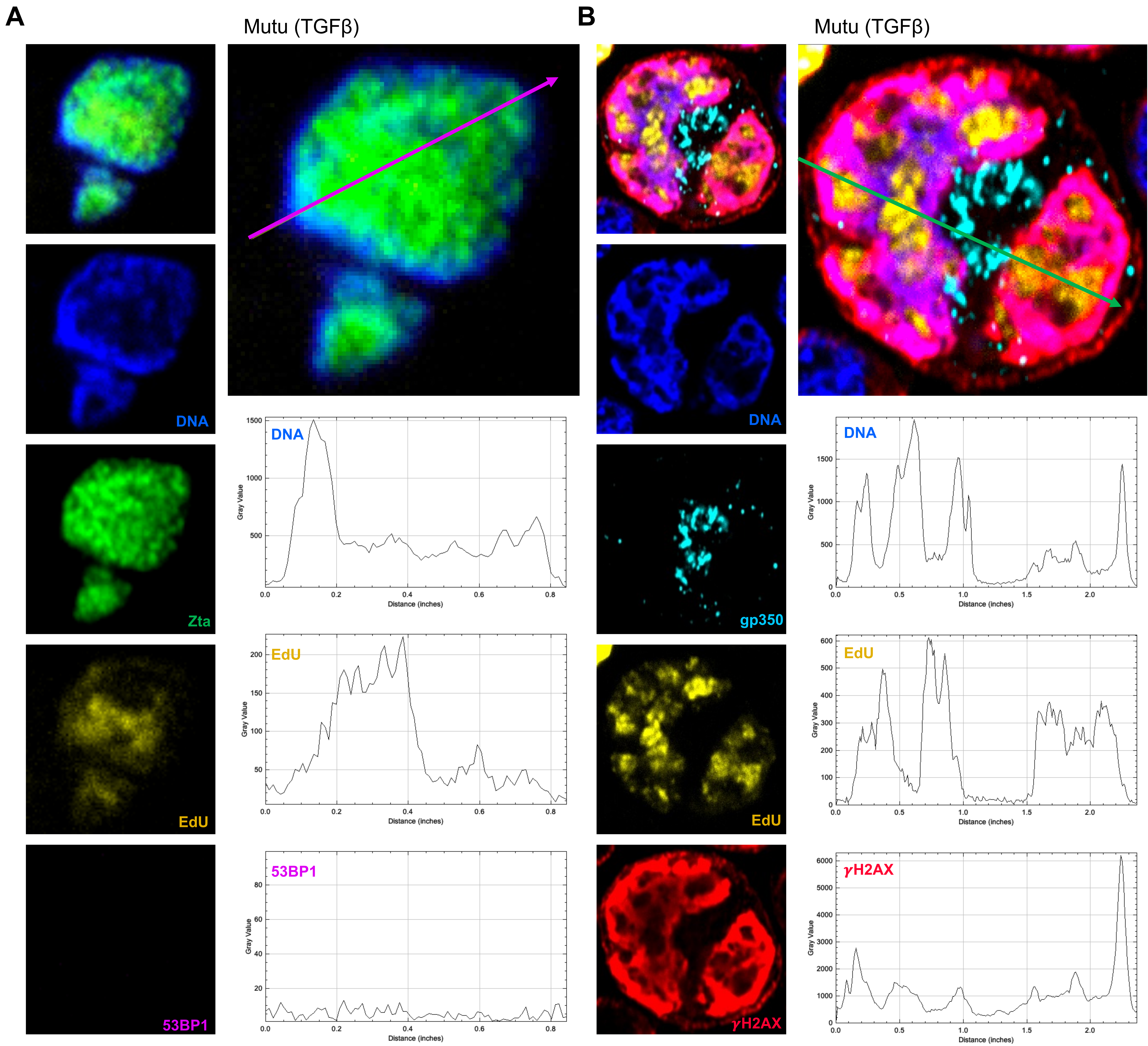

### Figure S14

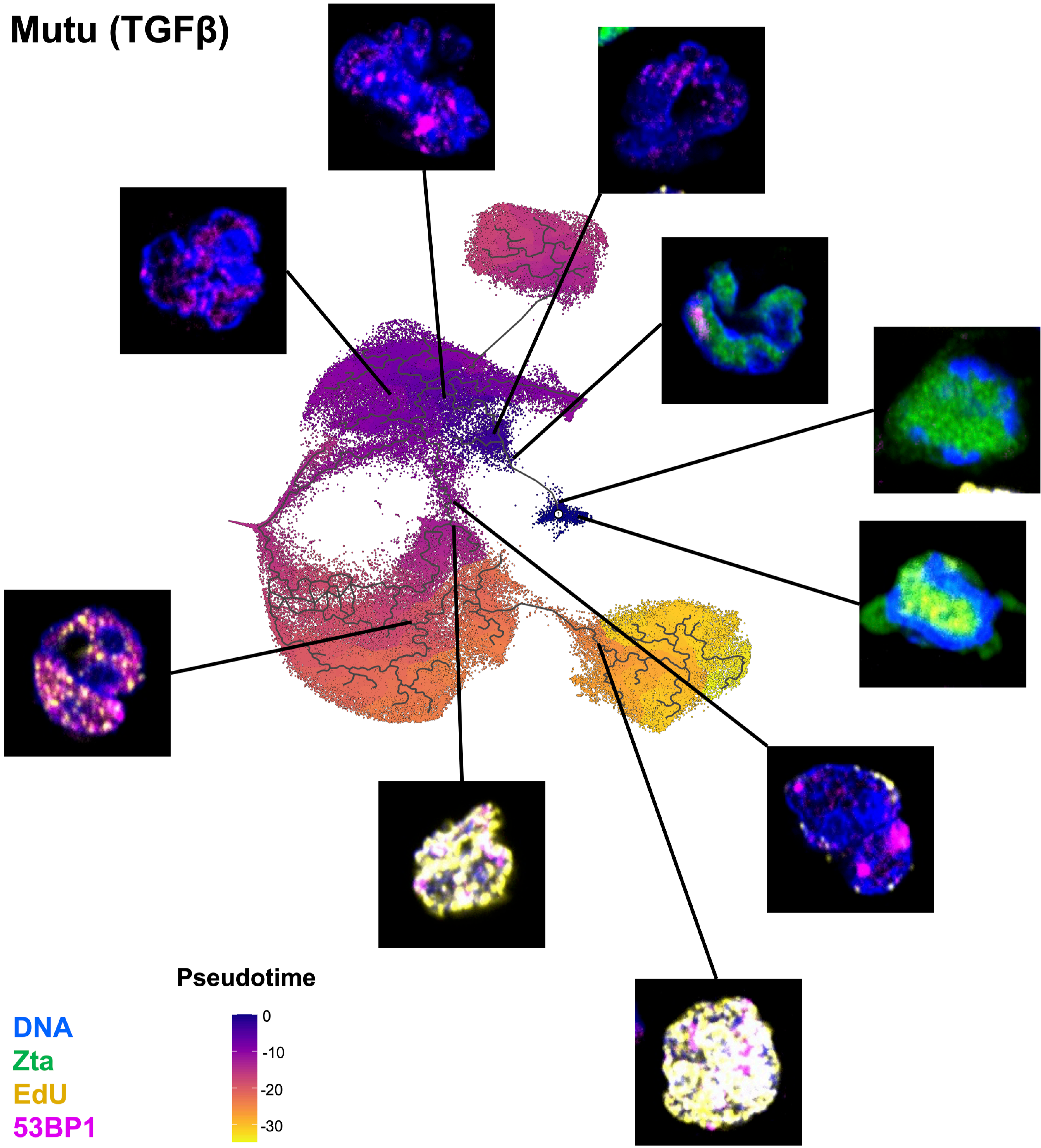

### Figure S15

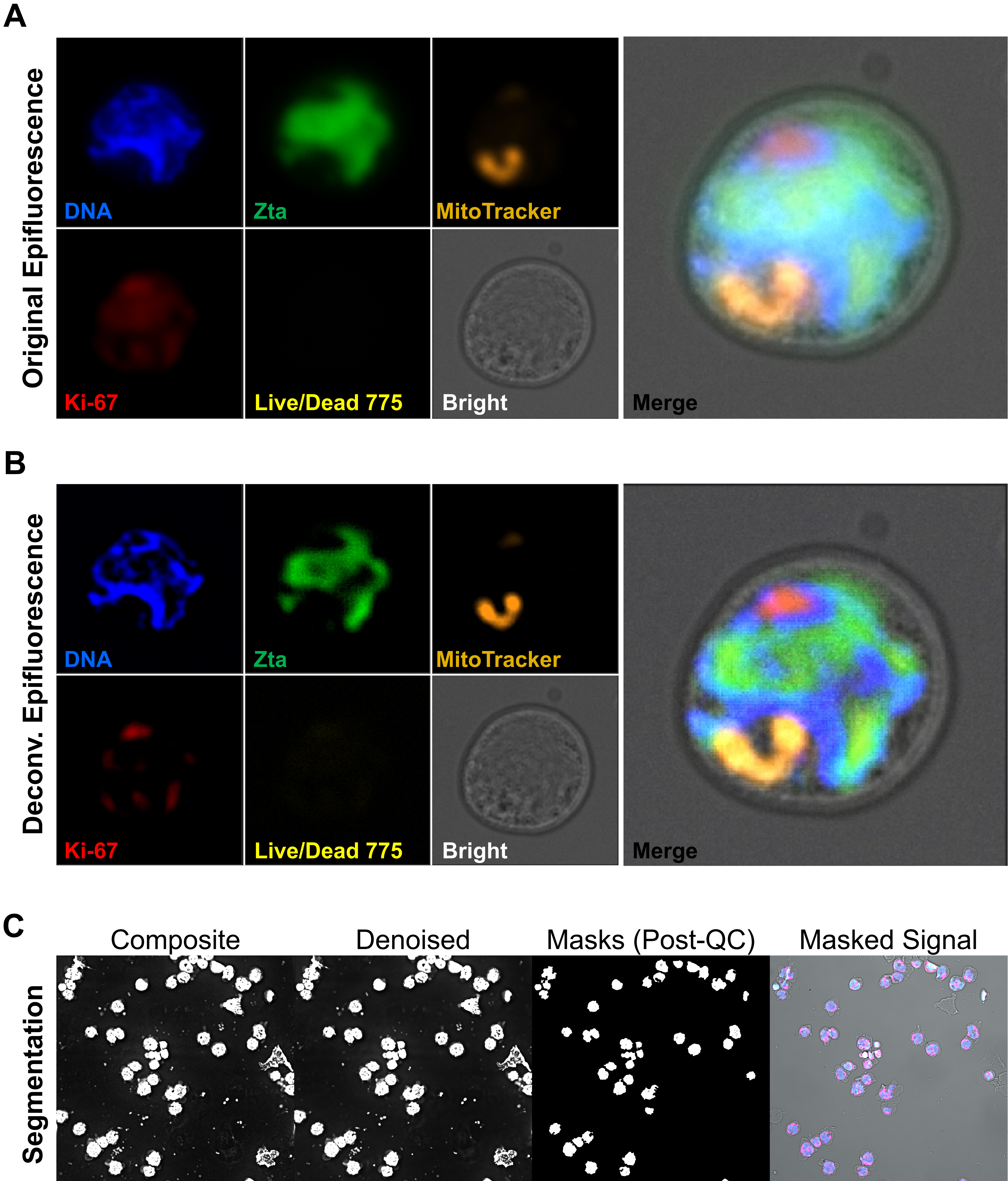
