## Supplementary material for "Reconstructing EBV reactivation and DNA damage response kinetics in morphologic pseudotime": Figure S7

**A**

| Dox & Etop (n=34)* | Zta+ | Zta- | Total |
|---|---|---|---|
| Pan-nuclear $\gamma$H2AX | 17 | 4 | 21 |
| Other $\gamma$H2AX | 2 | 11 | 13 |
| Total | 19 | 15 | 34 |

| Expected | Zta+ | Zta- | Total |
|---|---|---|---|
| Pan-nuclear $\gamma$H2AX | 11.7 | 9.3 | 21 |
| Other $\gamma$H2AX | 7.3 | 5.7 | 13 |
| Total | 19 | 15 | 34 |

**Chi-squared test p = 1.83e-4**

**B**

| Panobinostat (n=195)** | Zta+ | Zta- | Total |
|---|---|---|---|
| Pan-nuclear $\gamma$H2AX | 115 | 28 | 143 |
| Other $\gamma$H2AX | 18 | 34 | 52 |
| Total | 133 | 62 | 195 |

| Expected | Zta+ | Zta- | Total |
|---|---|---|---|
| Pan-nuclear $\gamma$H2AX | 97.5 | 45.5 | 143 |
| Other $\gamma$H2AX | 35.5 | 16.5 | 52 |
| Total | 133 | 62 | 195 |

**Chi-squared test p = 1.25e-9**
