## Supplementary material for "Reconstructing EBV reactivation and DNA damage response kinetics in morphologic pseudotime": Figure S13

**A**

| P3HR1 TGFβ (n=107) | Zta+ | Zta- | Total |
|---|---|---|---|
| 53BP1 Puncta + | 3 | 49 | 52 |
| 53BP1 Puncta - | 21 | 34 | 55 |
| Total | 24 | 83 | 107 |

| Expected | Zta+ | Zta- | Total |
|---|---|---|---|
| 53BP1 Puncta + | 11.7 | 40.3 | 52 |
| 53BP1 Puncta - | 12.3 | 42.7 | 55 |
| Total | 24 | 83 | 107 |

**Chi-squared test p = 5.88e-5**

**B**

| Mutu TGFβ (n=125) | Zta+ | Zta- | Total |
|---|---|---|---|
| 53BP1 Puncta + | 1 | 67 | 68 |
| 53BP1 Puncta - | 24 | 33 | 57 |
| Total | 25 | 100 | 125 |

| Expected | Zta+ | Zta- | Total |
|---|---|---|---|
| 53BP1 Puncta + | 13.6 | 54.4 | 68 |
| 53BP1 Puncta - | 11.4 | 45.6 | 57 |
| Total | 25 | 100 | 125 |

**Chi-squared test p = 1.54e-8**
