## Supplementary material for "Reconstructing EBV reactivation and DNA damage response kinetics in morphologic pseudotime": File S2

CellProfiler Pipeline: http://www.cellprofiler.org
Version:5
DateRevision:428
GitHash:
ModuleCount:29
HasImagePlaneDetails:False

Images:[module_num:1|svn_version:'Unknown'|variable_revision_number:2|show_window:False|notes:['To begin creating your project, use the Images module to compile a list of files and/or folders that you want to analyze. You can also specify a set of rules to include only the desired files in your selected folders.']|batch_state:array([], dtype=uint8)|enabled:True|wants_pause:False]
    :
    Filter images?:Custom
    Select the rule criteria:and (extension does isimage) (file does contain "ES-0001") (file does contain "Z04")

Metadata:[module_num:2|svn_version:'Unknown'|variable_revision_number:6|show_window:False|notes:['The Metadata module optionally allows you to extract information describing your images (i.e, metadata) which will be stored along with your measurements. This information can be contained in the file name and/or location, or in an external file.']|batch_state:array([], dtype=uint8)|enabled:True|wants_pause:False]
    Extract metadata?:Yes
    Metadata data type:Text
    Metadata types:{}
    Extraction method count:2
    Metadata extraction method:Extract from file/folder names
    Metadata source:File name
    Regular expression to extract from file name:^(?P<PlateType>.*)_(?P<WellID>.*)_T(?P<timepoint>[0-9]{4})F(?P<Field>.*)L01A(?P<acqcycle>[0-9]{2})Z(?P<Zslice>[0-9]{2})C(?P<dye>[0-9]{2}).tif
    Regular expression to extract from folder name:(?P<Date>[0-9]{4}_[0-9]{2}_[0-9]{2})$
    Extract metadata from:Images matching a rule
    Select the filtering criteria:and (file does contain "Z04")
    Metadata file location:Elsewhere...|
    Match file and image metadata:[]
    Use case insensitive matching?:No
    Metadata file name:None
    Does cached metadata exist?:No
    Metadata extraction method:Extract from file/folder names
    Metadata source:Folder name
    Regular expression to extract from file name:^(?P<Plate>.*)_(?P<Well>[A-P][0-9]{2})_s(?P<Site>[0-9])_w(?P<ChannelNumber>[0-9])
    Regular expression to extract from folder name:.*[\\/](?P<PlateID>.*)[\\/](?P<Folder>.*)$
    Extract metadata from:All images
    Select the filtering criteria:and (file does contain "")
    Metadata file location:Elsewhere...|
    Match file and image metadata:[]
    Use case insensitive matching?:No
    Metadata file name:None
    Does cached metadata exist?:No

NamesAndTypes:[module_num:3|svn_version:'Unknown'|variable_revision_number:8|show_window:False|notes:['The NamesAndTypes module allows you to assign a meaningful name to each image by which other modules will refer to it.']|batch_state:array([], dtype=uint8)|enabled:True|wants_pause:False]
    Assign a name to:Images matching rules
    Select the image type:Grayscale image
    Name to assign these images:DNA
    Match metadata:[]
    Image set matching method:Order
    Set intensity range from:Image metadata
    Assignments count:5
    Single images count:0
    Maximum intensity:255.0
    Process as 3D?:No
    Relative pixel spacing in X:1.0
    Relative pixel spacing in Y:1.0
    Relative pixel spacing in Z:1.0
    Select the rule criteria:and (file does contain "C01.tif") (metadata does Folder "ES-0001")
    Name to assign these images:DNA
    Name to assign these objects:Cell
    Select the image type:Grayscale image
    Set intensity range from:Image metadata
    Maximum intensity:255.0
    Select the rule criteria:and (file does contain "C02.tif") (metadata does Folder "ES-0001")
    Name to assign these images:gp350
    Name to assign these objects:Cell
    Select the image type:Grayscale image
    Set intensity range from:Image metadata
    Maximum intensity:255.0
    Select the rule criteria:and (file does contain "C03.tif") (metadata does Folder "ES-0001")
    Name to assign these images:EdU
    Name to assign these objects:Cell
    Select the image type:Grayscale image
    Set intensity range from:Image metadata
    Maximum intensity:255.0
    Select the rule criteria:and (file does contain "C04.tif") (metadata does Folder "ES-0001")
    Name to assign these images:yH2AX
    Name to assign these objects:Cell
    Select the image type:Grayscale image
    Set intensity range from:Image metadata
    Maximum intensity:255.0
    Select the rule criteria:and (file does contain "C05.tif") (metadata does Folder "ES-0001")
    Name to assign these images:brightfield
    Name to assign these objects:Cell
    Select the image type:Grayscale image
    Set intensity range from:Image metadata
    Maximum intensity:255.0

Groups:[module_num:4|svn_version:'Unknown'|variable_revision_number:2|show_window:False|notes:['The Groups module optionally allows you to split your list of images into image subsets (groups) which will be processed independently of each other. Examples of groupings include screening batches, microtiter plates, time-lapse movies, etc.']|batch_state:array([], dtype=uint8)|enabled:True|wants_pause:False]
    Do you want to group your images?:No
    grouping metadata count:1
    Metadata category:None

IdentifyPrimaryObjects:[module_num:5|svn_version:'Unknown'|variable_revision_number:15|show_window:False|notes:[]|batch_state:array([], dtype=uint8)|enabled:True|wants_pause:False]
    Select the input image:DNA
    Name the primary objects to be identified:Nucleus
    Typical diameter of objects, in pixel units (Min,Max):50,175
    Discard objects outside the diameter range?:Yes
    Discard objects touching the border of the image?:Yes
    Method to distinguish clumped objects:Shape
    Method to draw dividing lines between clumped objects:Shape
    Size of smoothing filter:50
    Suppress local maxima that are closer than this minimum allowed distance:50
    Speed up by using lower-resolution image to find local maxima?:Yes
    Fill holes in identified objects?:After both thresholding and declumping
    Automatically calculate size of smoothing filter for declumping?:No
    Automatically calculate minimum allowed distance between local maxima?:No
    Handling of objects if excessive number of objects identified:Continue
    Maximum number of objects:500
    Use advanced settings?:Yes
    Threshold setting version:12
    Threshold strategy:Global
    Thresholding method:Otsu
    Threshold smoothing scale:6
    Threshold correction factor:1
    Lower and upper bounds on threshold:0.0,1.0
    Manual threshold:0.0
    Select the measurement to threshold with:None
    Two-class or three-class thresholding?:Two classes
    Log transform before thresholding?:Yes
    Assign pixels in the middle intensity class to the foreground or the background?:Foreground
    Size of adaptive window:50
    Lower outlier fraction:0.05
    Upper outlier fraction:0.05
    Averaging method:Mean
    Variance method:Standard deviation
    # of deviations:2.0
    Thresholding method:Minimum Cross-Entropy

ImageMath:[module_num:6|svn_version:'Unknown'|variable_revision_number:5|show_window:False|notes:[]|batch_state:array([], dtype=uint8)|enabled:True|wants_pause:False]
    Operation:Add
    Raise the power of the result by:1.0
    Multiply the result by:1.0
    Add to result:0.0
    Set values less than 0 equal to 0?:Yes
    Set values greater than 1 equal to 1?:Yes
    Replace invalid values with 0?:Yes
    Ignore the image masks?:No
    Name the output image:Img_Math
    Image or measurement?:Image
    Select the first image:DNA
    Multiply the first image by:3.0
    Measurement:
    Image or measurement?:Image
    Select the second image:gp350
    Multiply the second image by:2.0
    Measurement:
    Image or measurement?:Image
    Select the third image:EdU
    Multiply the third image by:1.0
    Measurement:
    Image or measurement?:Image
    Select the fourth image:yH2AX
    Multiply the fourth image by:1.0
    Measurement:

IdentifySecondaryObjects:[module_num:7|svn_version:'Unknown'|variable_revision_number:10|show_window:False|notes:[]|batch_state:array([], dtype=uint8)|enabled:True|wants_pause:False]
    Select the input objects:Nucleus
    Name the objects to be identified:Cell
    Select the method to identify the secondary objects:Propagation
    Select the input image:Img_Math
    Number of pixels by which to expand the primary objects:10
    Regularization factor:0.05
    Discard secondary objects touching the border of the image?:No
    Discard the associated primary objects?:No
    Name the new primary objects:FilteredNuclei
    Fill holes in identified objects?:Yes
    Threshold setting version:12
    Threshold strategy:Global
    Thresholding method:Minimum Cross-Entropy
    Threshold smoothing scale:0.0
    Threshold correction factor:1.0
    Lower and upper bounds on threshold:0.0,1.0
    Manual threshold:0.0
    Select the measurement to threshold with:None
    Two-class or three-class thresholding?:Two classes
    Log transform before thresholding?:No
    Assign pixels in the middle intensity class to the foreground or the background?:Foreground
    Size of adaptive window:50
    Lower outlier fraction:0.05
    Upper outlier fraction:0.05
    Averaging method:Mean
    Variance method:Standard deviation
    # of deviations:2.0
    Thresholding method:Minimum Cross-Entropy

IdentifyTertiaryObjects:[module_num:8|svn_version:'Unknown'|variable_revision_number:3|show_window:False|notes:[]|batch_state:array([], dtype=uint8)|enabled:True|wants_pause:False]
    Select the larger identified objects:Cell
    Select the smaller identified objects:Nucleus
    Name the tertiary objects to be identified:Cytoplasm
    Shrink smaller object prior to subtraction?:Yes

MeasureObjectIntensity:[module_num:9|svn_version:'Unknown'|variable_revision_number:4|show_window:False|notes:[]|batch_state:array([], dtype=uint8)|enabled:True|wants_pause:False]
    Select images to measure:DNA, EdU, brightfield, gp350, yH2AX
    Select objects to measure:Cell, Cytoplasm, Nucleus

MeasureColocalization:[module_num:10|svn_version:'Unknown'|variable_revision_number:5|show_window:False|notes:[]|batch_state:array([], dtype=uint8)|enabled:True|wants_pause:False]
    Select images to measure:DNA, EdU, gp350, yH2AX
    Set threshold as percentage of maximum intensity for the images:15.0
    Select where to measure correlation:Within objects
    Select objects to measure:Cell, Cytoplasm, Nucleus
    Run all metrics?:Yes
    Calculate correlation and slope metrics?:Yes
    Calculate the Manders coefficients?:Yes
    Calculate the Rank Weighted Colocalization coefficients?:Yes
    Calculate the Overlap coefficients?:Yes
    Calculate the Manders coefficients using Costes auto threshold?:Yes
    Method for Costes thresholding:Faster

MeasureObjectSizeShape:[module_num:11|svn_version:'Unknown'|variable_revision_number:3|show_window:False|notes:[]|batch_state:array([], dtype=uint8)|enabled:True|wants_pause:False]
    Select object sets to measure:Cell, Nucleus
    Calculate the Zernike features?:Yes
    Calculate the advanced features?:Yes

MeasureTexture:[module_num:12|svn_version:'Unknown'|variable_revision_number:7|show_window:False|notes:[]|batch_state:array([], dtype=uint8)|enabled:True|wants_pause:False]
    Select images to measure:DNA, EdU, brightfield, gp350, yH2AX
    Select objects to measure:Cytoplasm, Nucleus
    Enter how many gray levels to measure the texture at:256
    Hidden:2
    Measure whole images or objects?:Objects
    Texture scale to measure:20
    Texture scale to measure:5

ExportToSpreadsheet:[module_num:13|svn_version:'Unknown'|variable_revision_number:13|show_window:False|notes:[]|batch_state:array([], dtype=uint8)|enabled:False|wants_pause:False]
    Select the column delimiter:Comma (",")
    Add image metadata columns to your object data file?:No
    Add image file and folder names to your object data file?:No
    Select the measurements to export:No
    Calculate the per-image mean values for object measurements?:No
    Calculate the per-image median values for object measurements?:No
    Calculate the per-image standard deviation values for object measurements?:No
    Output file location:Elsewhere...|/Users/sorelle/Desktop/microscopy/Yokogawa_CV8000/LyticDDRTimeCourse2Plate_20250711_154416/20250715_CellProfiler_Output/All_Fields/Spreadsheets
    Create a GenePattern GCT file?:No
    Select source of sample row name:Metadata
    Select the image to use as the identifier:None
    Select the metadata to use as the identifier:None
    Export all measurement types?:Yes
    Press button to select measurements:
    Representation of Nan/Inf:NaN
    Add a prefix to file names?:Yes
    Filename prefix:Lytic_DDR_Plate_A_Z03_All_Fields_
    Overwrite existing files without warning?:No
    Data to export:Do not use
    Combine these object measurements with those of the previous object?:No
    File name:DATA.csv
    Use the object name for the file name?:Yes

ExportToDatabase:[module_num:14|svn_version:'Unknown'|variable_revision_number:28|show_window:False|notes:[]|batch_state:array([], dtype=uint8)|enabled:True|wants_pause:False]
    Database type:SQLite
    Database name:DefaultDB
    Add a prefix to table names?:No
    Table prefix:Lytic_DDR_Plate_A_Z03_All_Fields_
    Output file location:Elsewhere...|/Users/sorelle/Desktop/microscopy/Yokogawa_CV8000/raw_data/Plate_A_DNA_gp350_EdU_yH2AX/CP_Database_Plate_A
    Create a CellProfiler Analyst properties file?:Yes
    Database host:
    Username:
    Password:
    Name the SQLite database file:DefaultDB.db
    Calculate the per-image mean values of object measurements?:Yes
    Calculate the per-image median values of object measurements?:No
    Calculate the per-image standard deviation values of object measurements?:No
    Calculate the per-well mean values of object measurements?:No
    Calculate the per-well median values of object measurements?:No
    Calculate the per-well standard deviation values of object measurements?:No
    Export measurements for all objects to the database?:All
    Select the objects:
    Maximum # of characters in a column name:64
    Create one table per object, a single object table or a single object view?:Single object table
    Enter an image url prepend if you plan to access your files via http:
    Write image thumbnails directly to the database?:No
    Select the images for which you want to save thumbnails:
    Auto-scale thumbnail pixel intensities?:Yes
    Select the plate type:96
    Select the plate metadata:PlateID
    Select the well metadata:WellID
    Include information for all images, using default values?:Yes
    Properties image group count:1
    Properties group field count:1
    Properties filter field count:0
    Workspace measurement count:1
    Experiment name:Lytic_DDR_Plate_A_Z04
    Which objects should be used for locations?:Nucleus
    Enter a phenotype class table name if using the Classifier tool in CellProfiler Analyst:
    Export object relationships?:Yes
    Overwrite without warning?:Never
    Access CellProfiler Analyst images via URL?:No
    Select the classification type:Object
    Select an image to include:None
    Use the image name for the display?:Yes
    Image name:Channel1
    Channel color:red
    Do you want to add group fields?:No
    Enter the name of the group:
    Enter the per-image columns which define the group, separated by commas:ImageNumber, Image_Metadata_Plate, Image_Metadata_Well
    Do you want to add filter fields?:No
    Automatically create a filter for each plate?:No
    Create a CellProfiler Analyst workspace file?:No
    Select the measurement display tool:ScatterPlot
    Type of measurement to plot on the X-axis:Image
    Enter the object name:None
    Select the X-axis measurement:None
    Select the X-axis index:ImageNumber
    Type of measurement to plot on the Y-axis:Image
    Enter the object name:None
    Select the Y-axis measurement:None
    Select the Y-axis index:ImageNumber

RescaleIntensity:[module_num:15|svn_version:'Unknown'|variable_revision_number:3|show_window:False|notes:[]|batch_state:array([], dtype=uint8)|enabled:False|wants_pause:False]
    Select the input image:DNA
    Name the output image:rDNA
    Rescaling method:Choose specific values to be reset to the full intensity range
    Method to calculate the minimum intensity:Custom
    Method to calculate the maximum intensity:Custom
    Lower intensity limit for the input image:0.0
    Upper intensity limit for the input image:1.0
    Intensity range for the input image:0.001,.04
    Intensity range for the output image:0.0,1.0
    Select image to match in maximum intensity:None
    Divisor value:1.0
    Divisor measurement:None

RescaleIntensity:[module_num:16|svn_version:'Unknown'|variable_revision_number:3|show_window:False|notes:[]|batch_state:array([], dtype=uint8)|enabled:False|wants_pause:False]
    Select the input image:EdU
    Name the output image:rEdU
    Rescaling method:Choose specific values to be reset to the full intensity range
    Method to calculate the minimum intensity:Custom
    Method to calculate the maximum intensity:Custom
    Lower intensity limit for the input image:0.0
    Upper intensity limit for the input image:1.0
    Intensity range for the input image:0.001,.02
    Intensity range for the output image:0.0,1.0
    Select image to match in maximum intensity:None
    Divisor value:1.0
    Divisor measurement:None

RescaleIntensity:[module_num:17|svn_version:'Unknown'|variable_revision_number:3|show_window:False|notes:[]|batch_state:array([], dtype=uint8)|enabled:False|wants_pause:False]
    Select the input image:gp350
    Name the output image:rgp350
    Rescaling method:Choose specific values to be reset to the full intensity range
    Method to calculate the minimum intensity:Custom
    Method to calculate the maximum intensity:Custom
    Lower intensity limit for the input image:0.0
    Upper intensity limit for the input image:1.0
    Intensity range for the input image:0.005,.02
    Intensity range for the output image:0.0,1.0
    Select image to match in maximum intensity:None
    Divisor value:1.0
    Divisor measurement:None

RescaleIntensity:[module_num:18|svn_version:'Unknown'|variable_revision_number:3|show_window:False|notes:[]|batch_state:array([], dtype=uint8)|enabled:False|wants_pause:False]
    Select the input image:brightfield
    Name the output image:rBF
    Rescaling method:Choose specific values to be reset to the full intensity range
    Method to calculate the minimum intensity:Minimum for each image
    Method to calculate the maximum intensity:Maximum for each image
    Lower intensity limit for the input image:0.0
    Upper intensity limit for the input image:0.005
    Intensity range for the input image:0.006,.01
    Intensity range for the output image:0.0,1.0
    Select image to match in maximum intensity:None
    Divisor value:1.0
    Divisor measurement:None

RescaleIntensity:[module_num:19|svn_version:'Unknown'|variable_revision_number:3|show_window:False|notes:[]|batch_state:array([], dtype=uint8)|enabled:False|wants_pause:False]
    Select the input image:yH2AX
    Name the output image:ryH2AX
    Rescaling method:Choose specific values to be reset to the full intensity range
    Method to calculate the minimum intensity:Custom
    Method to calculate the maximum intensity:Custom
    Lower intensity limit for the input image:0.0
    Upper intensity limit for the input image:1.0
    Intensity range for the input image:0.001,.04
    Intensity range for the output image:0.0,1.0
    Select image to match in maximum intensity:None
    Divisor value:1.0
    Divisor measurement:None

GrayToColor:[module_num:20|svn_version:'Unknown'|variable_revision_number:4|show_window:False|notes:[]|batch_state:array([], dtype=uint8)|enabled:False|wants_pause:False]
    Select a color scheme:Composite
    Rescale intensity:Yes
    Select the image to be colored red:rDNA
    Select the image to be colored green:rgp350
    Select the image to be colored blue:ryH2AX
    Name the output image:Composite
    Relative weight for the red image:1.0
    Relative weight for the green image:1.0
    Relative weight for the blue image:1.0
    Select the image to be colored cyan:Leave this black
    Select the image to be colored magenta:Leave this black
    Select the image to be colored yellow:Leave this black
    Select the image that determines brightness:Leave this black
    Relative weight for the cyan image:1.0
    Relative weight for the magenta image:1.0
    Relative weight for the yellow image:1.0
    Relative weight for the brightness image:1.0
    Hidden:4
    Image name:rDNA
    Color:#0011F4
    Weight:1.0
    Image name:rgp350
    Color:#00FCF6
    Weight:1.0
    Image name:rEdU
    Color:#FCC300
    Weight:1.0
    Image name:ryH2AX
    Color:#FF0000
    Weight:1.0

GrayToColor:[module_num:21|svn_version:'Unknown'|variable_revision_number:4|show_window:False|notes:[]|batch_state:array([], dtype=uint8)|enabled:False|wants_pause:False]
    Select a color scheme:Composite
    Rescale intensity:Yes
    Select the image to be colored red:rDNA
    Select the image to be colored green:rgp350
    Select the image to be colored blue:ryH2AX
    Name the output image:Composite_BF
    Relative weight for the red image:1.0
    Relative weight for the green image:1.0
    Relative weight for the blue image:1.0
    Select the image to be colored cyan:Leave this black
    Select the image to be colored magenta:Leave this black
    Select the image to be colored yellow:Leave this black
    Select the image that determines brightness:Leave this black
    Relative weight for the cyan image:1.0
    Relative weight for the magenta image:1.0
    Relative weight for the yellow image:1.0
    Relative weight for the brightness image:1.0
    Hidden:5
    Image name:rDNA
    Color:#0011F4
    Weight:1.0
    Image name:rgp350
    Color:#00F0F6
    Weight:1.0
    Image name:rEdU
    Color:#FCC300
    Weight:1.0
    Image name:ryH2AX
    Color:#FF0000
    Weight:1.0
    Image name:rBF
    Color:#9E9493
    Weight:1

GrayToColor:[module_num:22|svn_version:'Unknown'|variable_revision_number:4|show_window:False|notes:[]|batch_state:array([], dtype=uint8)|enabled:False|wants_pause:False]
    Select a color scheme:Composite
    Rescale intensity:Yes
    Select the image to be colored red:Leave this black
    Select the image to be colored green:Leave this black
    Select the image to be colored blue:Leave this black
    Name the output image:DNA_Color
    Relative weight for the red image:1.0
    Relative weight for the green image:1.0
    Relative weight for the blue image:1.0
    Select the image to be colored cyan:Leave this black
    Select the image to be colored magenta:Leave this black
    Select the image to be colored yellow:Leave this black
    Select the image that determines brightness:Leave this black
    Relative weight for the cyan image:1.0
    Relative weight for the magenta image:1.0
    Relative weight for the yellow image:1.0
    Relative weight for the brightness image:1.0
    Hidden:1
    Image name:rDNA
    Color:#001FF4
    Weight:1.0

GrayToColor:[module_num:23|svn_version:'Unknown'|variable_revision_number:4|show_window:False|notes:[]|batch_state:array([], dtype=uint8)|enabled:False|wants_pause:False]
    Select a color scheme:Composite
    Rescale intensity:Yes
    Select the image to be colored red:Leave this black
    Select the image to be colored green:Leave this black
    Select the image to be colored blue:Leave this black
    Name the output image:gp350_Color
    Relative weight for the red image:1.0
    Relative weight for the green image:1.0
    Relative weight for the blue image:1.0
    Select the image to be colored cyan:Leave this black
    Select the image to be colored magenta:Leave this black
    Select the image to be colored yellow:Leave this black
    Select the image that determines brightness:Leave this black
    Relative weight for the cyan image:1.0
    Relative weight for the magenta image:1.0
    Relative weight for the yellow image:1.0
    Relative weight for the brightness image:1.0
    Hidden:1
    Image name:rgp350
    Color:#00E9EB
    Weight:1.0

GrayToColor:[module_num:24|svn_version:'Unknown'|variable_revision_number:4|show_window:False|notes:[]|batch_state:array([], dtype=uint8)|enabled:False|wants_pause:False]
    Select a color scheme:Composite
    Rescale intensity:Yes
    Select the image to be colored red:Leave this black
    Select the image to be colored green:Leave this black
    Select the image to be colored blue:Leave this black
    Name the output image:EdU_Color
    Relative weight for the red image:1.0
    Relative weight for the green image:1.0
    Relative weight for the blue image:1.0
    Select the image to be colored cyan:Leave this black
    Select the image to be colored magenta:Leave this black
    Select the image to be colored yellow:Leave this black
    Select the image that determines brightness:Leave this black
    Relative weight for the cyan image:1.0
    Relative weight for the magenta image:1.0
    Relative weight for the yellow image:1.0
    Relative weight for the brightness image:1.0
    Hidden:1
    Image name:rEdU
    Color:#F0C100
    Weight:1.0

GrayToColor:[module_num:25|svn_version:'Unknown'|variable_revision_number:4|show_window:False|notes:[]|batch_state:array([], dtype=uint8)|enabled:False|wants_pause:False]
    Select a color scheme:Composite
    Rescale intensity:Yes
    Select the image to be colored red:Leave this black
    Select the image to be colored green:Leave this black
    Select the image to be colored blue:Leave this black
    Name the output image:yH2AX_Color
    Relative weight for the red image:1.0
    Relative weight for the green image:1.0
    Relative weight for the blue image:1.0
    Select the image to be colored cyan:Leave this black
    Select the image to be colored magenta:Leave this black
    Select the image to be colored yellow:Leave this black
    Select the image that determines brightness:Leave this black
    Relative weight for the cyan image:1.0
    Relative weight for the magenta image:1.0
    Relative weight for the yellow image:1.0
    Relative weight for the brightness image:1.0
    Hidden:1
    Image name:ryH2AX
    Color:#FF0000
    Weight:1.0

OverlayOutlines:[module_num:26|svn_version:'Unknown'|variable_revision_number:4|show_window:False|notes:[]|batch_state:array([], dtype=uint8)|enabled:False|wants_pause:False]
    Display outlines on a blank image?:No
    Select image on which to display outlines:rDNA
    Name the output image:OrigOverlay
    Outline display mode:Color
    Select method to determine brightness of outlines:Max of image
    How to outline:Thick
    Select outline color:#00F600
    Select objects to display:Nucleus

Tile:[module_num:27|svn_version:'Unknown'|variable_revision_number:1|show_window:False|notes:[]|batch_state:array([], dtype=uint8)|enabled:False|wants_pause:False]
    Select an input image:DNA_Color
    Name the output image:TiledImage
    Tile assembly method:Within cycles
    Final number of rows:2
    Final number of columns:4
    Image corner to begin tiling:top left
    Direction to begin tiling:row
    Use meander mode?:No
    Automatically calculate number of rows?:No
    Automatically calculate number of columns?:No
    Select an additional image to tile:gp350_Color
    Select an additional image to tile:Composite
    Select an additional image to tile:rBF
    Select an additional image to tile:EdU_Color
    Select an additional image to tile:yH2AX_Color
    Select an additional image to tile:OrigOverlay
    Select an additional image to tile:Composite_BF

RescaleIntensity:[module_num:28|svn_version:'Unknown'|variable_revision_number:3|show_window:False|notes:[]|batch_state:array([], dtype=uint8)|enabled:False|wants_pause:True]
    Select the input image:TiledImage
    Name the output image:rTiled
    Rescaling method:Stretch each image to use the full intensity range
    Method to calculate the minimum intensity:Custom
    Method to calculate the maximum intensity:Custom
    Lower intensity limit for the input image:0.0
    Upper intensity limit for the input image:1.0
    Intensity range for the input image:0.001,.004
    Intensity range for the output image:0.0,1.0
    Select image to match in maximum intensity:None
    Divisor value:1.0
    Divisor measurement:None

SaveImages:[module_num:29|svn_version:'Unknown'|variable_revision_number:16|show_window:False|notes:[]|batch_state:array([], dtype=uint8)|enabled:False|wants_pause:False]
    Select the type of image to save:Image
    Select the image to save:rTiled
    Select method for constructing file names:Single name
    Select image name for file prefix:None
    Enter single file name:\g<WellID>_\g<Field>
    Number of digits:4
    Append a suffix to the image file name?:No
    Text to append to the image name:
    Saved file format:jpeg
    Output file location:Elsewhere...|/Users/sorelle/Desktop/microscopy/Yokogawa_CV8000/raw_data/Plate_A_DNA_gp350_EdU_yH2AX/CP_Tiles_Plate_A
    Image bit depth:8-bit integer
    Overwrite existing files without warning?:Yes
    When to save:Every cycle
    Record the file and path information to the saved image?:No
    Create subfolders in the output folder?:No
    Base image folder:Elsewhere...|
    How to save the series:T (Time)
    Save with lossless compression?:Yes
