## Supplementary material for "Reconstructing EBV reactivation and DNA damage response kinetics in morphologic pseudotime": File S3

CellProfiler Pipeline: http://www.cellprofiler.org
Version:5
DateRevision:428
GitHash:
ModuleCount:36
HasImagePlaneDetails:False

Images:[module_num:1|svn_version:'Unknown'|variable_revision_number:2|show_window:False|notes:['To begin creating your project, use the Images module to compile a list of files and/or folders that you want to analyze. You can also specify a set of rules to include only the desired files in your selected folders.']|batch_state:array([], dtype=uint8)|enabled:True|wants_pause:False]
    :
    Filter images?:Custom
    Select the rule criteria:and (extension does istif) (file does contain "treat_A")

Metadata:[module_num:2|svn_version:'Unknown'|variable_revision_number:6|show_window:False|notes:['The Metadata module optionally allows you to extract information describing your images (i.e, metadata) which will be stored along with your measurements. This information can be contained in the file name and/or location, or in an external file.']|batch_state:array([], dtype=uint8)|enabled:True|wants_pause:False]
    Extract metadata?:Yes
    Metadata data type:Text
    Metadata types:{}
    Extraction method count:1
    Metadata extraction method:Extract from file/folder names
    Metadata source:File name
    Regular expression to extract from file name:^(........)_(..................................)_(?P<Row>[A-H])_(?P<Col>[0-9])_(...)_s(?P<Stage>[0-9][0-9])_ch(?P<Channel>[0-9][0-9])
    Regular expression to extract from folder name:(?P<Date>[0-9]{4}_[0-9]{2}_[0-9]{2})$
    Extract metadata from:All images
    Select the filtering criteria:and (file does contain "")
    Metadata file location:Elsewhere...|
    Match file and image metadata:[]
    Use case insensitive matching?:No
    Metadata file name:None
    Does cached metadata exist?:No

NamesAndTypes:[module_num:3|svn_version:'Unknown'|variable_revision_number:8|show_window:False|notes:['The NamesAndTypes module allows you to assign a meaningful name to each image by which other modules will refer to it.']|batch_state:array([], dtype=uint8)|enabled:True|wants_pause:False]
    Assign a name to:Images matching rules
    Select the image type:Grayscale image
    Name to assign these images:DNA
    Match metadata:[]
    Image set matching method:Order
    Set intensity range from:Image metadata
    Assignments count:6
    Single images count:0
    Maximum intensity:255.0
    Process as 3D?:No
    Relative pixel spacing in X:1.0
    Relative pixel spacing in Y:1.0
    Relative pixel spacing in Z:1.0
    Select the rule criteria:and (file does contain "ch00")
    Name to assign these images:DNA
    Name to assign these objects:Cell
    Select the image type:Grayscale image
    Set intensity range from:Image metadata
    Maximum intensity:255.0
    Select the rule criteria:and (file does contain "ch01")
    Name to assign these images:Zta
    Name to assign these objects:Nucleus
    Select the image type:Grayscale image
    Set intensity range from:Image metadata
    Maximum intensity:255.0
    Select the rule criteria:and (file does contain "ch02")
    Name to assign these images:MTO
    Name to assign these objects:Cytoplasm
    Select the image type:Grayscale image
    Set intensity range from:Image metadata
    Maximum intensity:255.0
    Select the rule criteria:and (file does contain "ch03")
    Name to assign these images:Ki67
    Name to assign these objects:Speckle
    Select the image type:Grayscale image
    Set intensity range from:Image metadata
    Maximum intensity:255.0
    Select the rule criteria:and (file does contain "ch04")
    Name to assign these images:Dead
    Name to assign these objects:Object1
    Select the image type:Grayscale image
    Set intensity range from:Image metadata
    Maximum intensity:255.0
    Select the rule criteria:and (file does contain "ch05")
    Name to assign these images:Bright
    Name to assign these objects:Object2
    Select the image type:Grayscale image
    Set intensity range from:Image metadata
    Maximum intensity:255.0

Groups:[module_num:4|svn_version:'Unknown'|variable_revision_number:2|show_window:False|notes:['The Groups module optionally allows you to split your list of images into image subsets (groups) which will be processed independently of each other. Examples of groupings include screening batches, microtiter plates, time-lapse movies, etc.']|batch_state:array([], dtype=uint8)|enabled:True|wants_pause:False]
    Do you want to group your images?:No
    grouping metadata count:1
    Metadata category:None

RescaleIntensity:[module_num:5|svn_version:'Unknown'|variable_revision_number:3|show_window:False|notes:[]|batch_state:array([], dtype=uint8)|enabled:True|wants_pause:False]
    Select the input image:DNA
    Name the output image:DNA_rescale
    Rescaling method:Choose specific values to be reset to a custom range
    Method to calculate the minimum intensity:Minimum for each image
    Method to calculate the maximum intensity:Custom
    Lower intensity limit for the input image:0.0
    Upper intensity limit for the input image:1
    Intensity range for the input image:0.0,1.0
    Intensity range for the output image:0.0,20
    Select image to match in maximum intensity:None
    Divisor value:1.0
    Divisor measurement:None

RescaleIntensity:[module_num:6|svn_version:'Unknown'|variable_revision_number:3|show_window:False|notes:[]|batch_state:array([], dtype=uint8)|enabled:True|wants_pause:False]
    Select the input image:Zta
    Name the output image:Zta_rescale
    Rescaling method:Choose specific values to be reset to a custom range
    Method to calculate the minimum intensity:Minimum for each image
    Method to calculate the maximum intensity:Custom
    Lower intensity limit for the input image:0.0
    Upper intensity limit for the input image:1
    Intensity range for the input image:0.0,1.0
    Intensity range for the output image:0.0,20
    Select image to match in maximum intensity:None
    Divisor value:1.0
    Divisor measurement:None

RescaleIntensity:[module_num:7|svn_version:'Unknown'|variable_revision_number:3|show_window:False|notes:[]|batch_state:array([], dtype=uint8)|enabled:True|wants_pause:False]
    Select the input image:MTO
    Name the output image:MTO_rescale
    Rescaling method:Choose specific values to be reset to a custom range
    Method to calculate the minimum intensity:Minimum for each image
    Method to calculate the maximum intensity:Custom
    Lower intensity limit for the input image:0.0
    Upper intensity limit for the input image:1
    Intensity range for the input image:0.0,1.0
    Intensity range for the output image:0.0,20
    Select image to match in maximum intensity:None
    Divisor value:1.0
    Divisor measurement:None

RescaleIntensity:[module_num:8|svn_version:'Unknown'|variable_revision_number:3|show_window:False|notes:[]|batch_state:array([], dtype=uint8)|enabled:True|wants_pause:False]
    Select the input image:Ki67
    Name the output image:Ki67_rescale
    Rescaling method:Choose specific values to be reset to a custom range
    Method to calculate the minimum intensity:Minimum for each image
    Method to calculate the maximum intensity:Custom
    Lower intensity limit for the input image:0.0
    Upper intensity limit for the input image:1
    Intensity range for the input image:0.0,1.0
    Intensity range for the output image:0.0,20
    Select image to match in maximum intensity:None
    Divisor value:1.0
    Divisor measurement:None

RescaleIntensity:[module_num:9|svn_version:'Unknown'|variable_revision_number:3|show_window:False|notes:[]|batch_state:array([], dtype=uint8)|enabled:True|wants_pause:False]
    Select the input image:Dead
    Name the output image:Dead_rescale
    Rescaling method:Choose specific values to be reset to a custom range
    Method to calculate the minimum intensity:Minimum for each image
    Method to calculate the maximum intensity:Custom
    Lower intensity limit for the input image:0.0
    Upper intensity limit for the input image:1
    Intensity range for the input image:0.0,1.0
    Intensity range for the output image:0.0,20
    Select image to match in maximum intensity:None
    Divisor value:1.0
    Divisor measurement:None

RescaleIntensity:[module_num:10|svn_version:'Unknown'|variable_revision_number:3|show_window:False|notes:[]|batch_state:array([], dtype=uint8)|enabled:True|wants_pause:False]
    Select the input image:Bright
    Name the output image:Bright_rescale
    Rescaling method:Choose specific values to be reset to a custom range
    Method to calculate the minimum intensity:Minimum for each image
    Method to calculate the maximum intensity:Custom
    Lower intensity limit for the input image:0.0
    Upper intensity limit for the input image:1
    Intensity range for the input image:0.0,1.0
    Intensity range for the output image:0.0,10
    Select image to match in maximum intensity:None
    Divisor value:1.0
    Divisor measurement:None

ImageMath:[module_num:11|svn_version:'Unknown'|variable_revision_number:5|show_window:False|notes:[]|batch_state:array([], dtype=uint8)|enabled:True|wants_pause:False]
    Operation:Add
    Raise the power of the result by:1.0
    Multiply the result by:1.0
    Add to result:0.0
    Set values less than 0 equal to 0?:Yes
    Set values greater than 1 equal to 1?:Yes
    Replace invalid values with 0?:Yes
    Ignore the image masks?:No
    Name the output image:Composite_rescale
    Image or measurement?:Image
    Select the first image:DNA_rescale
    Multiply the first image by:1.3
    Measurement:
    Image or measurement?:Image
    Select the second image:Zta_rescale
    Multiply the second image by:1.0
    Measurement:
    Image or measurement?:Image
    Select the third image:MTO_rescale
    Multiply the third image by:1.0
    Measurement:
    Image or measurement?:Image
    Select the fourth image:Ki67_rescale
    Multiply the fourth image by:1.0
    Measurement:
    Image or measurement?:Image
    Select the fifth image:Dead_rescale
    Multiply the fifth image by:1.0
    Measurement:

ReduceNoise:[module_num:12|svn_version:'Unknown'|variable_revision_number:1|show_window:False|notes:[]|batch_state:array([], dtype=uint8)|enabled:True|wants_pause:False]
    Select the input image:Composite_rescale
    Name the output image:Comp_denoised
    Size:7
    Distance:11
    Cut-off distance:0.1

ErodeImage:[module_num:13|svn_version:'Unknown'|variable_revision_number:1|show_window:False|notes:[]|batch_state:array([], dtype=uint8)|enabled:True|wants_pause:False]
    Select the input image:Comp_denoised
    Name the output image:Comp_erode
    Structuring element:disk,5

RemoveHoles:[module_num:14|svn_version:'Unknown'|variable_revision_number:1|show_window:False|notes:[]|batch_state:array([], dtype=uint8)|enabled:True|wants_pause:False]
    Select the input image:Comp_erode
    Name the output image:Comp_filled
    Size of holes to fill:50

Watershed:[module_num:15|svn_version:'Unknown'|variable_revision_number:3|show_window:False|notes:[]|batch_state:array([], dtype=uint8)|enabled:True|wants_pause:False]
    Select the input image:Comp_filled
    Name the output object:Watershed
    Use advanced settings?:Yes
    Generate from:Distance
    Markers:None
    Mask:Leave blank
    Connectivity:1
    Compactness:0.0
    Footprint:12
    Downsample:1
    Separate watershed labels:No
    Declump method:Shape
    Reference Image:None
    Segmentation distance transform smoothing factor:1.0
    Minimum distance between seeds:30
    Minimum absolute internal distance:0.0
    Pixels from border to exclude:0
    Maximum number of seeds:-1
    Structuring element for seed dilation:disk,3

FillObjects:[module_num:16|svn_version:'Unknown'|variable_revision_number:2|show_window:False|notes:[]|batch_state:array([], dtype=uint8)|enabled:True|wants_pause:False]
    Select the input object:Watershed
    Name the output object:Watershed_Cells
    Minimum hole size:64.0
    Planewise fill:No
    Filling method:Holes

FilterObjects:[module_num:17|svn_version:'Unknown'|variable_revision_number:10|show_window:False|notes:[]|batch_state:array([], dtype=uint8)|enabled:True|wants_pause:False]
    Select the objects to filter:Watershed_Cells
    Name the output objects:Filtered_Cells
    Select the filtering mode:Image or mask border
    Select the filtering method:Limits
    Select the objects that contain the filtered objects:None
    Select the location of the rules or classifier file:Elsewhere...|
    Rules or classifier file name:rules.txt
    Class number:1
    Measurement count:1
    Additional object count:0
    Assign overlapping child to:Both parents
    Keep removed objects as a separate set?:No
    Name the objects removed by the filter:RemovedObjects
    Select the measurement to filter by:AreaShape_Area
    Filter using a minimum measurement value?:Yes
    Minimum value:0.0
    Filter using a maximum measurement value?:Yes
    Maximum value:1.0
    Allow fuzzy feature matching?:No

MeasureObjectSizeShape:[module_num:18|svn_version:'Unknown'|variable_revision_number:3|show_window:False|notes:[]|batch_state:array([], dtype=uint8)|enabled:True|wants_pause:False]
    Select object sets to measure:Filtered_Cells
    Calculate the Zernike features?:Yes
    Calculate the advanced features?:Yes

MeasureObjectIntensity:[module_num:19|svn_version:'Unknown'|variable_revision_number:4|show_window:False|notes:[]|batch_state:array([], dtype=uint8)|enabled:True|wants_pause:False]
    Select images to measure:Bright, DNA, Dead, Ki67, MTO, Zta
    Select objects to measure:Filtered_Cells

MeasureObjectIntensityDistribution:[module_num:20|svn_version:'Unknown'|variable_revision_number:6|show_window:False|notes:[]|batch_state:array([], dtype=uint8)|enabled:True|wants_pause:False]
    Select images to measure:Bright, DNA, Ki67, MTO, Zta
    Hidden:1
    Hidden:1
    Hidden:0
    Calculate intensity Zernikes?:Magnitudes and phase
    Maximum zernike moment:9
    Select objects to measure:Filtered_Cells
    Object to use as center?:These objects
    Select objects to use as centers:None
    Scale the bins?:Yes
    Number of bins:4
    Maximum radius:100

MeasureTexture:[module_num:21|svn_version:'Unknown'|variable_revision_number:7|show_window:False|notes:[]|batch_state:array([], dtype=uint8)|enabled:True|wants_pause:False]
    Select images to measure:Bright, DNA, Dead, Ki67, MTO, Zta
    Select objects to measure:Filtered_Cells
    Enter how many gray levels to measure the texture at:256
    Hidden:1
    Measure whole images or objects?:Objects
    Texture scale to measure:3

MeasureGranularity:[module_num:22|svn_version:'Unknown'|variable_revision_number:4|show_window:False|notes:[]|batch_state:array([], dtype=uint8)|enabled:False|wants_pause:False]
    Select images to measure:Bright, DNA, Dead, Ki67, MTO, Zta
    Measure within objects?:Yes
    Select objects to measure:Filtered_Cells
    Subsampling factor for granularity measurements:0.25
    Subsampling factor for background reduction:0.25
    Radius of structuring element:10
    Range of the granular spectrum:16

FilterObjects:[module_num:23|svn_version:'Unknown'|variable_revision_number:10|show_window:False|notes:[]|batch_state:array([], dtype=uint8)|enabled:True|wants_pause:False]
    Select the objects to filter:Filtered_Cells
    Name the output objects:QC_Cells
    Select the filtering mode:Measurements
    Select the filtering method:Limits
    Select the objects that contain the filtered objects:None
    Select the location of the rules or classifier file:Elsewhere...|
    Rules or classifier file name:rules.txt
    Class number:1
    Measurement count:2
    Additional object count:0
    Assign overlapping child to:Both parents
    Keep removed objects as a separate set?:No
    Name the objects removed by the filter:RemovedObjects
    Select the measurement to filter by:AreaShape_Area
    Filter using a minimum measurement value?:Yes
    Minimum value:2200
    Filter using a maximum measurement value?:Yes
    Maximum value:28000
    Select the measurement to filter by:AreaShape_Extent
    Filter using a minimum measurement value?:Yes
    Minimum value:0.5
    Filter using a maximum measurement value?:Yes
    Maximum value:1.0
    Allow fuzzy feature matching?:No

ImageMath:[module_num:24|svn_version:'Unknown'|variable_revision_number:5|show_window:False|notes:[]|batch_state:array([], dtype=uint8)|enabled:False|wants_pause:False]
    Operation:Add
    Raise the power of the result by:1.0
    Multiply the result by:1.0
    Add to result:0.0
    Set values less than 0 equal to 0?:Yes
    Set values greater than 1 equal to 1?:Yes
    Replace invalid values with 0?:Yes
    Ignore the image masks?:No
    Name the output image:Aesthetic
    Image or measurement?:Image
    Select the first image:Bright_rescale
    Multiply the first image by:1.0
    Measurement:
    Image or measurement?:Image
    Select the second image:DNA_rescale
    Multiply the second image by:1.0
    Measurement:
    Image or measurement?:Image
    Select the third image:Zta_rescale
    Multiply the third image by:1.0
    Measurement:
    Image or measurement?:Image
    Select the fourth image:MTO_rescale
    Multiply the fourth image by:1.0
    Measurement:
    Image or measurement?:Image
    Select the fifth image:Ki67_rescale
    Multiply the fifth image by:1.0
    Measurement:
    Image or measurement?:Image
    Select the sixth image:Dead_rescale
    Multiply the sixth image by:1.0
    Measurement:

MaskImage:[module_num:25|svn_version:'Unknown'|variable_revision_number:3|show_window:False|notes:[]|batch_state:array([], dtype=uint8)|enabled:False|wants_pause:False]
    Select the input image:DNA_rescale
    Name the output image:Masked_DNA
    Use objects or an image as a mask?:Objects
    Select object for mask:QC_Cells
    Select image for mask:None
    Invert the mask?:No

MaskImage:[module_num:26|svn_version:'Unknown'|variable_revision_number:3|show_window:False|notes:[]|batch_state:array([], dtype=uint8)|enabled:False|wants_pause:False]
    Select the input image:Zta_rescale
    Name the output image:Masked_Zta
    Use objects or an image as a mask?:Objects
    Select object for mask:QC_Cells
    Select image for mask:None
    Invert the mask?:No

MaskImage:[module_num:27|svn_version:'Unknown'|variable_revision_number:3|show_window:False|notes:[]|batch_state:array([], dtype=uint8)|enabled:False|wants_pause:False]
    Select the input image:MTO_rescale
    Name the output image:Masked_MTO
    Use objects or an image as a mask?:Objects
    Select object for mask:QC_Cells
    Select image for mask:None
    Invert the mask?:No

MaskImage:[module_num:28|svn_version:'Unknown'|variable_revision_number:3|show_window:False|notes:[]|batch_state:array([], dtype=uint8)|enabled:False|wants_pause:False]
    Select the input image:Ki67_rescale
    Name the output image:Masked_Ki67
    Use objects or an image as a mask?:Objects
    Select object for mask:QC_Cells
    Select image for mask:None
    Invert the mask?:No

MaskImage:[module_num:29|svn_version:'Unknown'|variable_revision_number:3|show_window:False|notes:[]|batch_state:array([], dtype=uint8)|enabled:False|wants_pause:False]
    Select the input image:Dead_rescale
    Name the output image:Masked_Dead
    Use objects or an image as a mask?:Objects
    Select object for mask:QC_Cells
    Select image for mask:None
    Invert the mask?:No

MaskImage:[module_num:30|svn_version:'Unknown'|variable_revision_number:3|show_window:False|notes:[]|batch_state:array([], dtype=uint8)|enabled:False|wants_pause:False]
    Select the input image:Aesthetic
    Name the output image:Aes_Mask
    Use objects or an image as a mask?:Objects
    Select object for mask:QC_Cells
    Select image for mask:None
    Invert the mask?:No

GrayToColor:[module_num:31|svn_version:'Unknown'|variable_revision_number:4|show_window:False|notes:[]|batch_state:array([], dtype=uint8)|enabled:False|wants_pause:False]
    Select a color scheme:Composite
    Rescale intensity:Yes
    Select the image to be colored red:MTO_quantify
    Select the image to be colored green:GFP_quantify
    Select the image to be colored blue:Hoechst_quantify
    Name the output image:Merge
    Relative weight for the red image:3.0
    Relative weight for the green image:1.0
    Relative weight for the blue image:4.0
    Select the image to be colored cyan:Leave this black
    Select the image to be colored magenta:Leave this black
    Select the image to be colored yellow:Leave this black
    Select the image that determines brightness:Leave this black
    Relative weight for the cyan image:1.0
    Relative weight for the magenta image:1.0
    Relative weight for the yellow image:1.0
    Relative weight for the brightness image:1.0
    Hidden:6
    Image name:Masked_DNA
    Color:#002AF4
    Weight:10
    Image name:Masked_Zta
    Color:#00FF55
    Weight:1
    Image name:Masked_MTO
    Color:#E900F3
    Weight:3
    Image name:Masked_Ki67
    Color:#C40000
    Weight:5
    Image name:Masked_Dead
    Color:#FFFE00
    Weight:1
    Image name:Bright_rescale
    Color:#959C9E
    Weight:3

OverlayOutlines:[module_num:32|svn_version:'Unknown'|variable_revision_number:4|show_window:False|notes:[]|batch_state:array([], dtype=uint8)|enabled:False|wants_pause:False]
    Display outlines on a blank image?:No
    Select image on which to display outlines:Merge
    Name the output image:SanityCheck
    Outline display mode:Color
    Select method to determine brightness of outlines:Max of image
    How to outline:Inner
    Select outline color:#FF00BE
    Select objects to display:QC_Cells

ExportToSpreadsheet:[module_num:33|svn_version:'Unknown'|variable_revision_number:13|show_window:False|notes:[]|batch_state:array([], dtype=uint8)|enabled:True|wants_pause:False]
    Select the column delimiter:Comma (",")
    Add image metadata columns to your object data file?:Yes
    Add image file and folder names to your object data file?:Yes
    Select the measurements to export:No
    Calculate the per-image mean values for object measurements?:No
    Calculate the per-image median values for object measurements?:No
    Calculate the per-image standard deviation values for object measurements?:No
    Output file location:Elsewhere...|/Users/sorelle/Desktop/microscopy/scEBV_Lab_THUNDER/20250902_DMi8_HCM_for_paper/cellprofiler/cp_output_csv
    Create a GenePattern GCT file?:No
    Select source of sample row name:Metadata
    Select the image to use as the identifier:None
    Select the metadata to use as the identifier:None
    Export all measurement types?:Yes
    Press button to select measurements:
    Representation of Nan/Inf:NaN
    Add a prefix to file names?:Yes
    Filename prefix:A_DMi8_HCM_
    Overwrite existing files without warning?:No
    Data to export:Do not use
    Combine these object measurements with those of the previous object?:No
    File name:DATA.csv
    Use the object name for the file name?:Yes

ExportToDatabase:[module_num:34|svn_version:'Unknown'|variable_revision_number:28|show_window:False|notes:[]|batch_state:array([], dtype=uint8)|enabled:False|wants_pause:True]
    Database type:SQLite
    Database name:DefaultDB
    Add a prefix to table names?:Yes
    Table prefix:Akata_Test_
    Output file location:Default Output Folder|
    Create a CellProfiler Analyst properties file?:Yes
    Database host:
    Username:
    Password:
    Name the SQLite database file:DefaultDB.db
    Calculate the per-image mean values of object measurements?:Yes
    Calculate the per-image median values of object measurements?:No
    Calculate the per-image standard deviation values of object measurements?:No
    Calculate the per-well mean values of object measurements?:No
    Calculate the per-well median values of object measurements?:No
    Calculate the per-well standard deviation values of object measurements?:No
    Export measurements for all objects to the database?:All
    Select the objects:
    Maximum # of characters in a column name:64
    Create one table per object, a single object table or a single object view?:Single object table
    Enter an image url prepend if you plan to access your files via http:
    Write image thumbnails directly to the database?:Yes
    Select the images for which you want to save thumbnails:RGB_image
    Auto-scale thumbnail pixel intensities?:Yes
    Select the plate type:None
    Select the plate metadata:None
    Select the well metadata:None
    Include information for all images, using default values?:Yes
    Properties image group count:1
    Properties group field count:1
    Properties filter field count:0
    Workspace measurement count:1
    Experiment name:Akata_Test_
    Which objects should be used for locations?:Watershed_Cells
    Enter a phenotype class table name if using the Classifier tool in CellProfiler Analyst:
    Export object relationships?:No
    Overwrite without warning?:Never
    Access CellProfiler Analyst images via URL?:No
    Select the classification type:Image
    Select an image to include:None
    Use the image name for the display?:Yes
    Image name:Channel1
    Channel color:red
    Do you want to add group fields?:No
    Enter the name of the group:
    Enter the per-image columns which define the group, separated by commas:ImageNumber, Image_Metadata_Plate, Image_Metadata_Well
    Do you want to add filter fields?:No
    Automatically create a filter for each plate?:No
    Create a CellProfiler Analyst workspace file?:No
    Select the measurement display tool:BoxPlot
    Type of measurement to plot on the X-axis:Image
    Enter the object name:None
    Select the X-axis measurement:None
    Select the X-axis index:ImageNumber
    Type of measurement to plot on the Y-axis:Image
    Enter the object name:None
    Select the Y-axis measurement:None
    Select the Y-axis index:ImageNumber

Tile:[module_num:35|svn_version:'Unknown'|variable_revision_number:1|show_window:False|notes:[]|batch_state:array([], dtype=uint8)|enabled:False|wants_pause:True]
    Select an input image:Composite_rescale
    Name the output image:Img_Tile
    Tile assembly method:Within cycles
    Final number of rows:1
    Final number of columns:4
    Image corner to begin tiling:top left
    Direction to begin tiling:row
    Use meander mode?:No
    Automatically calculate number of rows?:No
    Automatically calculate number of columns?:No
    Select an additional image to tile:Comp_denoised
    Select an additional image to tile:Aes_Mask
    Select an additional image to tile:SanityCheck

SaveImages:[module_num:36|svn_version:'Unknown'|variable_revision_number:16|show_window:False|notes:[]|batch_state:array([], dtype=uint8)|enabled:False|wants_pause:False]
    Select the type of image to save:Image
    Select the image to save:Img_Tile
    Select method for constructing file names:From image filename
    Select image name for file prefix:Bright
    Enter single file name:OrigBlue
    Number of digits:4
    Append a suffix to the image file name?:No
    Text to append to the image name:
    Saved file format:tiff
    Output file location:Elsewhere...|/Users/sorelle/Desktop/microscopy/scEBV_Lab_THUNDER/20250902_DMi8_HCM_for_paper/cellprofiler/cp_output_imgs
    Image bit depth:8-bit integer
    Overwrite existing files without warning?:No
    When to save:Every cycle
    Record the file and path information to the saved image?:No
    Create subfolders in the output folder?:No
    Base image folder:Elsewhere...|
    How to save the series:T (Time)
    Save with lossless compression?:Yes
